# Phylogenetic comparative approach reveals evolutionary conservatism, ancestral composition, and integration of vertebrate gut microbiota

**DOI:** 10.1101/2023.01.03.522549

**Authors:** Benoît Perez-Lamarque, Guilhem Sommeria-Klein, Loréna Duret, Hélène Morlon

## Abstract

How host-associated microbial communities evolve as their hosts diversify remains equivocal: How conserved is their composition? What was the composition of ancestral microbiota? Do microbial taxa covary in abundance over millions of years? Multivariate phylogenetic models of trait evolution are key to answering similar questions for complex host phenotypes, yet they are not directly applicable to relative abundances, which usually characterize microbiota. Here, we extend these models in this context, thereby providing a powerful approach for estimating phylosymbiosis (the extent to which closely related host species harbor similar microbiota), ancestral microbiota composition, and integration (evolutionary covariations in bacterial abundances). We apply our model to the gut microbiota of mammals and birds. We find significant phylosymbiosis that is not entirely explained by diet and geographic location, indicating that other evolutionary-conserved traits shape microbiota composition. We identify main shifts in microbiota composition during the evolution of the two groups and infer an ancestral mammalian microbiota consistent with an insectivorous diet. We also find remarkably consistent evolutionary covariations among bacterial orders in mammals and birds. Surprisingly, despite the substantial variability of present-day gut microbiota, some aspects of their composition are conserved over millions of years of host evolutionary history.

## Introduction

Host-associated microbial communities, referred to as the *microbiota*, often play central roles in the biology of the hosts and their interactions with the environment. As host clades diversify, the microbiota can furthermore play a key role in the adaptation of their hosts to different ecological conditions. This raises important questions on the evolution of the microbiota as hosts diversify. First, how much is microbiota composition conserved over host evolutionary timescales? While the microbiota can be quite labile within and between host species (Ley et al. 2008; David et al. 2014; Hacquard et al. 2015; Hird et al. 2015; Amato et al. 2019), more closely related host species often tend to have more similar microbiota, a pattern referred to as *phylosymbiosis* (Brooks et al. 2016; Lim and Bordenstein 2020). In animals, levels of phylosymbiosis appear to be heterogeneous across tissues (e.g. gut or skin microbiota) and lineages (Mazel et al. 2018; Lim and Bordenstein 2020; Song et al. 2020; Perez-Lamarque, Krehenwinkel, et al. 2022).

The presence of a phylogenetic signal in microbiota composition across hosts could potentially be used to reconstruct ancestral microbiota composition. Ancestral reconstructions could be particularly useful to detect events during host diversification associated with major shifts in microbiota composition or to verify hypotheses on ancestral diets. A phylogenetic signal in microbiota composition may also inform on potential long-term evolutionary covariations in abundances between microbial taxa. Positive or negative covariations may arise from direct interactions between microbial taxa, such as cross-feeding, trophic relationships, or competition (Faust et al. 2012; Foster et al. 2017; Kohl 2020), or from (anti)correlated responses to variations in the environment (*e*.*g*. similar or opposite responses to decreased pH). We refer to these covariations as *microbiota integration* by analogy with the often observed *phenotypic integration* between traits in complex phenotypes (Pigliucci 2003). Such covariations would indicate constraints in the evolution of microbiota composition.

Phylogenetic comparative methods offer a rich toolbox for quantifying phylogenetic signal, reconstructing ancestral states, and detecting integration in multidimensional phenotypes (Clavel et al. 2015). These methods rely on modeling the evolution of a set of phenotypic traits across evolutionarily related species through a multivariate stochastic process, such as the Brownian motion process, running along the species’ phylogenetic tree (Revell et al. 2008; Harmon 2017). The multivariate Brownian process models the gradual evolution of traits through the accumulation of stochastic changes drawn from a multivariate normal distribution with a variance-covariance matrix that reflects the magnitude of the changes for each trait (the variance terms) and the covariation in the changes between trait pairs (the covariance terms). This process is relevant to represent long-term variations in the abundances of the different microbial taxa that constitute the microbiota, as such variations are an emerging outcome of: (i) the stochastic accumulation of changes in the numerous host traits that can influence the microbiota, including both extrinsic (e.g. geographic location, habitat) and intrinsic (e.g. diet, antimicrobial excretions) traits (Moran et al. 2019; Kohl 2020; Lim and Bordenstein 2020) and (ii) interactions between microbial taxa (Foster et al. 2017). Indeed, the Brownian motion process has already been used to model variations in microbial abundances over host evolutionary time (Capunitan et al. 2020; Labrador et al. 2021). However, the process is not directly applicable to compositional data made of relative microbial abundances as it does not constrain its components to sum to 1, and absolute abundances are unfortunately typically not provided by mainstream metabarcoding technics used to characterize microbiota composition. Thus, current phylogenetic comparative methods cannot directly be used in the context of microbiota evolution without transgressing several model assumptions (Hird 2019).

Here, we develop an approach to apply the multivariate Brownian motion process to compositional data. We also include a widely-used tree transformation (Pagel 1999) that quantifies phylosymbiosis by evaluating how much host phylogeny contributes to explaining interspecific variation in present-day microbiota composition. Phylosymbiosis is typically assessed using correlative approaches such as Mantel tests (Lim and Bordenstein 2020), which are known to suffer from frequent false negatives, while process-based approaches such as ours tend to be more powerful (Harmon and Glor 2010; Hird 2019; Perez-Lamarque, Maliet, et al. 2022). We apply our new approach to the gut bacterial microbiota of mammals and birds. The gut microbiota is key to the functioning of their hosts, contributing to their nutrition, their protection, and their development (McFall-Ngai et al. 2013). Strong phylosymbiosis in gut bacterial microbiota has been reported for mammals, including primates and rodents (Ochman et al. 2010; Groussin et al. 2017; Kohl et al. 2018), while it is thought to be absent for birds, with some exceptions in a few young clades (Song et al. 2020; Trevelline et al. 2020; Bodawatta et al. 2022). We revisit this dichotomy here, on the premise that previous analyses may have not been powerful enough to detect phylosymbiosis in birds (Hird 2019). We analyze potential drivers of phylosymbiotic patterns, including diet, geographic location, and flying ability, we estimate the ancestral microbiota composition of mammals and birds, and we investigate patterns of microbiota integration.

## Results & Discussion

We developed a method to infer the dynamics of microbiota composition during host diversification from host-microbiota data (*i*.*e*. a fixed, bifurcating host phylogeny and microbiota relative abundances for each extant host species) using the multivariate Brownian motion process (Figure 1 and Methods). We assume that all microbial taxa are present in all hosts, potentially in very low (undetectable) abundances and that they were already present in the most recent common ancestor of all host species. These assumptions are met if we consider a taxonomic level in the definition of microbial taxa that is high enough given the host clade, such as bacterial orders in the vertebrate gut microbiota. We assume that, from ancestral values at the root *X*_0_, the log-absolute abundances of the different microbial taxa change on the host phylogeny following a multivariate Brownian motion model with variance-covariance matrix *R* (Figure 1a). Under this model, the log-absolute abundances fluctuate around their ancestral values (log*X*_0_) without directional change. In addition, we account for variation linked to present-day factors by including in the model the widely-used Pagel’s λ transformation of the host phylogenetic tree (Pagel 1999). This transformation extends the terminal branches of the tree by (1-λ) of the total tree depth while compressing the internal branches to keep the total tree depth constant, with λ ranging between 0 and 1 (see Figure 1b and Methods). λ estimates close to 1 indicate that an untransformed tree explains the data quite well, reflecting strong phylosymbiosis, whereas λ estimates close to 0 indicate that the tree has little explanatory power, reflecting weak or absent phylosymbiosis. Unlike the traditional case of the multivariate Brownian motion process applied to phenotypic data, where the phenotype is directly measured at present, in the case of the microbiota, relative rather than absolute abundances are measured. To handle this difficulty, we treat total microbial abundances in each host as latent variables, and sample from the joint posterior distribution of these latent variables and our parameters of interest: Pagel’s λ, which provides us with an estimate of phylosymbiosis, the *R* matrix which reflects microbiota integration, and *Z*_0_, which indicates the relative microbial abundances in the ancestral microbiota.

**Figure 1:**
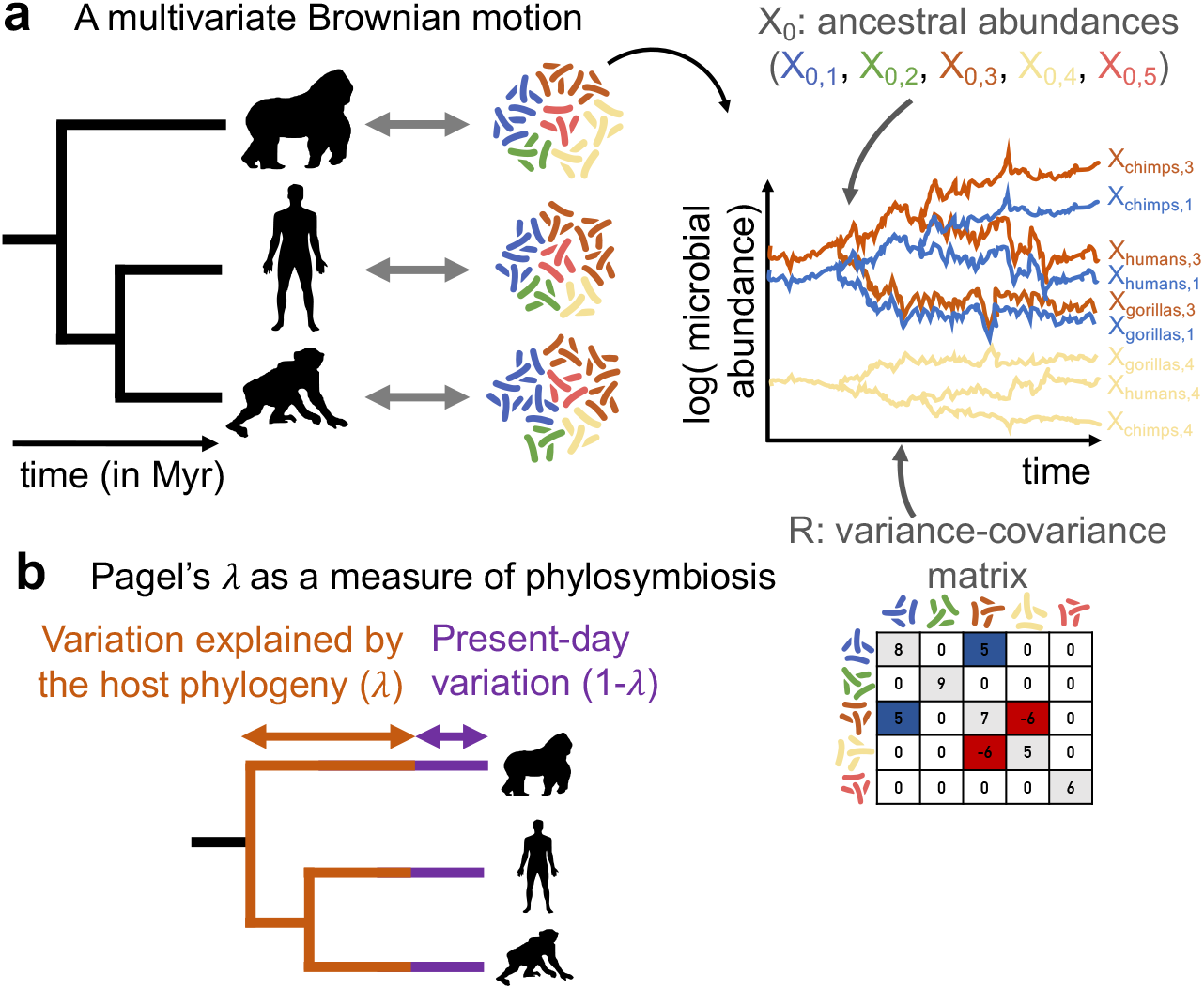
A comparative phylogenetic model for the dynamics of microbiota composition during host diversification: **(a)** We model fluctuations in the abundances of microbial taxa along a host phylogeny with a multivariate Brownian motion parametrized by the ancestral abundances (*X*_0_) and the variance-covariance matrix (*R*). The variance terms (on the diagonal) reflect the magnitude of the changes, while the covariance terms reflect positive or negative covariations in abundances between pairs of microbial taxa. The relative ancestral abundances (*Z*_0_) and the variance-covariance matrix *R* are estimated by adjusting the model to the host-microbiota data (host phylogeny and microbiota relative abundances for each host). **(b)** Following the widely-used Pagel’s λ transformation, we extend the terminal branches of the host phylogenetic tree by 1-λ of the total tree depth while compressing the internal branches to keep the total tree depth constant. λ is comprised between 0 and 1 and is co-estimated during inference. λ close to 1 indicates that closely related hosts tend to have similar microbiota due to shared evolutionary history (strong phylosymbiosis), while λ close to 0 indicates that microbiota composition is determined by present-day processes with little influence of host evolutionary history (weak or absent phylosymbiosis). The significance of phylosymbiosis is assessed with permutations.

We tested this inference method on data simulated from our model and found that we can accurately estimate the ancestral bacterial relative abundances *Z*_0_ (with a tendency for homogenization) and the variance-covariance matrix *R* between microbial taxa, provided that the number of host species (*n*) and bacterial taxa (*p*) are large enough (*n* ≥ 50 and *p* ≥ 5, see Supplementary Results 1). Similarly, the level of phylosymbiosis *λ* is accurately estimated for *p* ≥ 5, and its significance is correctly inferred for *n* ≥ 50 (see Supplementary Results 1). This approach provides a more powerful way to detect phylosymbiosis than Mantel tests, which often failed at detecting low levels of phylosymbiosis (0<λ<0.5; Table S1). This was expected, as Mantel tests are correlative and are known to suffer from frequent false negatives in comparison with more process-based approaches such as ours (Harmon and Glor 2010; Hird 2019; Perez-Lamarque, Maliet, et al. 2022).

We applied our model to the gut bacterial microbiota of 215 mammal species and 323 bird species from (Song et al. 2020) and found a pervasive signal of phylosymbiosis. We focused on the 14 most abundant bacterial orders, corresponding in abundance to 84% and 82% of the total gut bacterial microbiota of mammals and birds, respectively. We found a markedly higher level of phylosymbiosis in mammals (λ ≃ 0.65) than in birds (λ ≃ 0.31; Table S2, Figure S1), consistent with previous literature and our finding that microbiota composition is more species-specific in mammals than in birds (Table S3). Indeed, bird microbiota is generally more sensitive to short-term environmental changes such as anthropogenic perturbations or parasite infections (Bodawatta et al. 2022). By explicitly modeling the non-phylogenetic component of microbiota composition using a Pagel’s λ transformation, we detected a low but significant level of phylosymbiosis in the gut microbiota of birds (Table S2), contrary to previous conclusions (Song et al. 2020; Bodawatta et al. 2022) that relied on Mantel tests. *λ* values are higher at the level of bacterial phyla (Table S2; Figure S1), suggesting that microbiota composition is more evolutionarily conserved at higher taxonomic levels. Testing model performance on data simulated directly on the mammal and bird phylogenetic trees, we found a low type-I error rate and a high statistical power, suggesting that the phylosymbiosis we detected in birds is not due to false detection by our method, but rather to a higher power than previously used methods (Table S4). Phylosymbiosis is not linked to an effect of captivity nor the spurious concatenation of different studies either (Supplementary Results 2). Phylosymbiosis is particularly strong in Primates, Passeriformes, and Cetartiodactyla, lower but significant in Columbiformes, Chiroptera, and Carnivora, and non-significant in Rodentia, Charadriiformes, and Anseriformes (Table S2). Non-significant phylosymbiosis in these orders is likely due to an insufficient number of sampled species (*n* < 25, see Supplementary Results 1). It appears that vertebrate orders with mainly herbivorous diets have stronger phylosymbiosis, although this would need to be tested more robustly with a better species coverage (Table S2).

Our results suggest that phylosymbiosis is only partially explained by evolutionary conservatism in flying ability, diet, or geographic location. First, excluding flying mammals (Chiroptera) or non-flying birds did not impact our estimates of phylosymbiosis (Table S2). Second, permutation tests shuffling the microbiota of host species having the same diet, geographic location, flying ability, or combination of these traits resulted in much lower λ values (Figures 2 & S2). In mammals, λ values resulting from such shuffling are still significant (Figure 2), suggesting that the evolutionary conservatism of flying ability, diet, and geographic location contributes to phylosymbiosis without fully explaining it (Moran et al. 2019). In birds, shuffling often resulted in non-significant λ values (Figure 2), indicating a weak or absent contribution of diet or geographic location in the observed phylosymbiosis. Similarly, the conservatism of these traits is not sufficient to explain the phylosymbiosis measured in some of the larger mammal and bird clades, such as Primates, Cetartiodactyla, and Passeriformes (Figure S3). Thus, we suspect that other evolutionary-conserved physiological, immunological, or ecological traits act as host filters (Foster et al. 2017; Moran et al. 2019) and contribute to phylosymbiosis in the gut microbiota of mammals and birds (Goodrich et al. 2016; Mazel et al. 2018).

**Figure 2:**
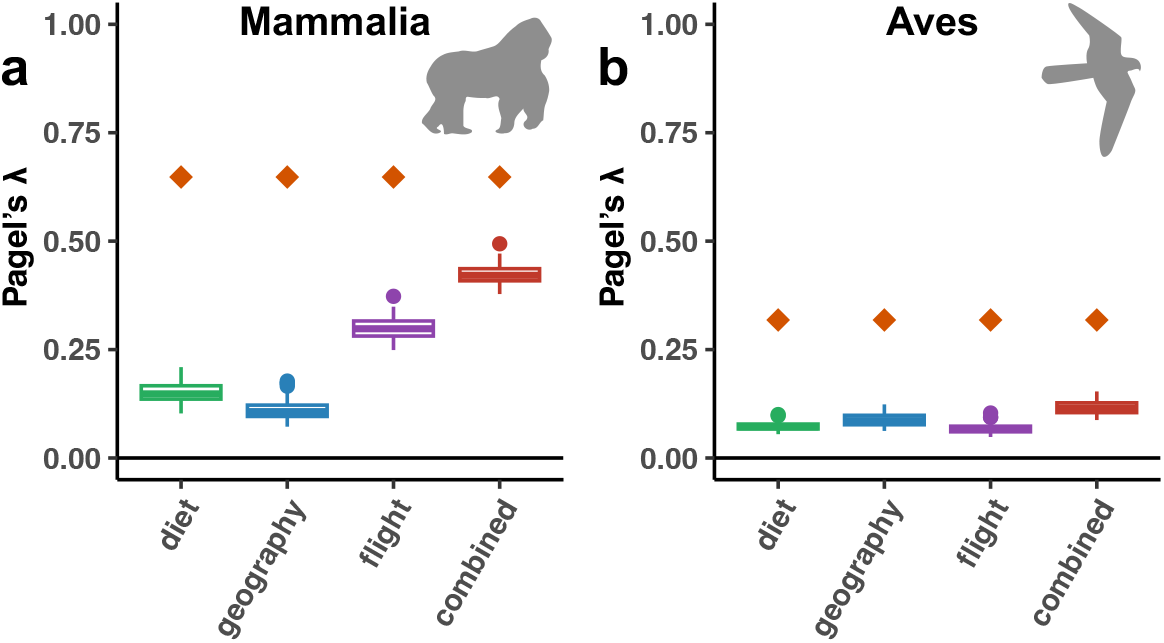
Phylogenetically-conserved diets, geographic locations, or flying abilities partially contribute to phylosymbiosis in the gut microbiota of mammals, but not birds. For both mammals and birds, we compared the estimated level of phylosymbiosis (mean λ value in orange) to levels of phylosymbiosis (λ values) estimated when shuffling the species that have the same diet (green boxplot), geographic location (blue boxplot), flying ability (flying or non-flying; purple boxplot), or combination of the latter traits (in red). For each shuffling strategy, we performed 100 randomizations. Combining all traits strongly constrains the possible permutations, which may consequently retain a phylogenetic signal in the shuffling and lead to high λ values although the traits are actually not strongly contributing to phylosymbiosis.

Our ancestral reconstructions of the microbiota of early mammals and birds suggest that Proteobacteria and Firmicutes were much more abundant in the ancestral gut microbiota of birds than mammals (Figures 3, 4, S4 & S5). As common in phylogenetic ancestral reconstruction, the uncertainty is quite high (Figure S6); it is larger in mammals than in birds because of the long branches that separate marsupials and eutherians at the origin of all mammals. In the absence of fossil constraints, ancestral reconstructions are a phylogenetically-weighted average of extant characteristics. Estimated ancestral compositions are thus expectedly close from the average microbiota compositions of extant bird and mammal species, yet they are distinct (Figure S7). Comparing the ancestral microbiota composition of mammals to that of the extant wild mammal species, we found the highest similarity with invertebrate feeders (distance to the centroid: *d*=1.46), such as the insectivorous armadillos (*Zaedyus pichiy*), and frugivores (*d*=1.24; Figures 5 & S8; see Methods), and the lowest similarity with specialist consumers feeding on plants (*d*=2.50) or meat (*d*=2.82). This result is robust to uncertainty in our estimate of ancestral microbiota composition (Figure S6b) and when including species sampled in captivity (Figure S7). Given that mammals originated before fleshy fruit plants (Eriksson 2016), this suggests that ancestral mammals were generalist invertebrate feeders, which is consistent with the current hypothesis, based on the fossil record and ancestral diet reconstruction, of a generalist insectivorous diet in early mammals (Gill et al. 2014; Grossnickle et al. 2019). We found the gut microbiota composition of modern birds to be only weakly structured by diet compared to that of mammals, making the inferred ancestral microbiota composition of birds less informative in this respect (no strong clustering in the PCA plots; PermANOVA testing the effect of diet: R^2^∼0.03, p<0.001 in birds *versus* R^2^∼0.22, p<0.001 in mammals; Figures 5, S7 & S8; Table S5). In addition, the fact that, under the assumptions of our model, most extant microbiota compositions in both mammals and birds remain centered around the estimated ancestral microbiota composition suggests that only a minority of the extant species experienced major shifts in their microbiota composition during their evolution.

**Figure 3:**
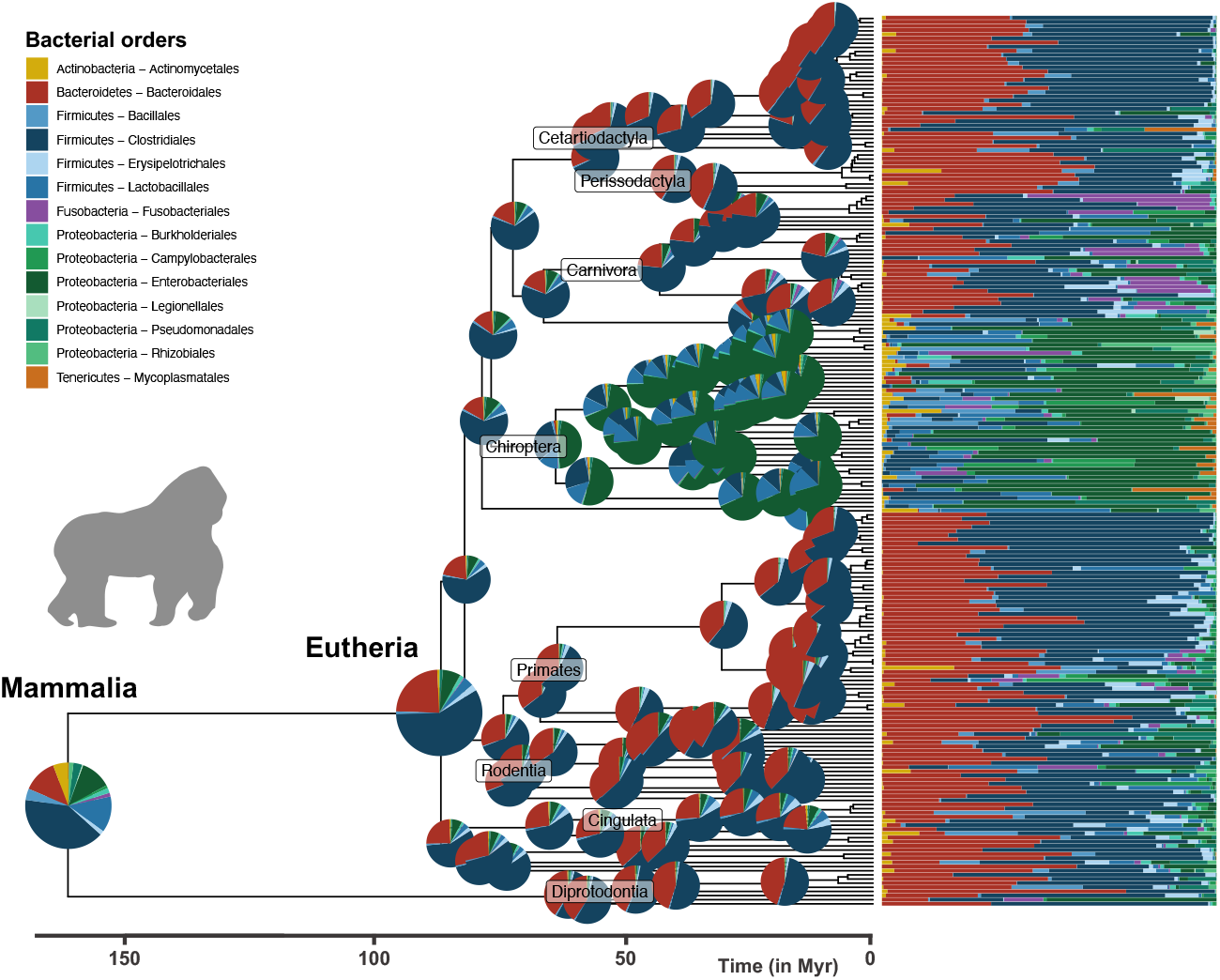
Ancestral reconstruction of mammalian gut microbiota: Phylogenetic tree of the sampled mammal species and associated relative abundances of the 14 most abundant bacterial orders (bar charts on the right). Pie charts at the root and nodes of the tree represent estimated ancestral microbiota compositions (mean of the posterior distribution of *Z*_0_ at the root and generalized least squares estimates at other internal nodes). Compositions are not represented at the most recent nodes for the sake of clarity.

**Figure 4:**
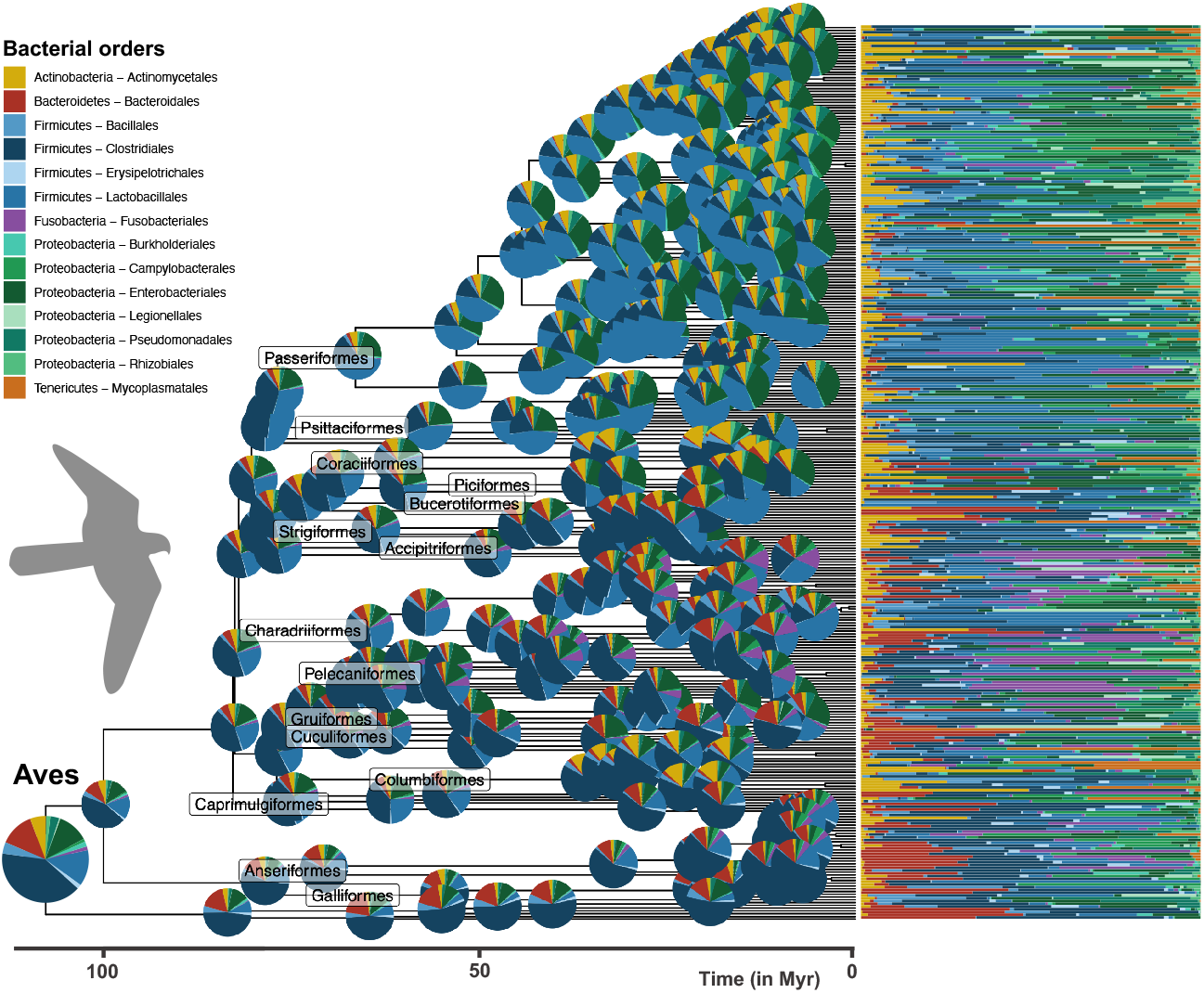
Ancestral reconstruction of avian gut microbiota: Phylogenetic tree of the sampled birds, and associated relative abundances of the 14 most abundant bacterial orders (bar charts on the right). Pie charts at the root and nodes of the tree represent estimated ancestral microbiota compositions (mean of the posterior distribution of *Z*_0_ at the root and generalized least squares estimates at other internal nodes). Compositions are not represented at the most recent nodes for the sake of clarity.

**Figure 5:**
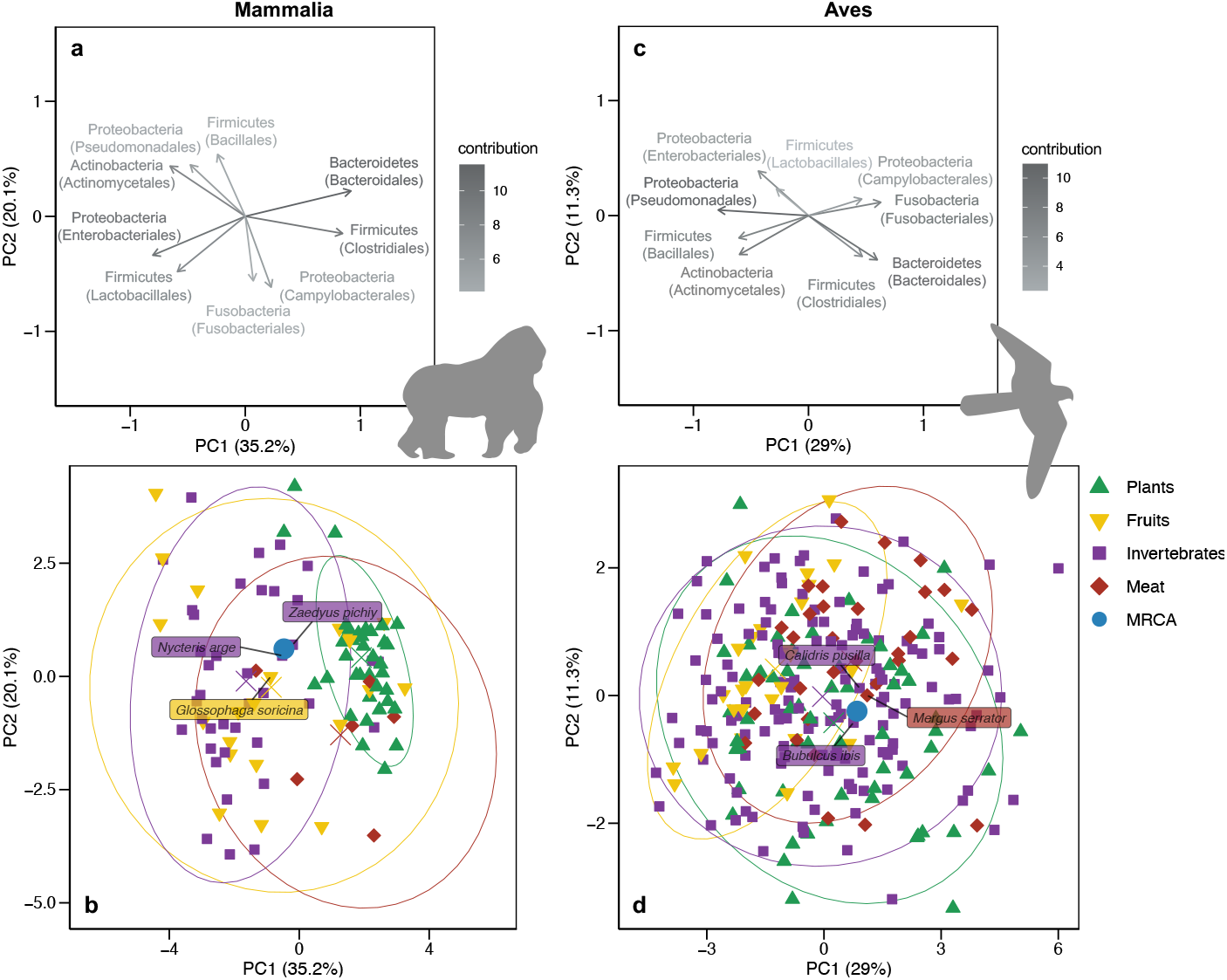
Projection of the estimated ancestral gut microbiota of mammals and birds onto the space of present-day gut microbiota. Top panels: projection of bacterial orders contributing to the two principal components (PC). Colors represent the contribution of the taxa to the principal components. Percentages indicate the explained variance of each PC. Only the 9 most abundant orders are represented for the sake of clarity. Bottom panels: Projection of the extant and ancestral microbiota compositions. Extant microbiota of species sampled in the wild are colored according to the species’ diet. For each diet, the ellipse contains on average 95% of the distribution approximated by a multivariate t-distribution and the centroid is indicated by a diagonal cross. Ancestral microbiota compositions of mammals and birds are represented in blue. On each PCA plot, we indicated the three extant species with microbiota compositions closest to the ancestral microbiota composition. The ancestral gut microbiota of mammals is closest to the gut microbiota of present-day invertebrate feeders; the gut microbiota of birds does not strongly reflect diet.

We detected significant changes in microbiota composition in the ancestors of some mammal and bird orders (Figures 3, 4, & S9). In mammals, the largest shift in microbiota composition occurred in the ancestor of Chiroptera, with an increased proportion of Enterobacteriales (Proteobacteria), Mycoplasmatales (Tenericutes), and to a lesser extent Actinomycetales (Actinobacteria), as well as a decreased proportion of Bacteroidales (Bacteroidetes), and in Firmicutes, Clostridiales were replaced by Bacillales and Lactobacillales (Figure 3; Table S6). Other shifts occurred in the ancestor of Carnivora, with an increased proportion of Fusobacteriales (Fusobacteria), and in the ancestors of Primates and Cingulata, with an increased proportion of some Firmicutes orders (*e*.*g*. Erysipelotrichales; Figures 3 & S9). In addition, Proteobacteria (especially Enterobacteriales and Pseudomonadales) almost disappeared in the ancestral microbiota of Ungulata and Simiiformes (New and Old World monkeys; Table S6). In birds, we found a shift in microbiota composition in the ancestor of Passeriformes, with more Bacillales and Enterobacteriales, and to a lesser extent Pseudomonadales, and a quasi-disappearance of Bacteroidales (Figures 4 & S5; Table S6). The ancestors of Anseriformes and Charadriiforms were characterized by a larger proportion of Bacteroidales, as well as a large proportion of Fusobacteriales, often absent or present in low abundances in other bird gut microbiota. Finally, the relative abundance of Actinomycetales increased in Columbiformes (Table S6). We found similar estimates of ancestral gut microbiota composition when running separate inferences for the different mammal and bird orders (Figure S9). Some of these compositional shifts might be linked to the ecological changes that these lineages experienced, such as the acquisition of flight for bats or carnivorous diets for Carnivora and Charadriiforms (Nishida and Ochman 2018; Song et al. 2020).

Far from varying as uncorrelated units during the evolutionary history of mammals and birds, we found significant covariances between many microbial taxa, both positive and negative (Figure 6a), suggesting strong constraints in the evolution of the microbiota. These patterns of microbiota integration are strikingly similar in mammals and birds (Figure 6b), indicating that they are conserved over long evolutionary times. Our simulation analyses on the mammal and bird trees suggest that these results are not artefactual, since we recover significant covariances only when we include them in the simulations (Table S7, Supplementary Results 1). Similar covariances were obtained when performing separate inferences on the different mammal and bird orders (Figure S10), which both confirms our results and suggests that the model assumption of a constant variance-covariance matrix across the host phylogenetic tree is reasonable. Combined with the high bacterial variability in time, across individuals, and across host species at low taxonomic levels, these consistent patterns at the level of bacterial orders on large time scales suggest that there is a certain level of functional redundancy among bacteria taxa within orders in the vertebrate gut microbiota.

**Figure 6:**
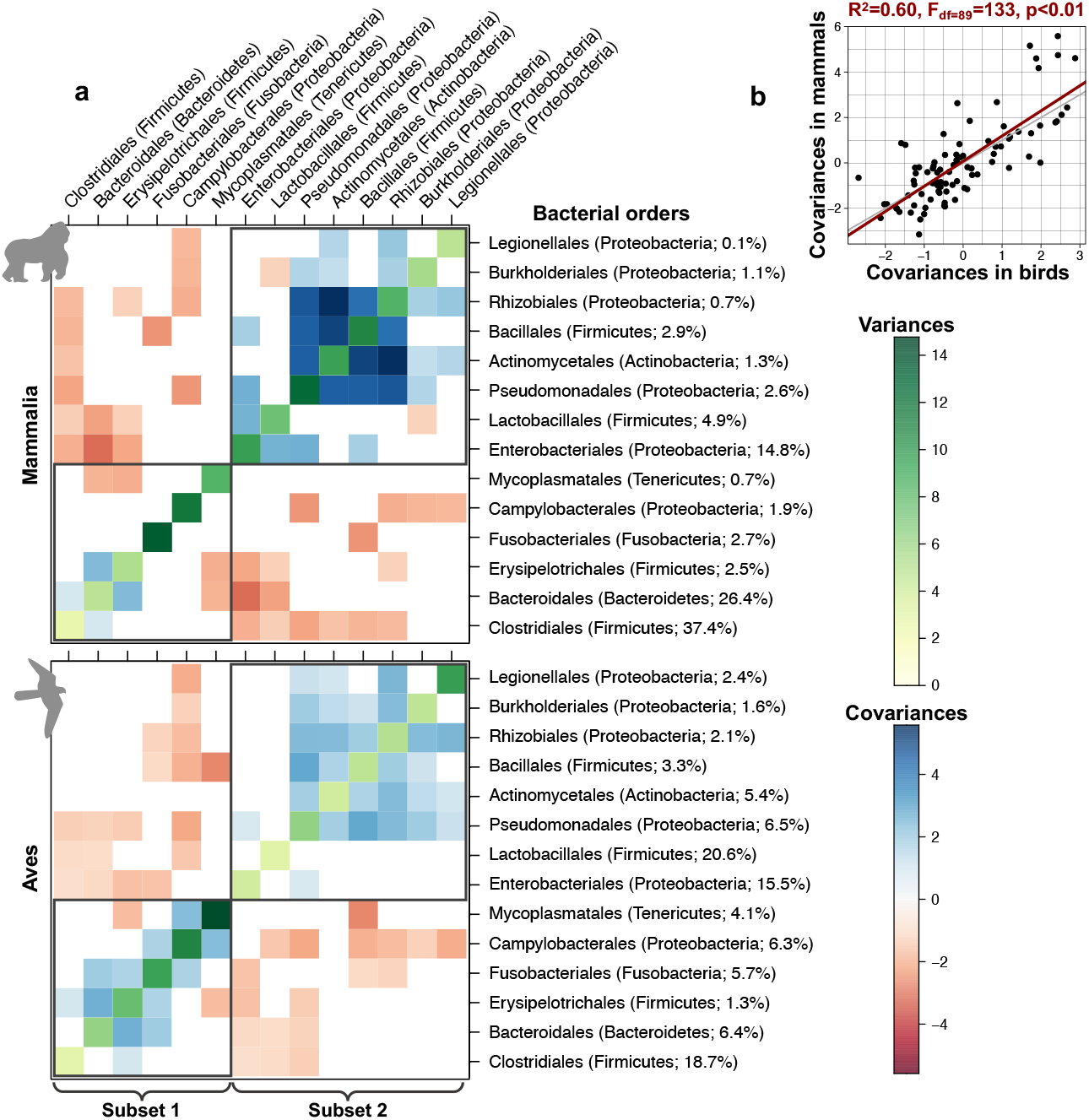
Estimated variances and covariances between the main bacterial taxa tend to be similar in the gut microbiota of mammals and birds. **(a)** For each variance-covariance matrix between bacterial taxa estimated using our model of host microbiota evolution, we represented negative covariances in red and positive covariances in blue, while variances are represented in shades of green. Non-significant covariances are represented in white. Grey rectangles correspond to subsets of bacterial orders that tend to covary positively. **(b)** Correlation between covariances between the main bacterial taxa estimated in the gut microbiota of mammals or birds. The red line indicates the corresponding linear model, while the grey line corresponds to y=x.

Both visual inspection and integration analyses of the covariances revealed that bacterial orders cluster into two main subsets within which taxa tend to covary in a concerted way, while taxa from different subsets tend to be anti-correlated (Figure 6; see Methods). The first subset (“subset 1”) is formed in particular by the orders Clostridiales, Bacteroidales, and Fusobacteriales, and the second subset (“subset 2”) is mainly composed of the orders Enterobacteriales, Lactobacillales, Pseudomonades, Actinomycetales, and Bacillales. Although some host species have a microbiota composed of an even mixture of these two bacterial subsets, one subset generally prevails, leading to the existence of two main gut microbiota profiles. The first subset is dominant in the microbiota of most mammals (excluding Chiroptera), the ancestors of birds, and some extant bird lineages (*e*.*g*. Anseriformes, Columbiformes, or Accipitriformes); the second subset predominates in the microbiota of Chiroptera and other bird lineages, including Passeriformes (Figure S11). This result suggests the existence of two main gut microbiota profiles conserved over millions of years across vertebrates.

We can only speculate on the processes underlying positive or negative covariances between bacterial orders: we cannot distinguish from our analyses whether they indicate direct interactions between bacterial taxa (*e*.*g*. cross-feeding or competition) or indirect interactions mediated by similar/opposed microbial responses to changes in the gut environment. For instance, the frequent and strong negative covariations observed between the abundant Enterobacteriales (Proteobacteria) and the major bacterial orders Clostridiales (Firmicutes) and Bacteroidales (Bacteroidetes) may result from direct competitions (Shealy et al. 2021), host immunological controls over Proteobacteria (Mirpuri et al. 2013), and/or be mediated by the oxygen concentration in the gut, as Proteobacteria are facultative anaerobes, while other phyla are obligate anaerobes (Shin et al. 2015). The strongest positive covariations we inferred between Actinomycetales, Pseudomonadales, and Rhizobiales, which are the most abundant bacterial orders in plant tissues (Wagner et al. 2016), may reflect a plant-based diet, which would lead to a concomitant increase of plant-associated bacteria in the gut microbiota of herbivorous vertebrates (Dion-Phénix et al. 2021). Some of the covariations we detected (e.g. the negative covariation between Lactobacillales and Bacteroidales) have also been observed in human microbiome data using co-occurrence network analyses (Faust et al. 2012), suggesting that at least some covariations between microbial taxa that occur over short timescales within host species are conserved over macroevolutionary timescales.

To test the adequacy of our model to the data, we simulated microbiota under our model using the parameters estimated on mammal and bird data. We found that simulated microbiota have compositions similar to those observed in extant mammals and birds (Figure S12), which indicates that, despite its simple assumptions, our multivariate Brownian motion model generates realistic gut microbiota (Hird 2019; Labrador et al. 2021). Nevertheless, the gut microbiota composition of mammals and birds appears more constrained than the sets of compositions we can simulate using multivariate Brownian motions (Figure S12). This is particularly true for mammals and may be linked to constraints that are not accounted for by our model, such as selective pressures toward particular microbiota compositions, the potential existence of carrying capacities for some bacterial orders, or non-constant or non-homogeneous variance-covariance matrices (*e*.*g*., more frequent shifts in microbiota composition early in clades history, effects of host traits such as diet or gut pH on covariation). Extensions of our multivariate Brownian motion approach could accommodate such constraints, but this may complexify inferences. We hope that this work will foster the development of more complex models that may better represent microbiota evolution in systems that present non-Brownian behaviors. As a first step, extensions that relax the constant variance assumption (*e*.*g*., the early-burst model; Harmon et al. 2010) would be relatively straightforward to implement and could be particularly relevant to account for the major shifts in microbiota composition that took place at the origin of some mammalian orders (*e*.*g*., in bats). Meanwhile, by relying on a simple and flexible Brownian motion process, our phylogenetic comparative model for microbiota evolution is general enough to be broadly applied across other host-microbiota systems and reveal the global trends of microbiota evolution.

Besides modeling assumptions, our results may be influenced by the inherent biases of metabarcoding data. Bacterial relative abundances characterized using metabarcoding techniques are a distortion of the actual relative abundances (Knight et al. 2018; Lavrinienko et al. 2021), since metabarcoding is sensitive to the number of rRNA copies in the bacterial genomes, primer biases, and the quality and completeness of the reference database for taxonomic assignation (at the bacterial order/phylum level in our case). These issues are unlikely to artefactually generate phylosymbiosis or covariations across bacterial taxa because we expect such biases to be homogeneous across host species; nevertheless, they are likely to affect our ancestral reconstructions of microbiota compositions.

Our approach to quantifying phylosymbiosis characterizes microbiota composition in terms of the relative abundances of higher bacterial taxa (orders or phyla). This characterization hides variations in the presence/absence of bacterial taxa at lower taxonomic levels (*e*.*g*. genus or species). Indeed, distinct mammal or bird species are known to host different bacterial species (Song et al. 2020), and this may not translate into abundance variations at higher taxonomic levels if the different bacterial species belong to the same higher taxa. Besides the widely-used Mantel tests, such variations could be accounted for by stochastic processes modeling the evolution of presence/absence on host phylogenies (Braga et al. 2020), although we are not aware that these approaches have been used to detect bacterial phylosymbiosis. Yet another level of variation in microbiota composition that can contribute to phylosymbiosis arises through genetic differentiation below the bacterial species level: if a bacterial species is vertically transmitted during host diversification, we expect bacterial strains from closely related host species to be more genetically similar (Sanders et al. 2014; Groussin et al. 2017; Perez-Lamarque and Morlon 2019). This latter process can be specifically tested thanks to cophylogenetic methods that consider the evolution of each microbial species separately (Dismukes et al. 2022; Perez-Lamarque and Morlon 2022). The above-mentioned methods are complementary, as they focus on different levels of variations in microbiota composition, and on the distinct processes that simultaneously generate phylosymbiosis (Moran et al. 2019; Lim and Bordenstein 2020).

Phylosymbiosis is a widespread pattern that has fascinated microbial ecologists and evolutionary biologists since its discovery, spurring debates on the main processes underlying the pattern. Drawing upon phylogenetic comparative methods, we have developed a new approach to studying phylosymbiosis. Our results on simulations and birds suggest that phylosymbiosis may be even more prevalent than currently recognized, but sometimes undetected with correlative approaches. We have shown that conservatisms in diet, geographic location, and flying ability are not enough to explain phylosymbiosis, calling for an investigation of the role of other host ecological traits, as well as physiological and immunological traits. One of our most striking results, in the face of the well-known high variability of the gut microbiota, is its high level of integration, with conserved covariations between bacterial orders over millions of years. The same two subsets of bacterial orders tend to covary in a concerted way in both mammals and birds, leading to the existence of two main gut microbiota profiles in vertebrates. Hence, microbial interactions combined with phylogenetically-conserved host traits shape microbiota composition over millions of years, supporting the view of vertebrate gut microbiota as ‘ecosystems on a leash’ (Foster et al. 2017).

## Methods

### A multivariate Brownian motion model for variations in microbiota composition over host evolutionary time

We denote by *p* the total number of microbial taxa detected across the microbiota of the *n* sampled host species. Standard metabarcoding techniques only measure the relative abundance of each microbial taxon *j* in each extant host species *i*, which we denote by *Z*_*ij*_ = *X*_*ij*_/*Y*_*i*_, where *X*_*ij*_ is the unmeasured absolute abundance of microbial taxon *j* in host *i* and *Y*_*i*_ = ∑_*j*_ *X*_*ij*_ is the unmeasured total microbial abundance in the microbiota of host *i*. We assume that the logarithms of microbial absolute abundances log*X*_*ij*_ vary along the host phylogenetic tree according to a multivariate Brownian motion starting from the ancestral abundances at the root, denoted by X_0*j*_ (Figure 1). Indeed, taking the logarithm of the abundances yields values on the real axis that are amenable to be modeled with a Brownian motion, similar to continuous phenotypic traits. This model implies a log-normal distribution of abundances, as is commonly observed in microbial communities (Quince et al. 2008), and it can easily accommodate undetected microbial taxa in some hosts by assigning them very low unobserved relative abundances. To make the model identifiable, we express the total abundances *Y*_*i*_ relative to the unknown total abundance at the root *Y*_0_, and we only infer 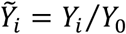 Each microbial taxon *i* is characterized by a certain variance and pairs of microbial taxa can affect each other through a covariance term, so that their changes in abundance over time can be positively or negatively correlated. All variance and covariance values are assumed to be constant along the host phylogeny and are summarized by the invertible variance-covariance matrix *R* (Figure 1a).

### Model inference

To infer the model parameters, we sampled from their joint posterior distribution 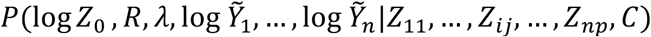 using a No U-turn Hamiltonian Monte Carlo sampler, a computationally efficient Markov Chain Monte Carlo algorithm for continuous variables (Supplementary Methods 1). We implemented it in the probabilistic programming language Stan and we ran and compiled it through the RStan interface (R Core Team 2022; Stan Development Team 2022). Inferences were performed with 4 independent chains and a minimum of 4,000 iterations per chain including a warmup of 2,000 iterations. We checked the convergence of the chains using the Gelman statistics and effective sample sizes (ESS). We extracted the mean posterior value of each parameter and its associated 95% credible interval across posterior samples.

We considered a covariance to be significant if 0 was not included in its 95% credible interval. We could not use the same approach for *λ*, because it only takes positive values. Furthermore, model selection using Bayes factors led to many false negatives on simulated data (Supplementary Methods 2 & Results 1). Therefore, we assessed the significance of *λ* using permutations. We shuffled at random the extant host species to break the phylogenetic structure and ran again model inference of the randomized dataset. We performed 100 replications and compared the distribution of *λ* values thus obtained to the original *λ* estimate: if the original *λ* was greater than at least 95% of the *λ* values obtained through permutations, we considered that there was a significant impact of host evolution on microbiota evolution.

### Simulations

We evaluated our approach using simulations. We simulated the evolution of a microbiota along a host phylogeny using Multivariate Brownian motions for log-abundances. We simulated phylogenies with *n* = 20, 50, 100, or 250 extant host species using a pure birth model (*pbtree* function in the phytools R-package (Revell 2012)). We considered microbiota with *p* = 3, 5, 10, or 15 microbial taxa and uniformly sampled the logarithms of their ancestral abundances at the root of the host phylogeny between-4 and 0 before normalizing them so that ∑_*j*_ *Z*_0*j*_ = 1. We generated random positive definite variance-covariance matrices *R* following (Uyeda et al. 2015) and (Clavel et al. 2019) with eigenvalues of 1/4. Finally, we applied Pagel’s *λ* transformations with *λ* = 1, 0.75, 0.5, 0.25, or 0. For each combination of *n, p*, and *λ* values, we performed 100 independent simulations, leading to a total of 8,000 simulations. We verified that our approach correctly estimates the parameters *λ, Z*_0_, and *R*, and detects phylosymbiosis (significant *λ*) and covariations (significant *R* components) when they are simulated. We compared the performances of our approach for detecting phylosymbiosis to that of Mantel tests (Perez-Lamarque, Maliet, et al. 2022).

We also evaluated our inference approach using data simulated on the phylogenetic tree of mammals or birds, and using conditions and parameters matching the empirical data. We performed simulations with 7 taxa (corresponding to the 7 bacterial phyla in the data, see below) and 14 taxa (corresponding to the 14 bacterial orders in the data). We used values of λ = 1, 0.75, 0.5, 0.25, or 0, and values for the other model parameters similar to those estimated from the empirical data (Figure S13). We performed 100 simulations per condition (thus reaching a total of 2,000 simulations).

### Empirical application

We downloaded the dataset of (Song et al. 2020) that gathered the gut microbiota of 2,677 mammal individuals from >200 species and 1,630 bird individuals from >300 species, characterized by metabarcoding using the V4 region of the 16S rRNA gene. Only studies using the standard protocol of the Earth Microbiome Project (Thompson et al. 2017) were included (see (Song et al. 2020) for details), making samples comparable across different studies (Knight et al. 2018). Song *et al*. converted bacterial reads into amplicon sequence variants (ASV), assigned each ASV taxonomically using the Greengenes database (DeSantis et al. 2006; Song et al. 2020), and rarefied ASV tables at 10,000 reads per sample. We complemented their dataset with the consensus phylogenetic trees of (Upham et al. 2019) and (Jetz et al. 2012) for mammals and birds, respectively. We only kept the species having their microbiota compositions characterized by at least 2 microbiota samples (Table S2). We checked that gut microbiota from the same host species were more similar than gut microbiota from different species using PermANOVA (Oksanen et al. 2016). Then, we obtained the microbiota composition of each host species by averaging the samples per species and extracted the relative abundances of the main bacterial orders and phyla per host species. We verified that similar results were obtained when repeating our analyses by randomly sampling one individual per host species (Figure S14). We only considered the 14 most abundant bacterial orders, *i*.*e*. those that each represented more than 1% of the total bacterial abundance (which correspond in abundance to 84% and 82% of the total gut bacterial microbiota of mammals and birds, respectively) and the 7 most abundant bacterial phyla (95% and 96% of the gut microbiota of mammals and birds respectively; Figure S15). We also repeated all analyses using only the 9 (resp. 5) most abundant orders (resp. phyla). We did not apply our model at lower taxonomic levels mainly because the assumptions of our model (all microbial taxa are present in all hosts, potentially in undetectable abundances and they were already present in the most recent common ancestor of all host species) are more likely to be met at high taxonomic levels. At lower taxonomic levels, the microbiota evolution of mammals and birds may be better represented using models of colonization and extinction (Song et al. 2020) than models of fluctuations in bacterial abundances such as ours. In addition, running the model with several hundreds of taxa would be computationally intensive. Finally, the quality of the taxonomic assignation and the number of taxa representing more than 1% of the gut microbiota decreased sharply at low taxonomic levels: only 81% and 45% of the gut microbiota of mammals and birds are assigned at the family and genus levels, respectively, and among them, only 60% and 18% of the bacterial taxa represent more than 1% of the gut microbiota.

Our multivariate Brownian motion model of microbiota does not explicitly consider losses of bacterial taxa from the microbiota through time. Yet, some bacterial taxa can be absent or undetected in the gut microbiota of mammals and birds. We assumed that the absence of a particular taxon came from a very low abundance, below the detection threshold: we thus arbitrarily set the relative abundances of absent taxa to 0.001%. Setting the minimal relative abundances of absent taxa to 0.01% reduced the estimated variance of the rare taxa but did not affect other estimates (Figure S16).

We applied the model separately on all mammals and all birds, getting estimates of Pagel’s *λ*, the ancestral microbiota composition *Z*_0_, and the variance-covariance matrix *R* for each vertebrate class.

### Effect of host traits on phylosymbiosis

We gathered data on host species traits from (Song et al. 2020) for diet, geographic location, and flying ability. We assigned a dominant diet to each host species as either “plants”, “fruits”, “invertebrates” or “meat” following the EltonTraits database (Wilman et al. 2014). We assigned a geographic location to each species by picking the biogeographic realm (Afrotropical, Antarctic, Australasian, Nearctic, Neotropic, Oriental, or Palearctic) where the highest number of wild individuals were sampled, or if not available, where the highest number of captive individuals were sampled (this was the case for 48% of the mammalian species and 18% of the avian ones). We treated flying ability as binary (yes/no). First, we assessed the influence of flight on the gut microbiota by performing inferences on non-flying mammal species only (*i*.*e*. excluding bats) and on flying bird species only. Similarly, we investigated the effect of captivity on our inferences by replicating them using only the gut microbiota of wild or captive individuals. Second, we tested whether the evolutionary conservatism of diet, geographic location, or flying ability may explain phylosymbiosis in mammals and birds by performing permutations. We shuffled host species having the same diet, geographic location, and/or flying ability and re-ran the inferences on these randomized datasets. For each tested trait, we performed 100 independent randomizations. Finally, we verified that phylosymbiosis did not artefactually arise from the concatenation of the separate studies composing this dataset by randomizing the species that came from the same study.

### Comparison between ancestral and present-day microbiota composition

We compared the estimated ancestral microbiota composition *Z*_0_ of all mammals or birds to that of extant species using principal component analysis (PCA) after applying a centered log-ratio transform to the abundances (Aitchison 1983). Given *Z*_0_, we also jointly estimated the ancestral abundances at each node of the host phylogenetic tree using generalized least squares following (Martins and Hansen 1997; Cunningham et al. 1998; Clavel et al. 2019). As a first attempt to infer past diet based on the estimated ancestral microbiota composition *Z*_0_, we computed the centroid of each of the four diet categories and computed the distance *d*_*i*_ between *Z*_0_ and each centroid on the first five PC axes. We additionally performed separate model inference for all orders of mammals (Carnivora, Cetartiodactyla, Chiroptera, Primates, and Rodentia; Table S2) and birds (Anseriformes, Charadriiformes, Columbiformes, and Passeriformes) represented by at least 15 species, and compared the ancestral microbiota composition obtained with separate and joint inferences.

### Integration analyses

We identified the significantly positive or negative covariances between bacterial orders. In addition, to characterize potential subsets of bacterial taxa that tend to vary in a concerted way, we clustered taxa using the *cluster_fast_greedy* function in the R-package igraph (Csardi and Nepusz 2006), based on the estimated variance-covariance matrix *R*, modified to retain information of only positive covariances (negative ones were set to 0).

### Model adequacy

To assess whether our model for the evolution of the gut microbiota of mammals and birds yields realistic microbiota compositions, we simulated the process of microbiota evolution on the mammal or bird phylogenies using the parameters estimated for mammals and birds (log*Z*_0_, *R*, and λ). Next, we compared the simulated microbiota compositions to the empirical microbiota compositions of the extant mammal or bird species using principal component analysis (PCA). We performed 20 independent simulations for each of our model inferences.

### Data availability

Raw data and processed data from (Song et al. 2020) used to perform the empirical applications are available in Qiita (https://qiita.ucsd.edu/study/description/11166).

Our phylogenetic comparative method, referred to as ABDOMEN (A Brownian moDel Of Microbiota EvolutioN), is available on GitHub with a tutorial: https://github.com/BPerezLamarque/ABDOMEN.

## Acknowledgments

The authors acknowledge Julien Clavel, Jonathan Drury, Félix Foutel--Rodier, and members of the BioDiv team at IBENS for helpful discussions, as well as the Editor Aurélien Tellier and two anonymous reviewers for their constructive comments. They also acknowledge the Hubert Curien Alliance program for funding workshops that initiated the project. GSK acknowledges support from the Academy of Finland (decision 340314) and the Sakari Alhopuro foundation (grant 20210172). This work was performed using HPC resources from GENCI-IDRIS (Grants 2021-A0100312405 and 2022-AD010313735).

## Author Contributions

BPL, GSK, LD, and HM designed the study. BPL and GSK implemented the model. BPL performed the simulations and the empirical applications. BPL, GSK, and HM wrote the manuscript.

