## Supplementary for "Phylogenetic comparative approach reveals evolutionary conservatism, ancestral composition, and integration of vertebrate gut microbiota"

**This PDF contains:**

**Supplementary Methods (S1-S2)**

**Supplementary Results (S1-S2)**

**Supplementary Tables (S1-S9)**

**Supplementary Figures (S1-S24)**
Empirical application
Simulations
Simulations on empirical phylogenies

#### Supplementary Methods:

##### Supplementary Methods 1: Likelihood of the model and inference

Under a multivariate Brownian motion, the logarithm of the abundance of microbial taxon *j* in host *i,* denoted by $\log X_{ij}$, is expected to vary around its ancestral abundance at the tree root $\log X_{0i}$ according to the variance-covariance matrix $R$ between microbial taxa. The diagonal elements of $R$ reflect the magnitude of changes in log-abundance overt time and its off-diagonal elements the positive or negative covariations in log-abundance between pairs of microbial taxa.

We denote by $C$ the phylogenetic variance-covariance matrix that gives the total shared branch length from the root between any pair of host species. Its elements range from 0, for two host species without any shared branch length, to 1 for the diagonal elements. $C$ describes how similar the microbiota composition of a pair of host species is expected to be due to their phylogenetic relatedness under a Brownian motion process. We then introduce the matrix $C_\lambda$ obtained by multiplying all off-diagonal elements of $C$ by a factor λ between 0 and 1 and retaining 1s on the diagonal. This so-called "Pagel's λ transformation" can be interpreted as follows: for $\lambda = 0$, $C_\lambda$ corresponds to the identity matrix, hence all covariances between host species are equal to 0 and there is no more phylogenetic structure, whereas for $\lambda = 1$, $C_\lambda$ is equal to the original phylogenetic covariance structure $C$ resulting from a Brownian motion process running along the tree. Inferring $\lambda$ from data allows estimating the strength of phylosymbiosis.

Under this multivariate Brownian motion model with Pagel's $\lambda$ transformation, the joint distribution of all microbial log-abundances across all host species follows a multivariate normal distribution with a variance-covariance matrix given by the Kronecker product $R \otimes C_\lambda$. The log-likelihood $\mathcal{L}_X = \log[P(\log X \,|X_0, R)]$ of this process can be expressed as (Clavel *et al.,* 2019):

$$\mathcal{L}_X = -\frac{1}{2}\big(np\log(2\pi) + p\log|C_\lambda\,| + n\,\log|R| \\ + \mathrm{tr}\left[R^{-1}(\log X - \log X_0)^t {C_\lambda}^{-1}(\log X - \log X_0)\right]\big)$$

However, we are interested here in the distribution of the relative microbial abundances at present $Z_{ij} = X_{ij}/Y_i$, where $Y_i = \sum_j X_{ij}$, rather than that of the $X_{ij}$ since the latter are unmeasured. Obtaining this distribution requires marginalizing over the $Y_i$, which we could not do analytically and is very computationally intensive to do numerically.

Instead, we sought to perform Markov Chain Monte Carlo inference of the joint posterior distribution $P(\log Z_0\,, R, \lambda, \log Y_i, \dots, \log Y_n | Z_{11}, \dots, Z_{ij}, \dots, Z_{np}, C)$. Nevertheless, to make the model identifiable we expressed the total abundances $Y_i$ relative to their unknown value at the root $Y_0$ and we only inferred $\tilde{Y}_i = Y_i/Y_0$. We used a No U-turn

Hamiltonian Monte Carlo sampler, a computationally efficient Markov Chain Monte Carlo algorithm for continuous variables, implemented in the probabilistic programming language Stan. We used the following prior distributions:

$$\lambda \sim Unif(0,1)$$
$$Z_0 \sim Dirichlet(\mathbf{1}_p)$$
$$\log \tilde{Y}_i \sim \mathcal{N}(\mu_{\tilde{Y}}, C_\lambda \sigma_{\tilde{Y}})$$
$$R \sim \mathcal{W}^{-1}(p, \mathbf{1}_p)$$

where $\mathcal{W}^{-1}$ denotes the Inverse-Whishart distribution. We applied a non-centered parametrization with Cholesky decomposition to the $\log \tilde{Y}_i$, which sped up the inference considerably, and we set $\mu_{\tilde{Y}} = 0$ and $\sigma_{\tilde{Y}} = 2$.

Finally, we can remark that since the absolute abundances $X_{ij}$ follow a multivariate log-normal distribution, the total absolute abundance $Y_i$ in host *i* is the sum of correlated log-normal random variables. Lo (2013) provide an approximation of the distribution of $Y_i$ as a lognormal distribution with parameters $\mu_{Y_i}$ and $\sigma^2_{Y_i}$ given by:

$$\mu_{Y_i} = \log(\mathbb{E}[Y_i]) - \sigma^2_{Y_i}/2$$
$$\sigma^2_{Y_i} = \frac{1}{\mathbb{E}[Y_i]^2} + \sum_{k,l \in [1,p]} \mathrm{R}_{kl}(\mathrm{R}_{kk}\mathrm{R}_{ll})^{-1/2}\mathrm{E}[X_{ik}]\mathrm{E}[X_{il}]$$
$$= \frac{1}{\mathbb{E}[Y_i]^2} + \sum_{k,l \in [1,p]} \mathrm{R}_{kl}(\mathrm{R}_{kk}\mathrm{R}_{ll})^{-1/2} X_{0k} e^{R_{kk}/2} X_{0l} e^{R_{ll}/2}$$

where $\mathbb{E}[Y_i]$ can be computed as:

$$\mathbb{E}[Y_i] = \mathbb{E}\left[\sum_{k=1}^{p} X_{ik}\right] = \sum_{k=1}^{p} X_{0k} e^{R_{kk}/2}$$

Thus, depending on the values of the variances and covariances, our model predicts either an average increase, $Y_i > Y_0$, or an average decrease, $Y_i < Y_0$, of the total bacterial abundances over time, where $Y_0$ is the total absolute abundance at the root of the tree.

**Supplementary Methods 2:**

We tested to evaluate the significance of $\lambda$ using Bayes factors. To do so, we used the *bridgesampling* R-package (Gronau et al. 2020) to estimate the marginal likelihood of the model and compared it to the marginal likelihood of a null model with $\lambda = 0$, *i.e.* where the abundances of the microbial taxa vary in each host lineages independently of the host phylogeny. We computed Bayes factors *BF* as the ratio of these marginal likelihoods and considered $BF > 1$ as a significant support for $\lambda > 0$.

#### Supplementary Results:

##### Supplementary Results 1:

Simulations revelated that our model can correctly estimate the ancestral relative bacterial abundances $Z_0$, especially when the number of host species $n$ and the number of bacterial taxa $p$ are large (Figure S17). When $n$ and $p$ are low (*e.g.* $p < 5$ and $n < 50$), low ancestral abundances tend to be overestimated and high ancestral bacterial abundances tend to be underestimated (Figure S17), *i.e.* the ancestral abundances tend to be evened out.

For covariances, >70% of the large positive and negative simulated covariances were correctly inferred for large $n$ and $p$ ($n = 250$ and $p = 15$, Table S8; Figure S18). The ability to infer significant covariances decreased with lower $n$ and $p$. In particular, we noticed that positive covariances were often inferred as non-significant in these conditions (low statistical power). Negative covariances were very rarely inferred as positive covariances (<2%), indicating that false positives are rare. Similarly, positive covariances were rarely inferred as negative covariances (<5%), except in the extreme case where $p \leq 5$, in which case false positive rates can increase above 40% (Table S8). For small positive and negative simulated covariances, the majority (~70%) were inferred as non-significant (low statistical power; Table S8). In addition, >50% of the small positive covariances were actually inferred as significantly negative (high false positive rate) when $p = 3$ and $n = 250$. Therefore, inferred covariances should only be interpreted when the number of bacterial taxa ($p$) is large enough, *i.e.* at least 5.

Estimations of the level of phylosymbiosis ($\lambda$) were correct overall (Figure S19), except when $p = 3$ (overestimation of $\lambda$) or when $p$ was large compared to $n$ (underestimation of $\lambda$). The evaluation of the significance of $\lambda$ based on Bayes factors led to contrasting results (Table S9): False positives were always very low (<1%), but the statistical power of this approach was also quite low in some cases. Indeed, when $n, p,$ and simulated $\lambda$ are low, we always infer $\lambda$ as non-significant. We only reached a satisfying statistical power (>50%) for $n = 250$ and $p = 15$. Bayes factors should therefore only be used for large $n$ and $p$. In contrast, the evaluation of the significance of $\lambda$ based on permutations led to better results (Table S4).

Finally, we correctly estimated increases in the total bacterial abundances in extant host species ($\log \tilde{Y} > 0$; Figure S20). In contrast, decreases in the total bacterial abundances ($\log \tilde{Y} < 0$) were often estimated close to 0, indicating that our inferences may be biased toward positive $\log \tilde{Y}$ (*i.e.*, increase of the total microbial abundances). Therefore, because decreasing total abundances ($\log \tilde{Y} < 0$) are often poorly estimated, interpretations of $\log \tilde{Y}$ estimates are best avoided.

Similar results were obtained when directly simulating microbiota evolution on the empirical trees: we obtained accurate estimations of $\lambda$, especially at the level of bacterial orders (Figure S21), and accurate estimations of the ancestral relative bacterial abundances $Z_0$ (Figure S22). Covariances between bacterial taxa were also correctly estimated (Figure S23), with a low type-I error rate in general and a high statistical power for large covariances (Table S7).

**Supplementary Results 2:**

We tested the effect of captivity on our results, since it is known to impact gut microbiota composition at low taxonomic levels (McKenzie et al. 2017). In mammals, $\lambda$ values computed based on captive *versus* wild individuals were similar (Table S2). In birds, the inference based on captive individuals yielded a lower λ value, but this value was similar to that obtained based on wild individuals from the same number of species (Table S2). It thus seems that captivity does not blur phylosymbiosis at the level of bacterial orders.

We also assessed the presence of any systematic difference between studies in the Song et al.'s dataset by performing permutations, *i.e*. by shuffling the microbiota that originated from the same study and re-evaluating λ. The permutations led phylosymbiosis to vanish in birds and to drop significantly in mammals (Figure S24). The bird dataset is composed of only 4 different studies that each span the whole bird phylogenetic tree, whereas the mammal dataset is composed of 14 different studies (Song et al. 2020) that each tends to target a specific clade of mammals. This likely explains why the phylosymbiosis pattern vanished entirely in birds but not in mammals, where there is a strong phylogenetic signal in the sampling design. We concluded that the concatenation of different studies did not generate spurious phylosymbiosis.

#### Supplementary Tables:

**Supplementary Table 1: Simulations indicate that Mantel tests generally have lower statistical power and higher type-I error rates than our process-based approach when assessing phylosymbiosis.**

For each number of host species ($n$) and each number of bacterial taxa ($p$), we indicated the statistical power (proportion of true positives) and the type-I error rate (proportion of false positives) of Mantel tests when testing for phylosymbiosis in simulations. Mantel tests were performed using weighted Jaccard distances and Pearson correlations (Perez-Lamarque et al. 2022). We evaluated the significance of Mantel tests using 10,000 permutations. The type-I error rate was evaluated using simulations with $\lambda$=0, whereas statistical power was evaluated using $\lambda$=0.25, 0.5, 0.75, or 1. For each combination of $n$, $p$, and $\lambda$ values, we performed 100 simulations.

| n | p | Power (lambda=1) | Power (lambda=0.75) | Power (lambda=0.5) | Power (lambda=0.25) | Type–I error (lambda=0) |
|---|---|---|---|---|---|---|
| 20 | 3 | 0.83 | 0.53 | 0.35 | 0.15 | 0.06 |
| 50 | 3 | 0.96 | 0.77 | 0.65 | 0.32 | 0.07 |
| 100 | 3 | 0.95 | 0.93 | 0.79 | 0.51 | 0.05 |
| 250 | 3 | 0.98 | 0.93 | 0.87 | 0.64 | 0.06 |
| 20 | 5 | 0.93 | 0.68 | 0.41 | 0.22 | 0.05 |
| 50 | 5 | 1 | 0.96 | 0.74 | 0.39 | 0.03 |
| 100 | 5 | 0.98 | 0.98 | 0.93 | 0.59 | 0.04 |
| 250 | 5 | 1 | 0.99 | 0.97 | 0.83 | 0.05 |
| 20 | 10 | 0.97 | 0.91 | 0.6 | 0.3 | 0.07 |
| 50 | 10 | 1 | 0.99 | 0.93 | 0.58 | 0.04 |
| 100 | 10 | 1 | 1 | 0.93 | 0.82 | 0.06 |
| 250 | 10 | 1 | 1 | 1 | 0.95 | 0.06 |
| 20 | 15 | 1 | 0.92 | 0.77 | 0.4 | 0.06 |
| 50 | 15 | 1 | 1 | 0.96 | 0.64 | 0.07 |
| 100 | 15 | 1 | 1 | 1 | 0.88 | 0.05 |
| 250 | 15 | 1 | 1 | 1 | 0.99 | 0.03 |

**Supplementary Table 2: Significant phylosymbiosis in the gut microbiota of mammals and birds.**

For each host clade and condition we tested, we indicate the number of sampled species, and the estimated mean and 95% credible interval from the posterior distribution of Pagel's λ. The significance of λ is indicated by a p-value based on permutations. Results are for the 14 most abundant bacterial orders and 7 most abundant phyla.

| Class | Order | Type | Number of species | Pagel's λ (95% CI) | |
|---|---|---|---|---|---|
| | | | | Bacterial orders | Bacterial phyla |
| Mammalia | all | all | 215 | 0.65 [0.59, 0.70] p<0.01 | 0.78 [0.70, 0.85] p<0.01 |
| Mammalia | all | wild | 105 | 0.49 [0.41, 0.57] p<0.01 | 0.79 [0.73, 0.84] p<0.01 |
| Mammalia | all | captive | 127 | 0.49 [0.39, 0.58] p<0.01 | 0.77 [0.69, 0.84] p<0.01 |
| Mammalia | all | non-flying | 172 | 0.62 [0.55, 0.69] p<0.01 | 0.83 [0.78, 0.86] p<0.01 |
| Mammalia | Carnivora | all | 30 | 0.11 [0.02, 0.23] p=0.04 | 0.18 [0.02, 0.41] p=0.03 |
| Mammalia | Cetartiodactyla | all | 33 | 0.23 [0.10, 0.36] p<0.01 | 0.55 [0.25, 0.78] p=0.01 |
| Mammalia | Chiroptera | all | 43 | 0.17 [0.05, 0.30] p=0.01 | 0.2 [0.03, 0.45] p=0.08 |
| Mammalia | Primates | all | 50 | 0.30 [0.18, 0.42] p<0.01 | 0.44 [0.26, 0.61] p<0.01 |
| Mammalia | Rodentia | all | 17 | 0.11 [0.01, 0.31] p=0.41 | 0.74 [0.58, 0.85] p=0.09 |
| Aves | all | all | 323 | 0.32 [0.26, 0.38] p<0.01 | 0.66 [0.58, 0.73] p<0.01 |
| Aves | all | wild | 261 | 0.32 [0.26, 0.39] p<0.01 | 0.67 [0.59, 0.75] p<0.01 |
| Aves | all | wild (reduced number) | 64 | 0.21 [0.12, 0.31] p<0.01 | 0.47 [0.27, 0.65] p<0.01 |
| Aves | all | captive | 64 | 0.17 [0.07, 0.28] p<0.01 | 0.29 [0.09, 0.52] p<0.01 |
| Aves | all | flying | 312 | 0.32 [0.26, 0.38] p<0.01 | 0.66 [0.57, 0.73] p<0.01 |

| | | | | | |
|---|---|---|---|---|---|
| Aves | Anseriformes | all | 23 | 0.04 [0, 0.12] p=0.06 | 0.15 [0.01, 0.43] p=0.31 |
| Aves | Charadriiformes | all | 34 | 0.07 [0.01, 0.16] p=0.14 | 0.22 [0.04, 0.44] p=0.09 |
| Aves | Columbiformes | all | 15 | 0.22 [0.01, 0.52] p=0.01 | 0.57 [0.07, 0.97] p<0.01 |
| Aves | Passeriformes | all | 137 | 0.19 [0.12, 0.27] p<0.01 | 0.83 [0.78, 0.87] p<0.01 |

**Supplementary Table 3: Individuals from the same species have more similar gut microbiota compositions than individuals from different species.**

Results of a PermANOVA testing the effect of host species on Bray-Curtis dissimilarities of the gut microbiota compositions. PermANOVA were performed at the bacterial phylum level (using the 5 or 7 most abundant phyla) or order level (using the 9 or 14 most abundant orders) and separately for mammals and birds. For each PermANOVA, we reported the proportion of the variance ($R^2$) explained by host species and the associated p-value assessed based on 1,000 permutations.

| Bacterial taxonomy | Number of taxa | Clade | $R^2$ | p-value |
|---|---|---|---|---|
| Phylum | 5 | Mammals | 0.70 | <0.001 |
| | | Birds | 0.40 | <0.001 |
| | 7 | Mammals | 0.68 | <0.001 |
| | | Birds | 0.40 | <0.001 |
| Order | 9 | Mammals | 0.68 | <0.001 |
| | | Birds | 0.42 | <0.001 |
| | 14 | Mammals | 0.67 | <0.001 |
| | | Birds | 0.41 | <0.001 |

**Supplementary Table 4: Simulations validate the correct statistical performances of the evaluation of phylosymbiosis: low type-I error rate and high statistical power using randomizations.**

For each number of host species (n) and each number of bacterial taxa (p), we indicated the statistical power (proportion of true positives) and the type-I error rate (proportion of false positives) when testing for phylosymbiosis in simulations. We evaluated the significance by shuffling the species names 50 times and comparing the obtained $\lambda$. The type-I error rate was evaluated using simulations with $\lambda$=0, whereas statistical power was evaluated using $\lambda$=0.25, 0.5, 0.75, or 1. For each combination of $n$, $p$, and $\lambda$ values, we performed 100 simulations.

Because permutations are much more computationally intensive than computing Bayes factors, we only tested the permutation strategy for a subset of the simulations ($p = 5$ or $p = 10$).

| n | p | Power (lambda=1) | Power (lambda=0.75) | Power (lambda=0.5) | Power (lambda=0.25) | Type–I error (lambda=0) |
|---|---|---|---|---|---|---|
| 20 | 5 | 1 | 0.93 | 0.78 | 0.39 | 0.06 |
| 50 | 5 | 1 | 1 | 1 | 0.77 | 0.04 |
| 100 | 5 | 1 | 1 | 1 | 0.9 | 0 |
| 20 | 10 | 1 | 1 | 0.75 | 0.33 | 0 |
| 50 | 10 | 1 | 1 | 1 | 0.97 | 0.04 |
| 100 | 10 | 1 | 1 | 1 | 1 | 0 |

**Supplementary Table 5: Species sharing the same diet have similar microbiota.**

Results of a PermANOVA testing the effect of diet on Bray-Curtis dissimilarities of the gut microbiota compositions. PermANOVA were performed at the bacterial phylum level (using the 5 or 7 most abundant phyla) or order level (using the 9 or 14 most abundant orders) and separately for mammals and birds. For each PermANOVA, we reported the proportion of the variance ($R^2$) explained by diet and the associated p-value assessed based on 1,000 permutations.

| Bacterial taxonomy | Number of taxa | Clade | $R^2$ | p-value |
|---|---|---|---|---|
| Phylum | 5 | Mammals | 0.25 | <0.001 |
| | | Birds | 0.04 | <0.001 |
| | 7 | Mammals | 0.24 | <0.001 |
| | | Birds | 0.04 | <0.001 |
| Order | 9 | Mammals | 0.23 | <0.001 |
| | | Birds | 0.03 | <0.001 |
| | 14 | Mammals | 0.22 | <0.001 |
| | | Birds | 0.03 | <0.001 |

**Supplementary Table 6: Main shifts in the gut microbiota composition of mammals and birds.**

For each main shift that we observed in the microbiota occurring at the origin of a given host clade, we indicate, for each key bacterial taxa, the percentage of variation of the estimated proportion of the bacterial taxa that occurred between the stem and the crown nodes (most common recent ancestors, MRCA).

| Moderate to large shifts in microbiota composition | Percentage of variation at the MRCA of the clade |
|---|---|
| increased proportion of Enterobacteriales in Chiroptera | +341% |
| increased proportion of Mycoplasmatales in Chiroptera | +230% |
| increased proportion of Actinomycetales in Chiroptera | +54% |
| decreased proportion of Bacteroidales in Chiroptera | -89% |
| decreased proportion of Clostridiales in Chiroptera | -49% |
| increased proportion of Bacillales in Chiroptera | +21% |
| increased proportion of Lactobacillales in Chiroptera | +132% |
| increased proportion of Fusobacteriales in Carnivora | +152% |
| increased proportion of Erysipelotrichales in Cingulata | +37% |
| increased proportion of Erysipelotrichales in Primates | +6% |
| decreased proportion of Enterobacteriales in Simiiformes | -68% |
| decreased proportion of Pseudomonadales in Simiiformes | -22% |
| decreased proportion of Enterobacteriales in Ungulata | -90% |
| decreased proportion of Pseudomonadales in Ungulata | -45% |
| increased proportion of Enterobacteriales in Passeriformes | +23% |
| increased proportion of Pseudomonadales in Passeriformes | +20% |
| increased proportion of Bacillales in Passeriformes | +3% |
| decreased proportion of Bacteroidales in Passeriformes | -43% |
| increased proportion of Bacteroidales in Anseriformes | +2% |
| increased proportion of Fusobacteriales in Anseriformes | +12% |
| increased proportion of Bacteroidales in Charadriiforms | +40% |
| increased proportion of Fusobacteriales in Charadriiforms | +164% |
| increased proportion of Actinobacteria in Columbiformes | +152% |

**Supplementary Table 7: Simulations on empirical trees validate the correct estimations of the covariances between bacterial taxa:**

For each category of simulations (mammals/birds and phyla/orders), we indicated the proportion of negative and positive simulated covariances (R) that were estimated as being (i) significantly negative, (ii) non-significant (n.s.), or (iii) significantly positive. A covariance is estimated as significantly positive (resp. negative) if the whole 95% credible interval is positive (resp. negative). On rows, we distinguished the small covariances ($-1 < R < 0$ and $0 < R < 1$) or large covariances ($R > 1$ or $R < -1$).
We highlighted in red the statistical power for the negative covariances, and in blue, the statistical power for the positive covariances. Type-I error rates correspond to negative (*resp.* positive) covariances estimated as being negative (*resp.* positive).

mammals - phyla

| sim.\est. | – | n.s. | + |
|---|---|---|---|
| ++ | 0.09 | 0.33 | 0.59 |
| + | 0.17 | 0.51 | 0.31 |
| – | 0.4 | 0.49 | 0.11 |
| –– | 0.74 | 0.23 | 0.03 |

birds - phyla

| sim.\est. | – | n.s. | + |
|---|---|---|---|
| ++ | 0.09 | 0.3 | 0.61 |
| + | 0.2 | 0.48 | 0.32 |
| – | 0.47 | 0.43 | 0.1 |
| –– | 0.82 | 0.16 | 0.02 |

mammals - orders

| sim.\est. | – | n.s. | + |
|---|---|---|---|
| ++ | 0.01 | 0.28 | 0.71 |
| + | 0.04 | 0.65 | 0.31 |
| – | 0.29 | 0.66 | 0.06 |
| –– | 0.75 | 0.24 | 0.01 |

birds - orders

| sim.\est. | – | n.s. | + |
|---|---|---|---|
| ++ | 0.01 | 0.22 | 0.77 |
| + | 0.05 | 0.61 | 0.34 |
| – | 0.37 | 0.59 | 0.04 |
| –– | 0.83 | 0.16 | 0 |

**Supplementary Table 8: Simulations validate the correct estimations of the covariances between bacterial taxa when n and p are large:**

For each number of host species (n) and each number of bacterial taxa (p), we indicated the proportion of negative and positive simulated covariances (R) that were estimated as being (i) significantly negative, (ii) non-significant (n.s.), or (iii) significantly positive. A covariance is estimated as significantly positive (resp. negative) if the whole 95% credible interval is positive (resp. negative). On rows, we distinguished the small covariances (-1<R<0 and 0<R<1) or large covariances (R>1 or R<-1). For each combination of n and p values, we performed 500 simulations (with $\lambda$=0, 0.25, 0.5, 0.75, or 1).
We highlighted in red the statistical power for the negative covariances, and in blue, the statistical power for the positive covariances. Type-I error rates correspond to negative (resp. positive) covariances estimated as being negative (resp. positive).

n=20; p=3

| sim.\est. | – | n.s. | + |
|---|---|---|---|
| ++ | 0.09 | 0.82 | 0.09 |
| + | 0.09 | 0.87 | 0.04 |
| – | 0.18 | 0.8 | 0.02 |
| –– | 0.48 | 0.52 | 0 |

n=50; p=3

| sim.\est. | – | n.s. | + |
|---|---|---|---|
| ++ | 0.21 | 0.62 | 0.17 |
| + | 0.26 | 0.64 | 0.09 |
| – | 0.47 | 0.49 | 0.04 |
| –– | 0.76 | 0.24 | 0 |

n=100; p=3

| sim.\est. | – | n.s. | + |
|---|---|---|---|
| ++ | 0.33 | 0.44 | 0.23 |
| + | 0.42 | 0.49 | 0.09 |
| – | 0.62 | 0.33 | 0.05 |
| –– | 0.82 | 0.15 | 0.02 |

n=250; p=3

| sim.\est. | – | n.s. | + |
|---|---|---|---|
| ++ | 0.44 | 0.3 | 0.26 |
| + | 0.53 | 0.31 | 0.16 |
| – | 0.77 | 0.19 | 0.05 |
| –– | 0.93 | 0.05 | 0.02 |

n=20; p=5

| sim.\est. | – | n.s. | + |
|---|---|---|---|
| ++ | 0.01 | 0.85 | 0.15 |
| + | 0.02 | 0.94 | 0.04 |
| – | 0.07 | 0.92 | 0.01 |
| –– | 0.3 | 0.7 | 0 |

n=50; p=5

| sim.\est. | – | n.s. | + |
|---|---|---|---|
| ++ | 0.05 | 0.66 | 0.29 |
| + | 0.08 | 0.83 | 0.1 |
| – | 0.24 | 0.74 | 0.02 |
| –– | 0.58 | 0.41 | 0.01 |

n=100; p=5

| sim.\est. | – | n.s. | + |
|---|---|---|---|
| ++ | 0.07 | 0.54 | 0.39 |
| + | 0.17 | 0.66 | 0.17 |
| – | 0.38 | 0.57 | 0.05 |
| –– | 0.74 | 0.25 | 0.01 |

n=250; p=5

| sim.\est. | – | n.s. | + |
|---|---|---|---|
| ++ | 0.16 | 0.33 | 0.51 |
| + | 0.29 | 0.47 | 0.24 |
| – | 0.54 | 0.37 | 0.09 |
| –– | 0.83 | 0.15 | 0.02 |

n=20; p=10

| sim.\est. | – | n.s. | + |
|---|---|---|---|
| ++ | 0 | 0.77 | 0.23 |
| + | 0 | 0.94 | 0.06 |
| – | 0.04 | 0.95 | 0.01 |
| –– | 0.25 | 0.75 | 0 |

n=50; p=10

| sim.\est. | – | n.s. | + |
|---|---|---|---|
| ++ | 0 | 0.61 | 0.38 |
| + | 0.02 | 0.87 | 0.11 |
| – | 0.11 | 0.88 | 0.01 |
| –– | 0.47 | 0.53 | 0 |

n=100; p=10

| sim.\est. | – | n.s. | + |
|---|---|---|---|
| ++ | 0.01 | 0.47 | 0.52 |
| + | 0.03 | 0.79 | 0.18 |
| – | 0.2 | 0.77 | 0.02 |
| –– | 0.66 | 0.34 | 0 |

n=250; p=10

| sim.\est. | – | n.s. | + |
|---|---|---|---|
| ++ | 0.03 | 0.32 | 0.65 |
| + | 0.09 | 0.64 | 0.27 |
| – | 0.37 | 0.58 | 0.05 |
| –– | 0.8 | 0.19 | 0.01 |

n=20; p=15

| sim.\est. | – | n.s. | + |
|---|---|---|---|
| ++ | 0 | 0.74 | 0.25 |
| + | 0.01 | 0.93 | 0.07 |
| – | 0.05 | 0.94 | 0.01 |
| –– | 0.24 | 0.76 | 0 |

n=50; p=15

| sim.\est. | – | n.s. | + |
|---|---|---|---|
| ++ | 0 | 0.58 | 0.42 |
| + | 0.01 | 0.89 | 0.1 |
| – | 0.09 | 0.9 | 0.01 |
| –– | 0.48 | 0.52 | 0 |

n=100; p=15

| sim.\est. | – | n.s. | + |
|---|---|---|---|
| ++ | 0 | 0.41 | 0.59 |
| + | 0.01 | 0.83 | 0.16 |
| – | 0.16 | 0.83 | 0.01 |
| –– | 0.67 | 0.33 | 0 |

n=250; p=15

| sim.\est. | – | n.s. | + |
|---|---|---|---|
| ++ | 0 | 0.26 | 0.74 |
| + | 0.03 | 0.69 | 0.28 |
| – | 0.32 | 0.65 | 0.03 |
| –– | 0.83 | 0.17 | 0 |

Average

| sim.\est. | – | n.s. | + |
|---|---|---|---|
| ++ | 0.09 | 0.54 | 0.37 |
| + | 0.13 | 0.74 | 0.13 |
| – | 0.29 | 0.68 | 0.03 |
| –– | 0.61 | 0.38 | 0.01 |

Average (when p>5)

| sim.\est. | – | n.s. | + |
|---|---|---|---|
| ++ | 0.01 | 0.52 | 0.47 |
| + | 0.02 | 0.82 | 0.15 |
| – | 0.17 | 0.81 | 0.02 |
| –– | 0.55 | 0.45 | 0 |

**Supplementary Table 9: Simulations indicate that the evaluation of phylosymbiosis using Bayes factors has both a low type-I error rate and a low statistical power.**

For each number of host species (n) and each number of bacterial taxa (p), we indicated the statistical power (proportion of true positives) and the type-I error rate (proportion of false positives) when testing for phylosymbiosis in simulations. We evaluated the significance of phylosymbiosis using Bayes factors by testing the support of the null model with $\lambda$=0. The type-I error rate was evaluated using simulations with $\lambda$=0, whereas statistical power was evaluated using $\lambda$=0.25, 0.5, 0.75, or 1. For each combination of n, p, and $\lambda$ values, we performed 100 simulations.

| n | p | Power (lambda=1) | Power (lambda=0.75) | Power (lambda=0.5) | Power (lambda=0.25) | Type–I error (lambda=0) |
|---|---|---|---|---|---|---|
| 20 | 3 | 0.83 | 0.12 | 0.01 | 0 | 0 |
| 50 | 3 | 1 | 0.41 | 0.01 | 0 | 0 |
| 100 | 3 | 1 | 0.78 | 0.03 | 0 | 0 |
| 250 | 3 | 1 | 0.98 | 0.12 | 0 | 0 |
| 20 | 5 | 0.96 | 0.27 | 0.01 | 0 | 0 |
| 50 | 5 | 1 | 0.83 | 0.11 | 0 | 0 |
| 100 | 5 | 1 | 0.98 | 0.26 | 0 | 0 |
| 250 | 5 | 1 | 1 | 0.61 | 0 | 0 |
| 20 | 10 | 0.96 | 0.17 | 0 | 0 | 0 |
| 50 | 10 | 1 | 0.99 | 0.31 | 0 | 0 |
| 100 | 10 | 1 | 1 | 0.83 | 0.03 | 0 |
| 250 | 10 | 1 | 1 | 1 | 0.13 | 0 |
| 20 | 15 | 0.7 | 0.01 | 0 | 0 | 0.02 |
| 50 | 15 | 0.97 | 0.98 | 0.33 | 0.01 | 0.01 |
| 100 | 15 | 0.96 | 1 | 0.96 | 0.12 | 0 |
| 250 | 15 | 0.99 | 1 | 1 | 0.47 | 0 |

### Supplementary Figures:

#### Empirical application:

##### Supplementary Figure 1: Phylosymbiosis evaluated in the gut microbiota of mammals and birds:

We evaluated the levels of phylosymbiosis ($\lambda$) in mammals and birds (a) or in major orders of mammals and birds (b). For each clade, inferences were performed at the bacterial phylum level (using the 7 most abundant ones) or order level (using the 14 most abundant ones). Inferences in birds were run using (i) all species, (ii) only species with gut microbiota sampled in wild individuals, and (iii) only species with gut microbiota sampled in captive individuals. Inferences were also performed on non-flying mammal species only (*i.e.* excluding bats) and on flying bird species only to exclude the effect of flying ability. Because only a small fraction of bird species was sampled in captivity (66 species), we additionally performed inferences on 66 bird species sampled in wild individuals ("wild - low"), to exclude any bias of the number of host species on the estimation of $\lambda$.

We reported the mean $\lambda$ values and their associated 95% credible intervals. The significances of the phylosymbiosis values were evaluated in Table S2.

Results were qualitatively similar when using only the 5 or 9 most abundant phyla or orders respectively.

(a)

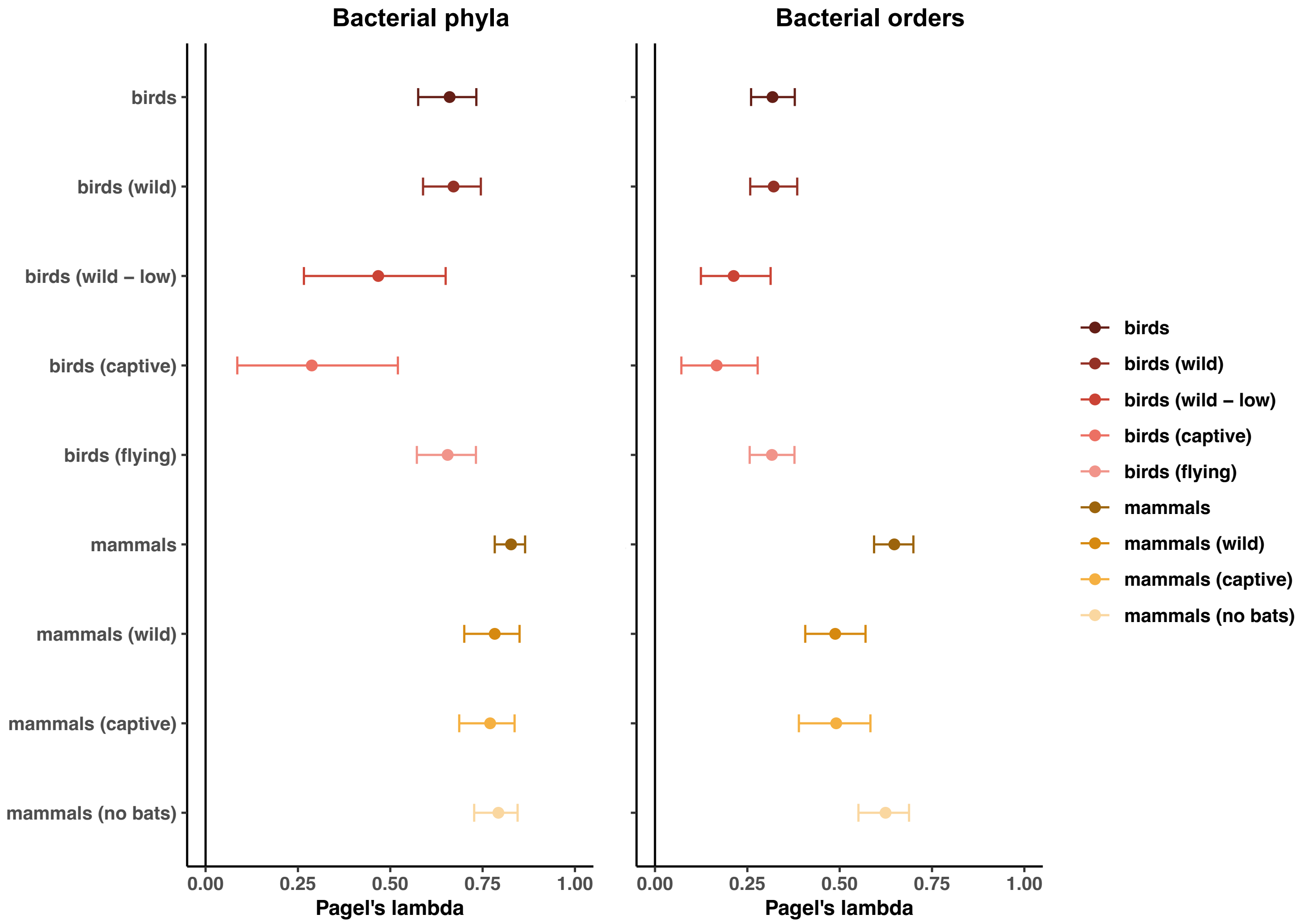
Bacterial phyla
Bacterial orders
birds
birds (wild)
birds (wild – low)
birds (captive)
birds (flying)
mammals
mammals (wild)
mammals (captive)
mammals (no bats)
0.00
0.25
0.50
0.75
1.00
Pagel's lambda
Pagel's lambda

(b)

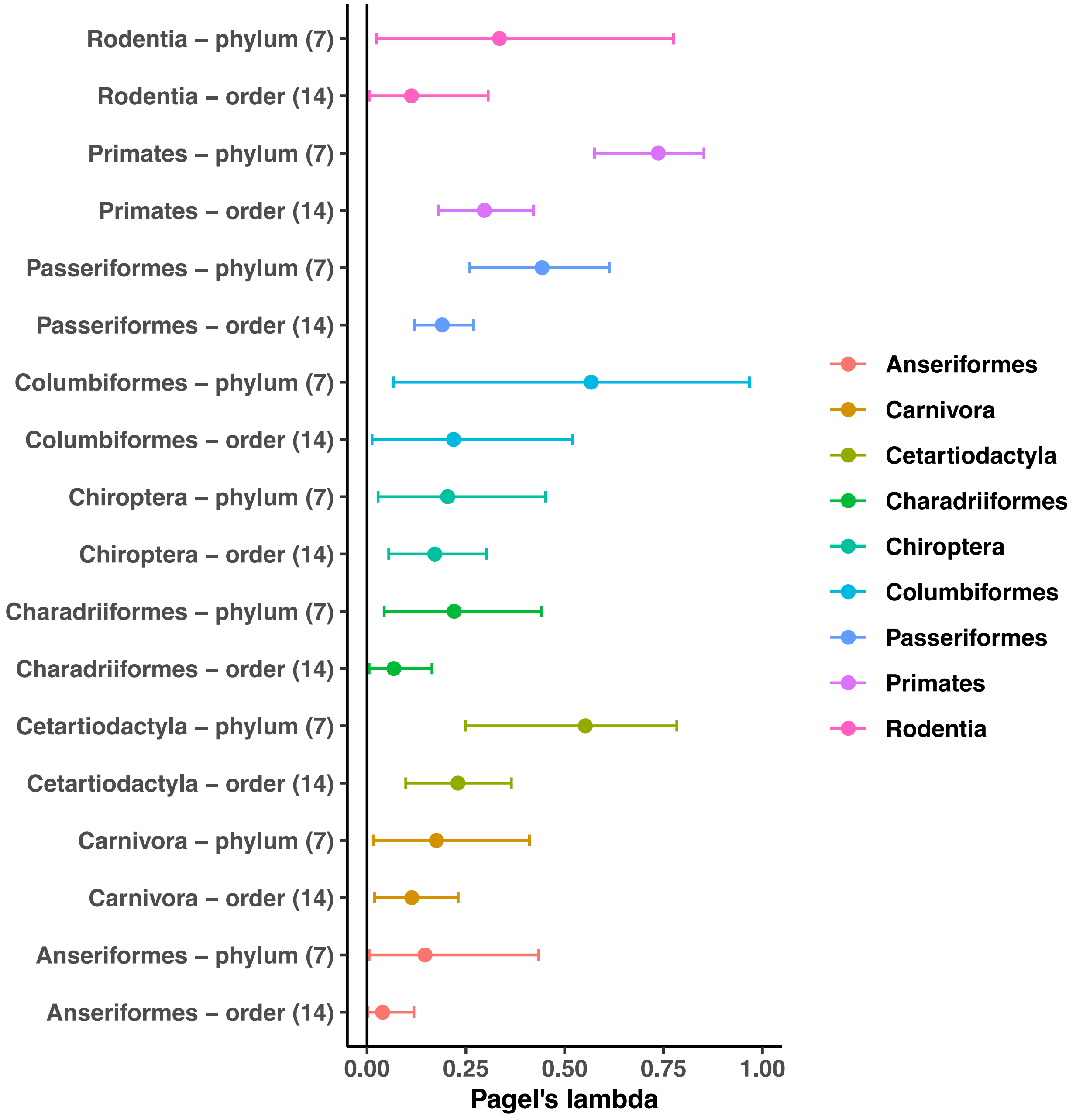
Rodentia – phylum (7)
Rodentia – order (14)
Primates – phylum (7)
Primates – order (14)
Passeriformes – phylum (7)
Passeriformes – order (14)
Columbiformes – phylum (7)
Columbiformes – order (14)
Chiroptera – phylum (7)
Chiroptera – order (14)
Charadriiformes – phylum (7)
Charadriiformes – order (14)
Cetartiodactyla – phylum (7)
Cetartiodactyla – order (14)
Carnivora – phylum (7)
Carnivora – order (14)
Anseriformes – phylum (7)
Anseriformes – order (14)
0.00
0.25
0.50
0.75
1.00
Pagel's lambda
Anseriformes
Carnivora
Cetartiodactyla
Charadriiformes
Chiroptera
Columbiformes
Passeriformes
Primates
Rodentia

**Supplementary Figure 2: At the bacterial phylum level, phylogenetically-conserved diets, geographic locations, or flying abilities may only partially contribute to the phylosymbiosis in the gut microbiota of mammals and birds.**

For both mammals and birds, we compared the estimated level of phylosymbiosis (mean λ value in orange) to levels of phylosymbiosis (λ values) estimated when shuffling the species having the same diet (green boxplot), the same geographic location (blue boxplot), flying or non-flying species (purple boxplot), or all of the latter (in red). For each shuffling strategy, we performed 100.
Boxplots indicate the median surrounded by the first and third quartiles, and whiskers extend to the extreme values but no further than 1.5 of the inter-quartile range.
Randomizations were performed on gut microbiota at the phylum level using the 7 most abundant phyla. Results were qualitatively similar when using only the 5 or 9 most abundant phyla or orders respectively.

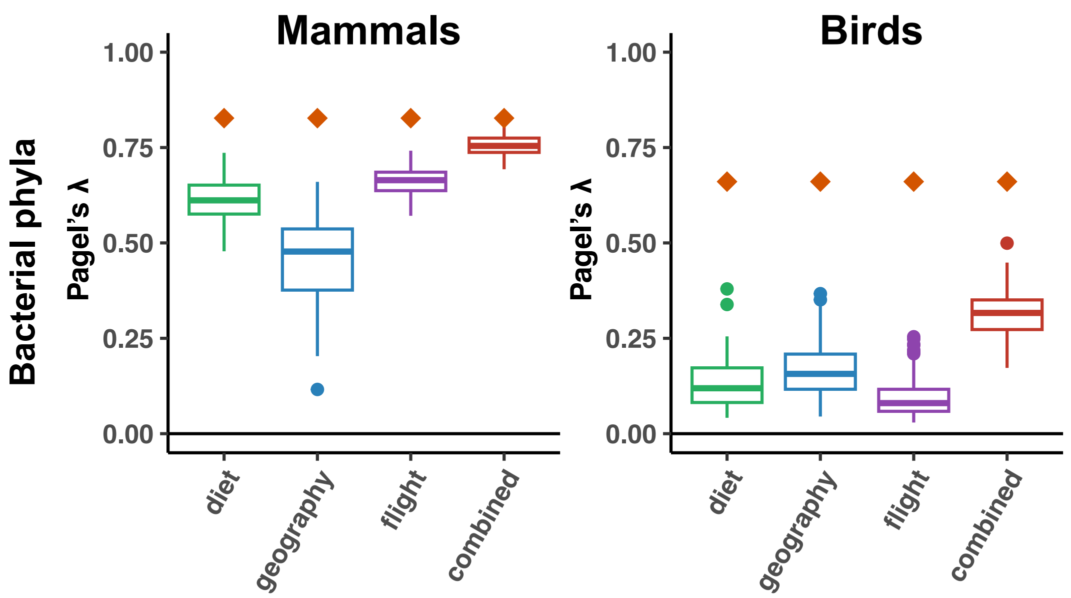

**Supplementary Figure 3: Phylogenetically-conserved diets, geographic locations, or flying abilities only partially contribute to the phylosymbiosis in the gut microbiota of mammal and bird orders.**

For large orders of mammals or birds, we compared the estimated level of phylosymbiosis (mean λ value in orange) to levels of phylosymbiosis (λ values) estimated when shuffling the species having the same diet (green boxplot) or the same geographic location (blue boxplot). For each shuffling strategy, we performed 100 randomizations. Randomizations were performed on gut microbiota at the phylum and order levels, using the 7 most abundant phyla or the 14 most abundant phyla respectively. Results were qualitatively similar when using only the 5 or 9 most abundant phyla or orders respectively. Boxplots indicate the median surrounded by the first and third quartiles, and whiskers extend to the extreme values but no further than 1.5 of the inter-quartile range.

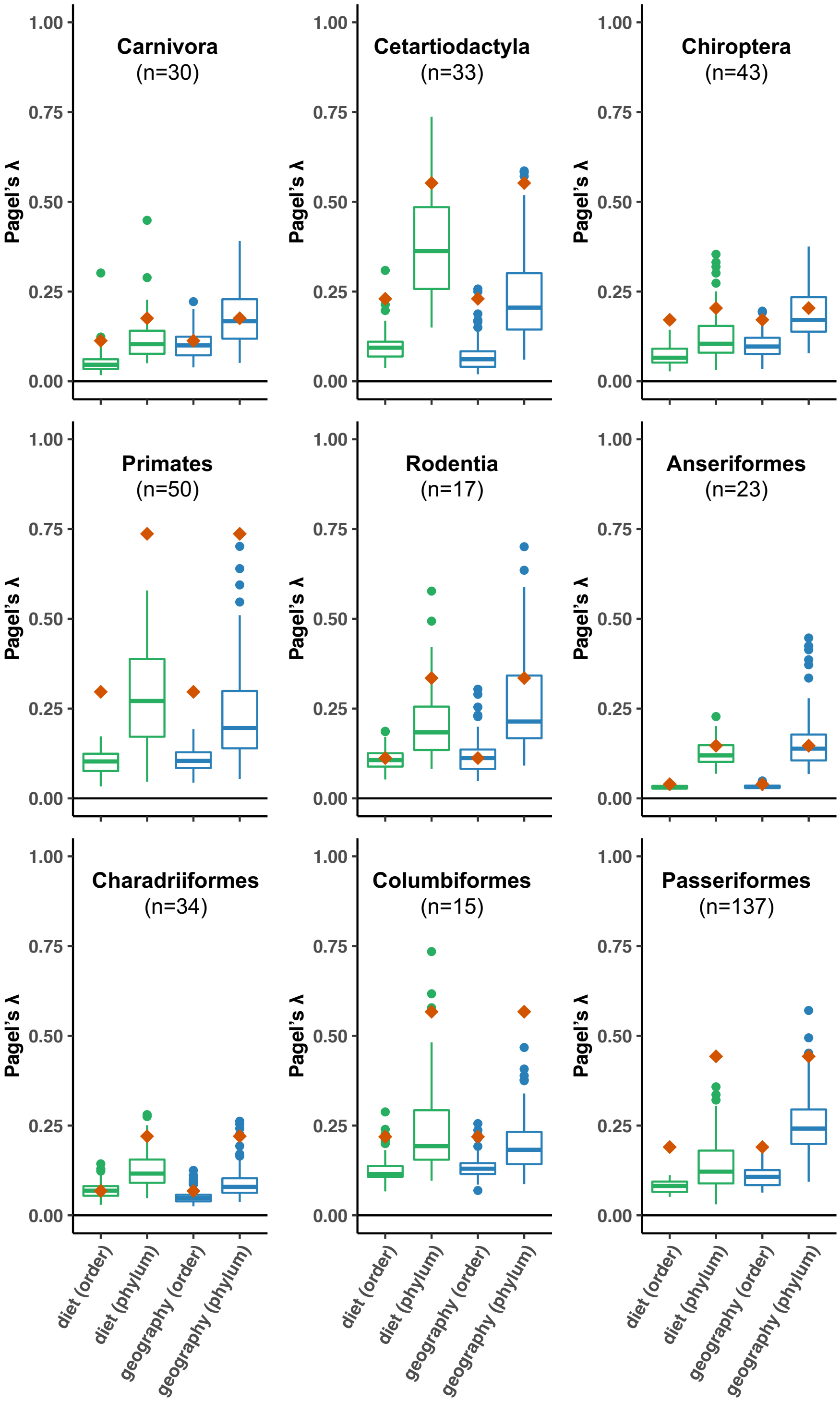

**Supplementary Figure 4: Ancestral gut microbiota compositions estimated for mammals and birds.**

Estimations of the relative abundances of the 5 or 7 most abundant bacterial phyla or the 9 or 14 most abundant bacterial orders (second row) in the gut microbiota of the most recent common ancestor of mammals and birds.
Because the ancestral microbiota composition of all mammals is quite uncertain (because of the large evolutionary time separating eutherians and marsupials), we also represent the ancestral microbiota compositions of Eutheria and Marsupials only.

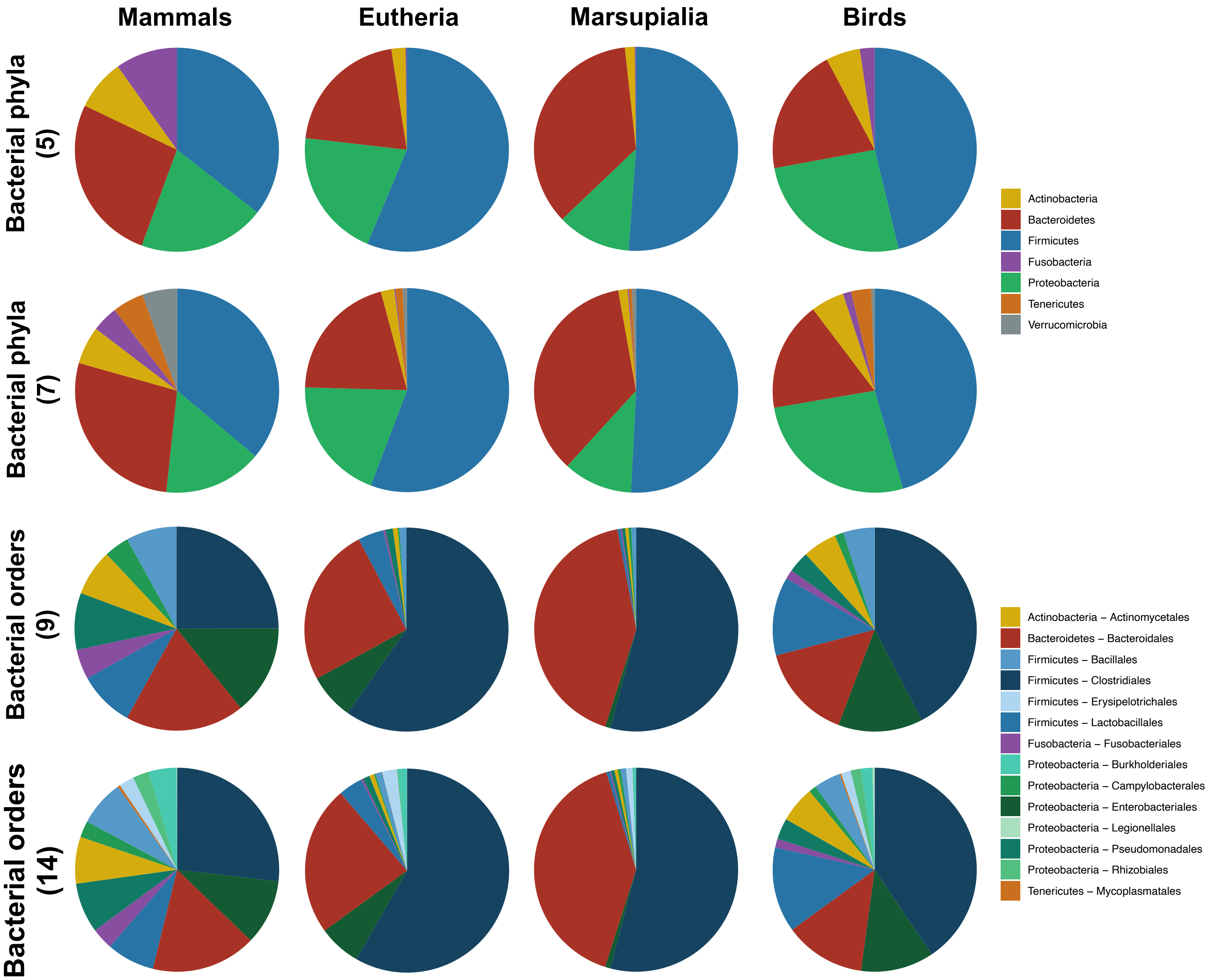

**Supplementary Figure 5: Ancestral gut microbiota compositions estimated for mammals and birds at the bacterial phylum level.**

Estimations of the relative abundances of the 7 most abundant bacterial phyla in the gut microbiota of the ancestors of mammals or birds. We inferred the ancestral abundances at each node of the phylogenetic trees using generalized least squares (following Martins & Hansen 1997, Cunningham et al. 1998, Clavel et al. 2018). Results were qualitatively similar when using only the 5 or 9 most abundant phyla or orders respectively.

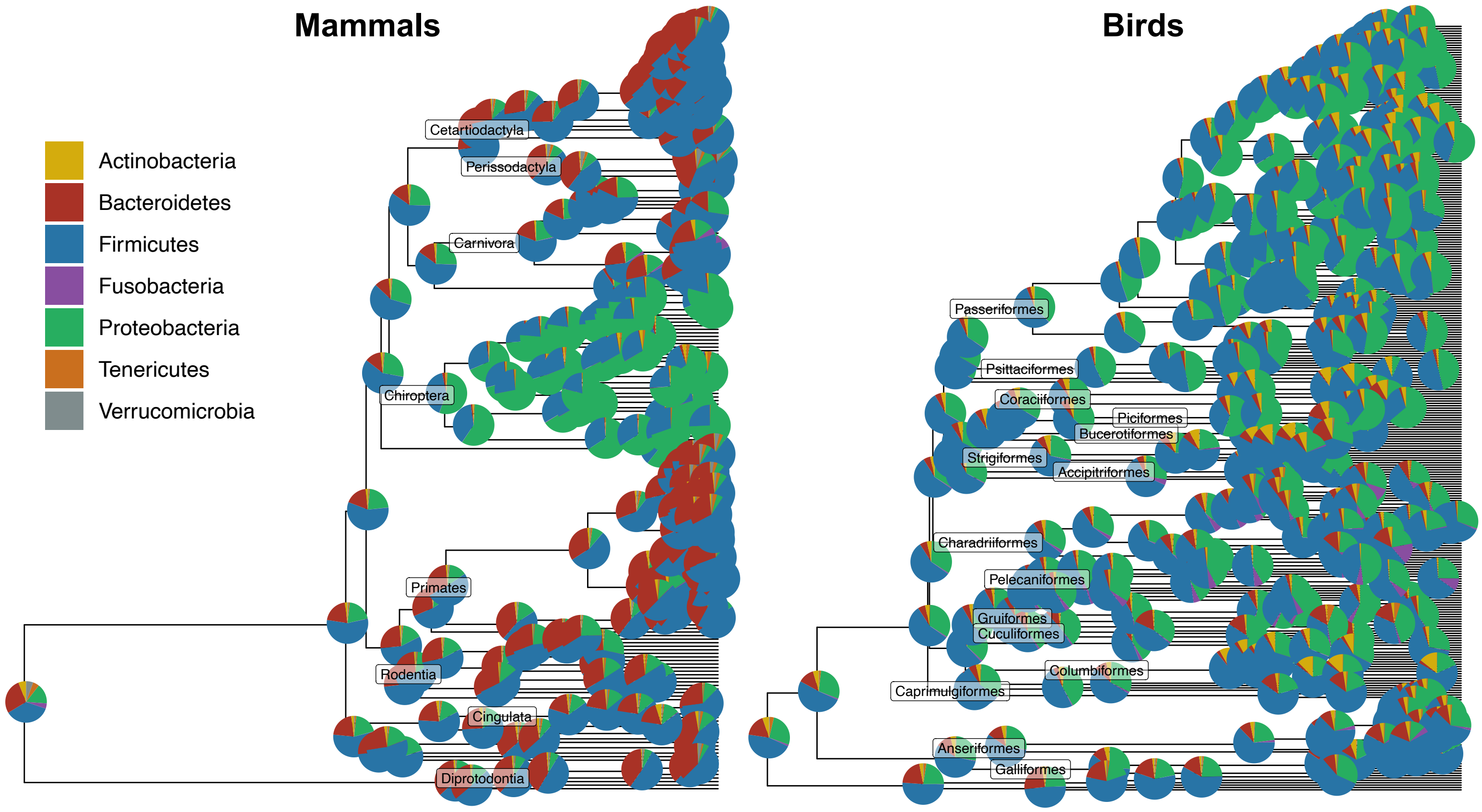

**Supplementary Figure 6: Compared with birds, estimations of the ancestral microbiota composition of mammal guts present a larger uncertainty.**

**(a)** 10 ancestral compositions (Z0) were randomly sampled in the posteriors of the inferences performed for mammals or birds at the phylum or order level. Results were qualitatively similar when using only the 5 or 9 most abundant phyla or orders respectively.

**(b)** Principal component analyses (PCA) of the gut microbiota of extant mammal (left) and bird species (right), plus the estimated microbiota compositions of the most recent common ancestor of mammals and birds (blue dots): 100 ancestral compositions sampled from the posterior are represented in blue (the mean value of the posterior is in dark blue). PCA were performed on the relative abundances of the 7 most abundant bacterial phyla (first rows) or of the 14 most abundant bacterial orders (second rows) following a centered log ratio transform.

(a) Posterior of ancestral compositions:

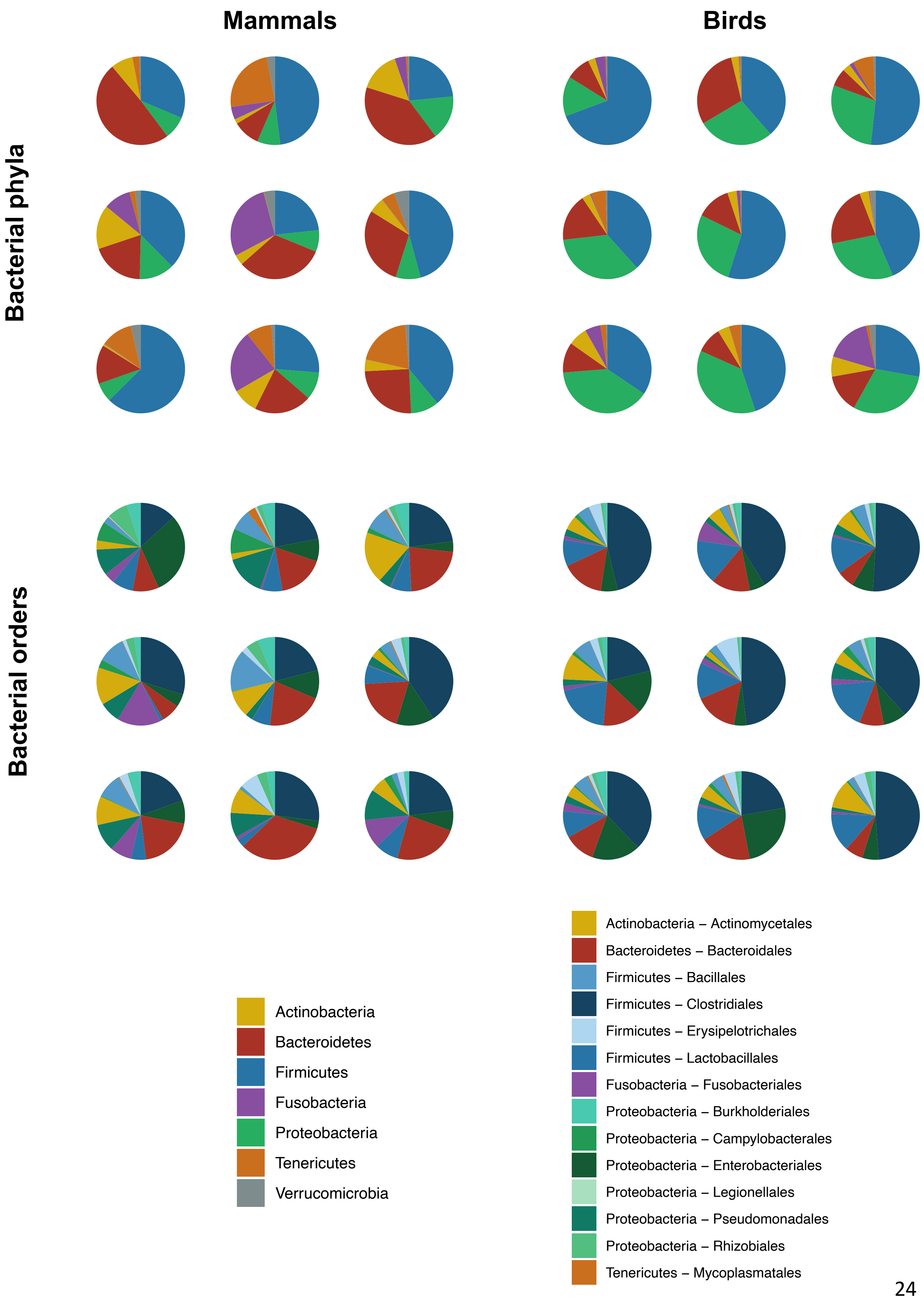

**(b) PCA including the posterior of ancestral compositions:**

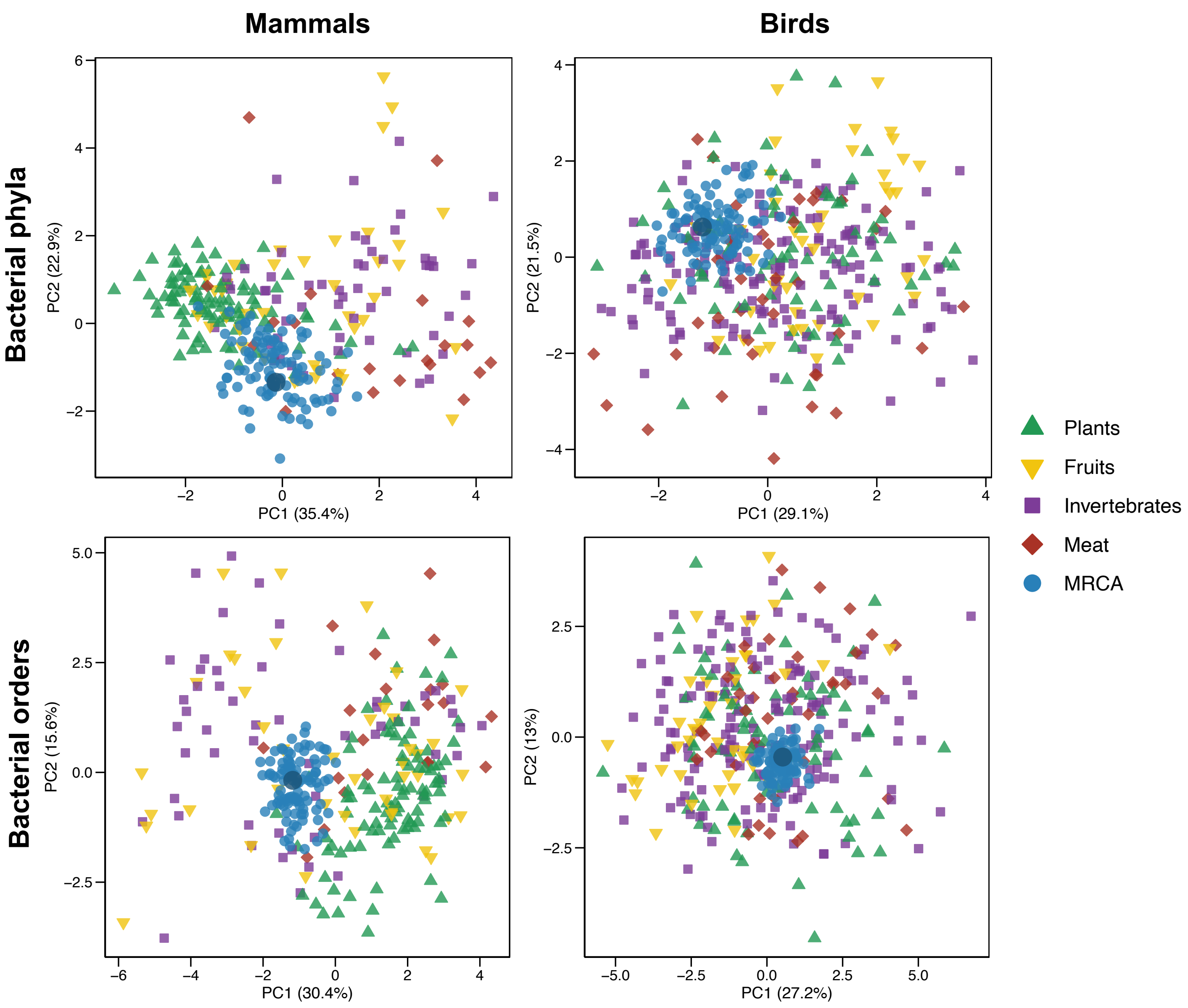

**Supplementary Figure 7: Projection of the estimated ancestral gut microbiota of mammals and birds onto the space of present-day gut microbiota sampled from both wild and captive individuals: Ancestral gut microbiota of mammals tend to be similar to the gut microbiota of extant species with invertebrate-based diets.**

Principal component analyses (PCA) of the gut microbiota of extant mammal (left) and bird species (right), plus the estimated microbiota of the most recent common ancestor of mammals and birds (blue dots). PCA were performed on the relative abundances of the 7 most abundant bacterial phyla (first rows) or of the 14 most abundant bacterial orders (second rows) following a centered log ratio transform. Results were qualitatively similar when using only the 5 or 9 most abundant phyla or orders respectively.

**(a) Projection of bacterial taxa contributing to the two principal components (PC).** Colors represent the contribution of the taxa to the principal components. The percentage for each principal component (PC) indicates its amount of explained variance.

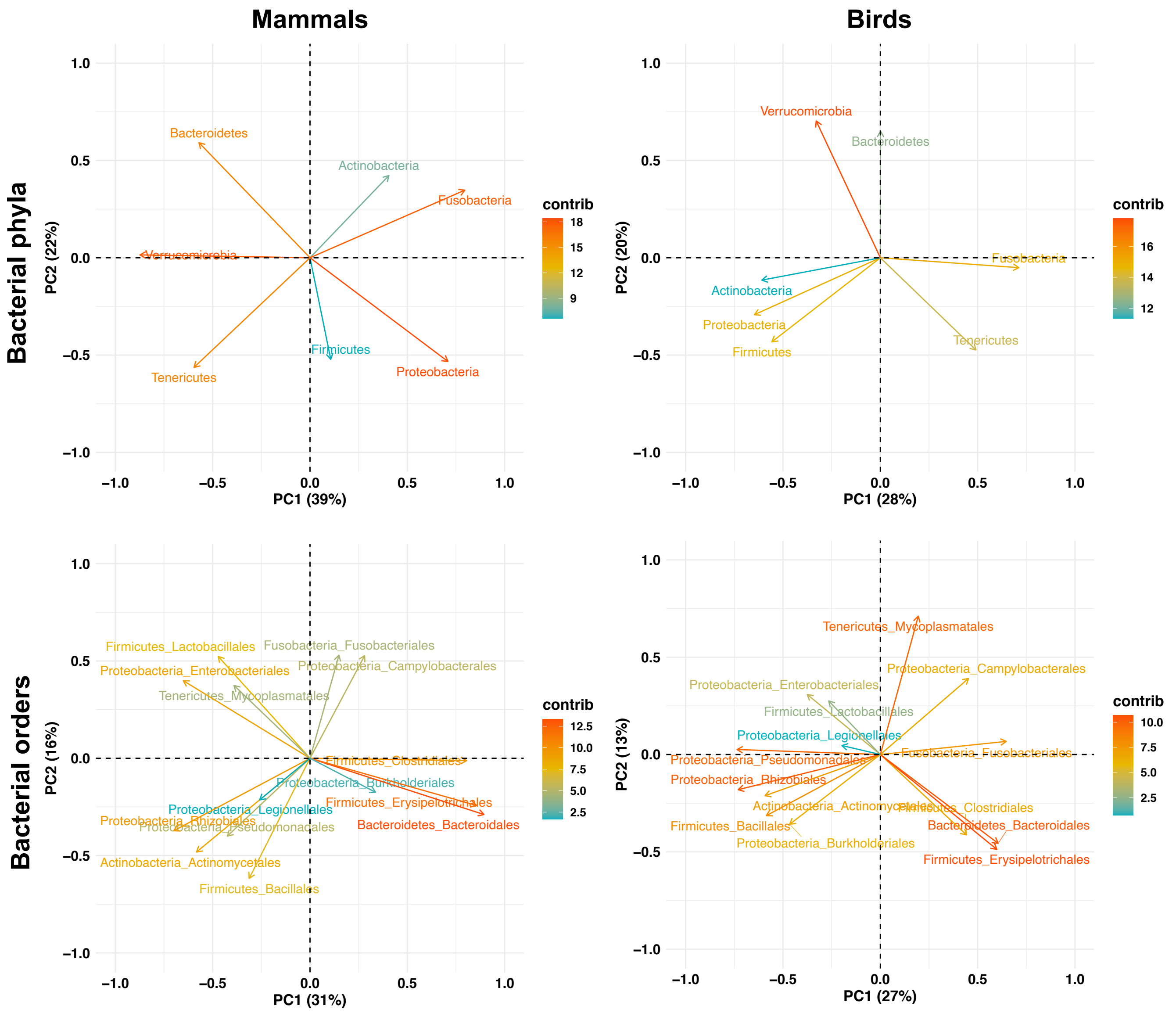

**(b) Projection of the extant and ancestral microbiota compositions.** Extant microbiota are colored according to the species diet. Ancestral microbiota compositions of birds and mammals (including that of Eutheria and Marsupialia) are represented in blue. For each PCA plot, we indicated the three extant species with microbiota compositions that are the closest to the estimated ancestral microbiota composition.

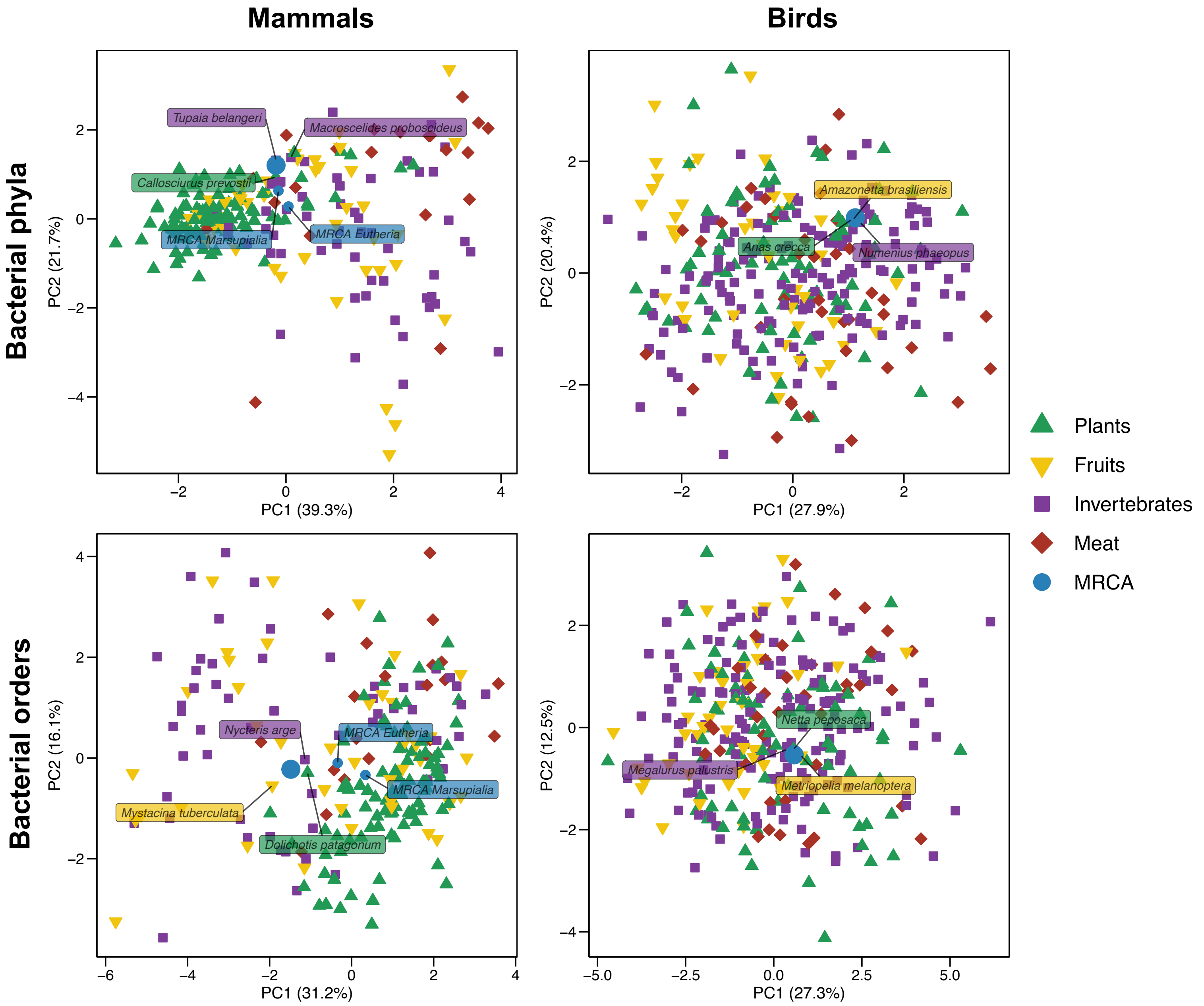

**(c) Estimated ancestral gut microbiota are distinct from the average microbiota of extant species:** Projection of the extant and ancestral microbiota compositions. Extant microbiota are colored according to the species diet. Ancestral microbiota compositions of birds and mammals are represented in blue and the average microbiota compositions of extant species are represented in orange.

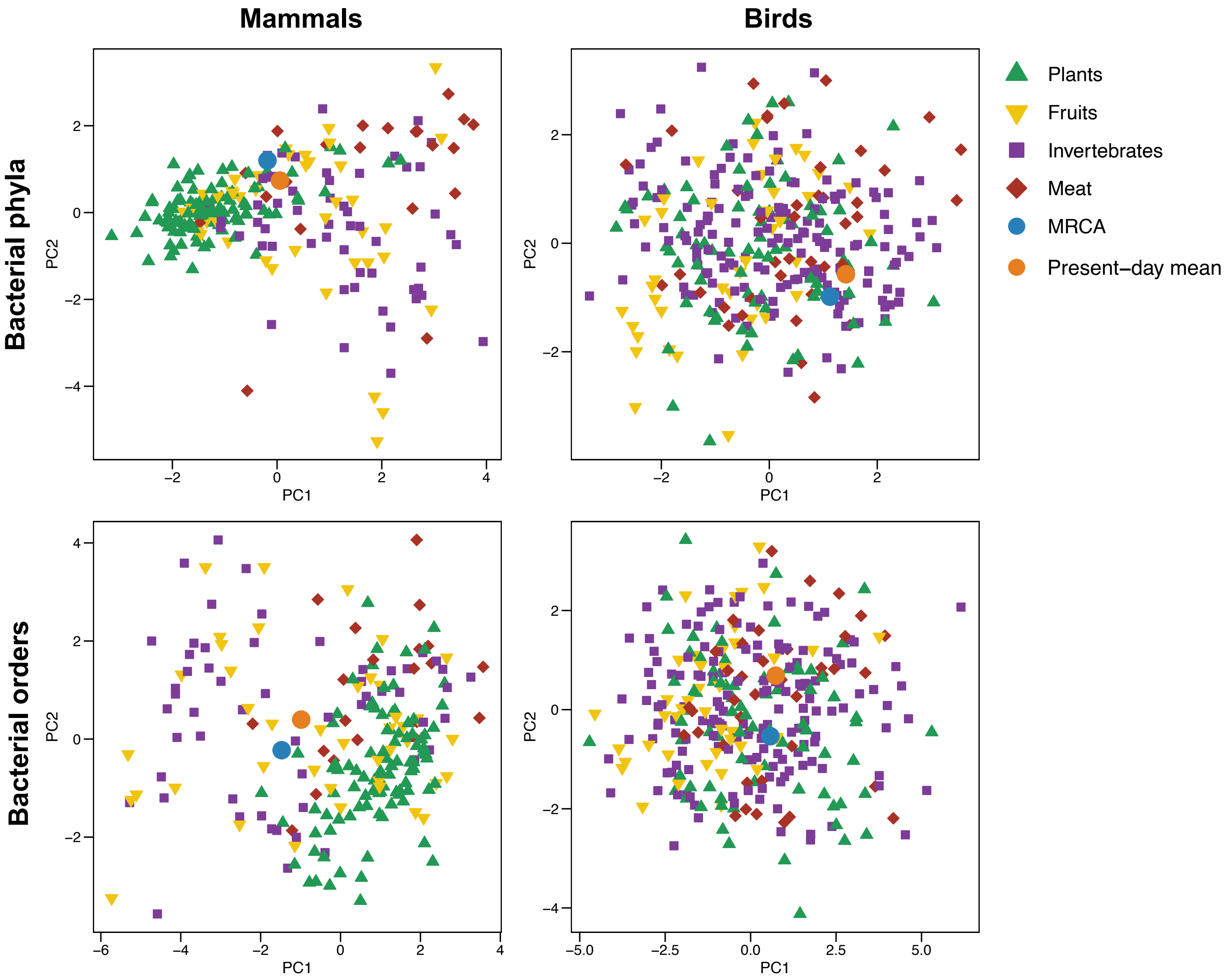

**Supplementary Figure 8: Projection of the estimated ancestral gut microbiota of mammals and birds onto the space of present-day gut microbiota sampled from both wild individuals only: At the bacterial phylum level, ancestral gut microbiota of mammals also tend to be similar to the gut microbiota of extant species with invertebrate-based diets.**

Principal component analyses (PCA) of the gut microbiota of extant mammal (left) and bird species (right), plus the estimated microbiota of the most recent common ancestor of mammals and birds (blue dots). PCA were performed on the relative abundances of the 7 most abundant bacterial phyla following a centered log ratio transform. Results were qualitatively similar when using only the 5 or 9 most abundant phyla or orders respectively.

**(a) Projection of bacterial taxa contributing to the two principal components (PC).** Colors represent the contribution of the taxa to the principal components. The percentage for each principal component (PC) indicates its amount of explained variance.

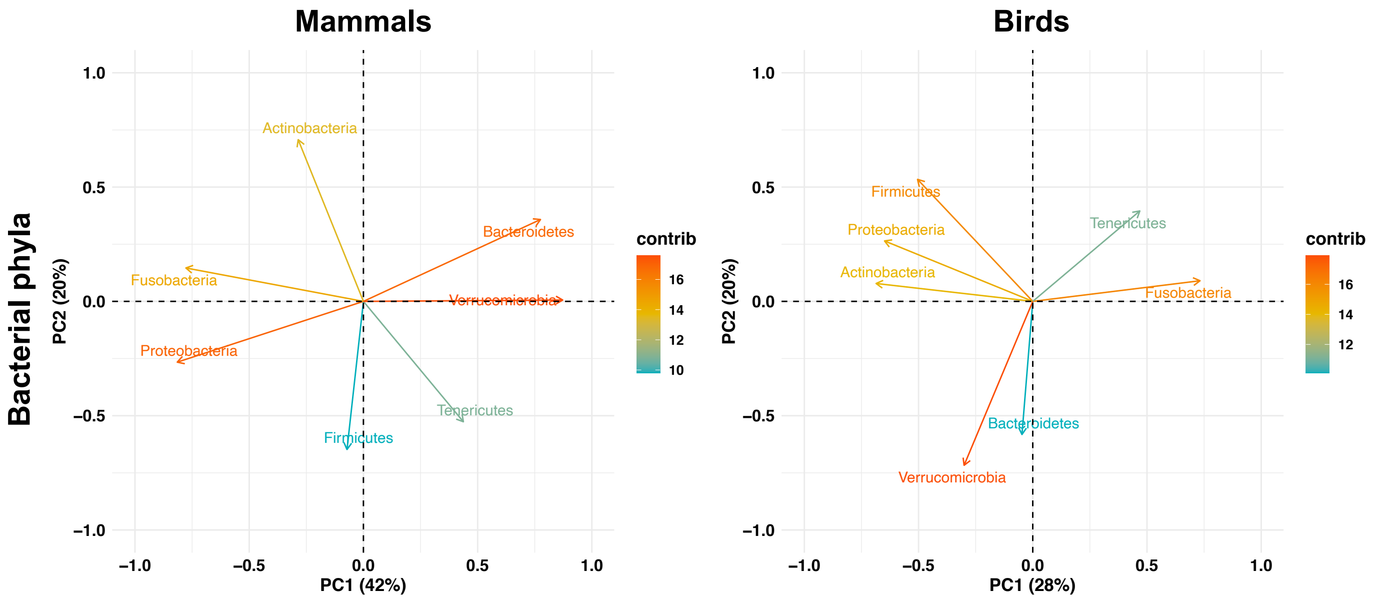

**(b) Projection of the extant and ancestral microbiota compositions.** Extant microbiota are colored according to the species diet. Ancestral microbiota compositions of birds and mammals (including that of Eutheria and Marsupialia) are represented in blue. For each PCA plot, we indicated the three extant species with microbiota compositions that are the closest to the estimated ancestral microbiota composition.

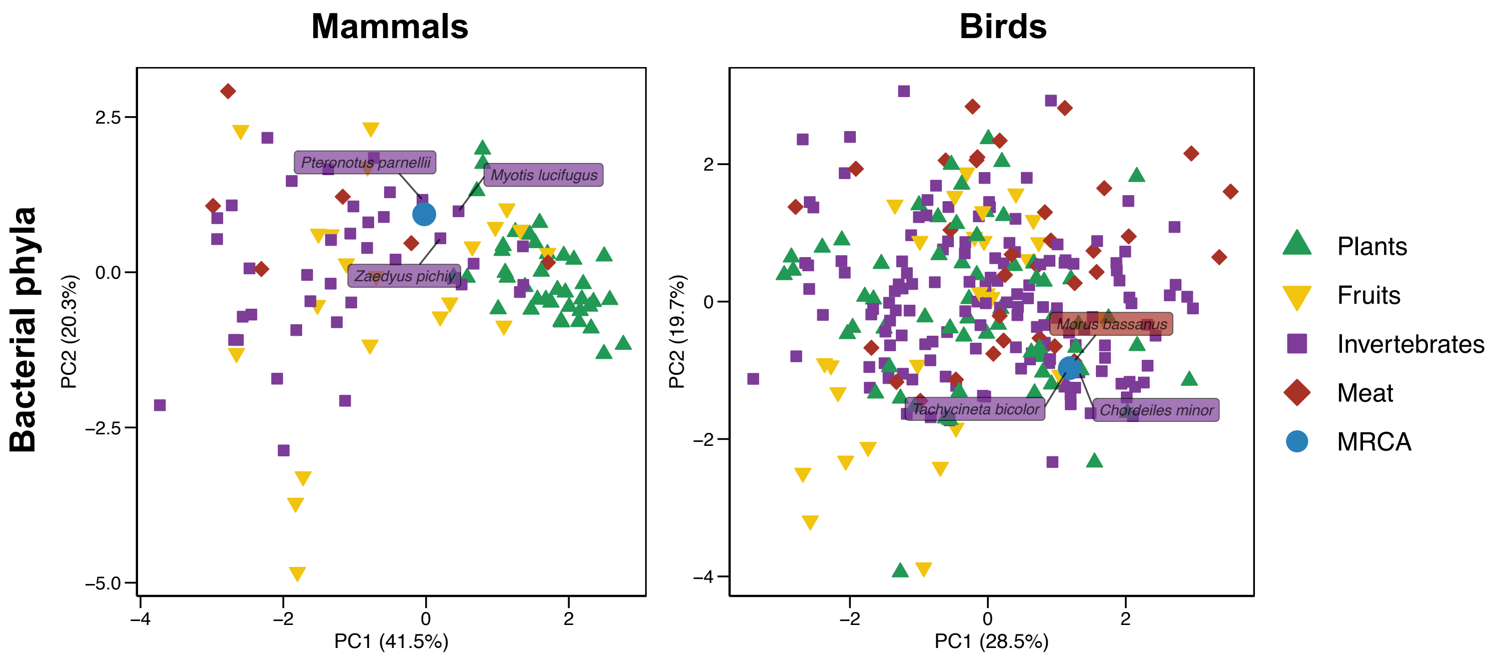

**Supplementary Figure 9: Ancestral gut microbiota compositions estimated for mammal and bird orders are similar when using joint or separate inferences.**

Estimations of the relative abundances of the 7 most abundant bacterial phyla (left) or the 14 most abundant bacterial orders (right) in the gut microbiota of the most recent common ancestor of the main mammal or bird orders using the joint inference (all mammals or all birds) or a separate inference (for each mammal or bird order). For the joint inferences, we inferred the ancestral abundances at each node of the phylogenetic trees using generalized least squares (following Martins & Hansen 1997, Cunningham et al. 1998, Clavel et al. 2018).
Results were qualitatively similar when using only the 5 or 9 most abundant phyla or orders respectively.

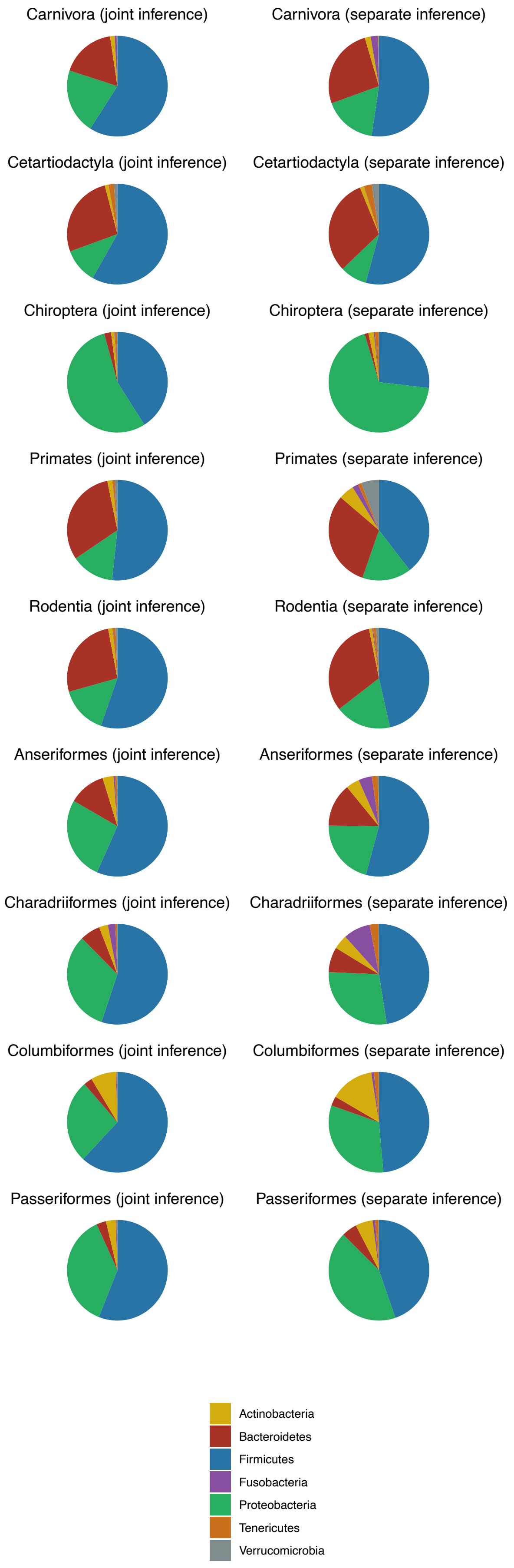
Carnivora (joint inference)
Carnivora (separate inference)
Cetartiodactyla (joint inference)
Cetartiodactyla (separate inference)
Chiroptera (joint inference)
Chiroptera (separate inference)
Primates (joint inference)
Primates (separate inference)
Rodentia (joint inference)
Rodentia (separate inference)
Anseriformes (joint inference)
Anseriformes (separate inference)
Charadriiformes (joint inference)
Charadriiformes (separate inference)
Columbiformes (joint inference)
Columbiformes (separate inference)
Passeriformes (joint inference)
Passeriformes (separate inference)
Actinobacteria
Bacteroidetes
Firmicutes
Fusobacteria
Proteobacteria
Tenericutes
Verrucomicrobia

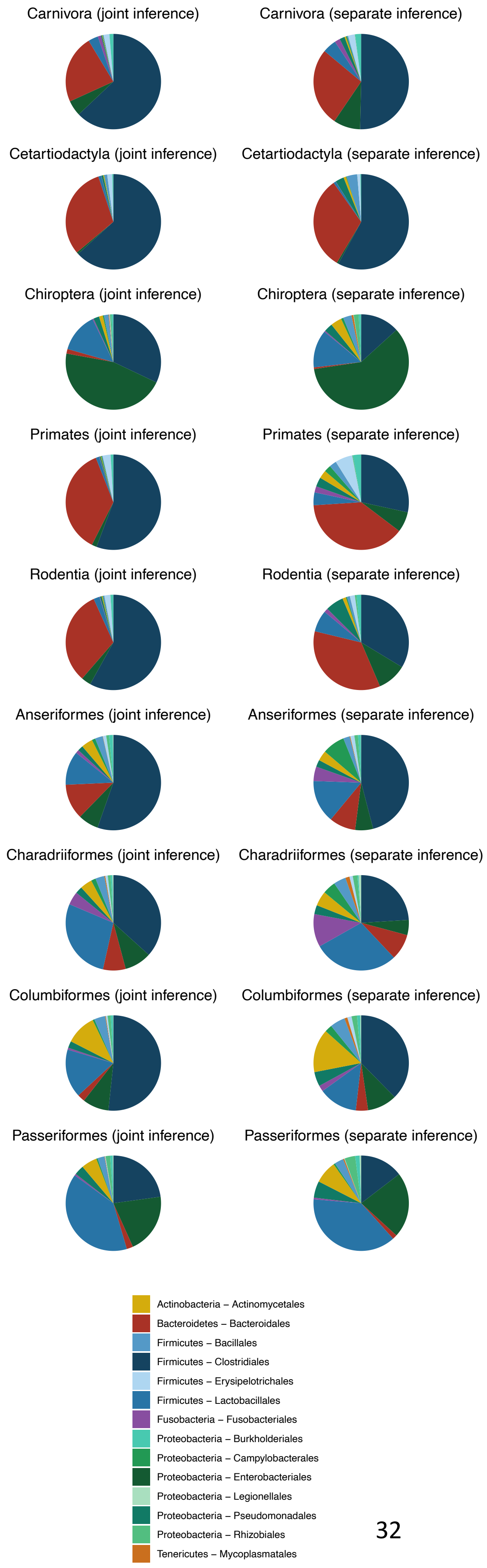
Carnivora (joint inference)
Carnivora (separate inference)
Cetartiodactyla (joint inference)
Cetartiodactyla (separate inference)
Chiroptera (joint inference)
Chiroptera (separate inference)
Primates (joint inference)
Primates (separate inference)
Rodentia (joint inference)
Rodentia (separate inference)
Anseriformes (joint inference)
Anseriformes (separate inference)
Charadriiformes (joint inference)
Charadriiformes (separate inference)
Columbiformes (joint inference)
Columbiformes (separate inference)
Passeriformes (joint inference)
Passeriformes (separate inference)
Actinobacteria – Actinomycetales
Bacteroidetes – Bacteroidales
Firmicutes – Bacillales
Firmicutes – Clostridiales
Firmicutes – Erysipelotrichales
Firmicutes – Lactobacillales
Fusobacteria – Fusobacteriales
Proteobacteria – Burkholderiales
Proteobacteria – Campylobacterales
Proteobacteria – Enterobacteriales
Proteobacteria – Legionellales
Proteobacteria – Pseudomonadales
Proteobacteria – Rhizobiales
Tenericutes – Mycoplasmatales

**Supplementary Figure 10: Correlations between the covariances between bacterial orders estimated using a joint inference (all mammals of all birds) or separate inferences performed for the main mammal or bird orders.**

For information, covariances estimated using separate inferences were often estimated as not significant when the number of host species was low, as predicted from simulations.
Results were qualitatively similar at the phylum level or when using only the 9 most abundant orders.

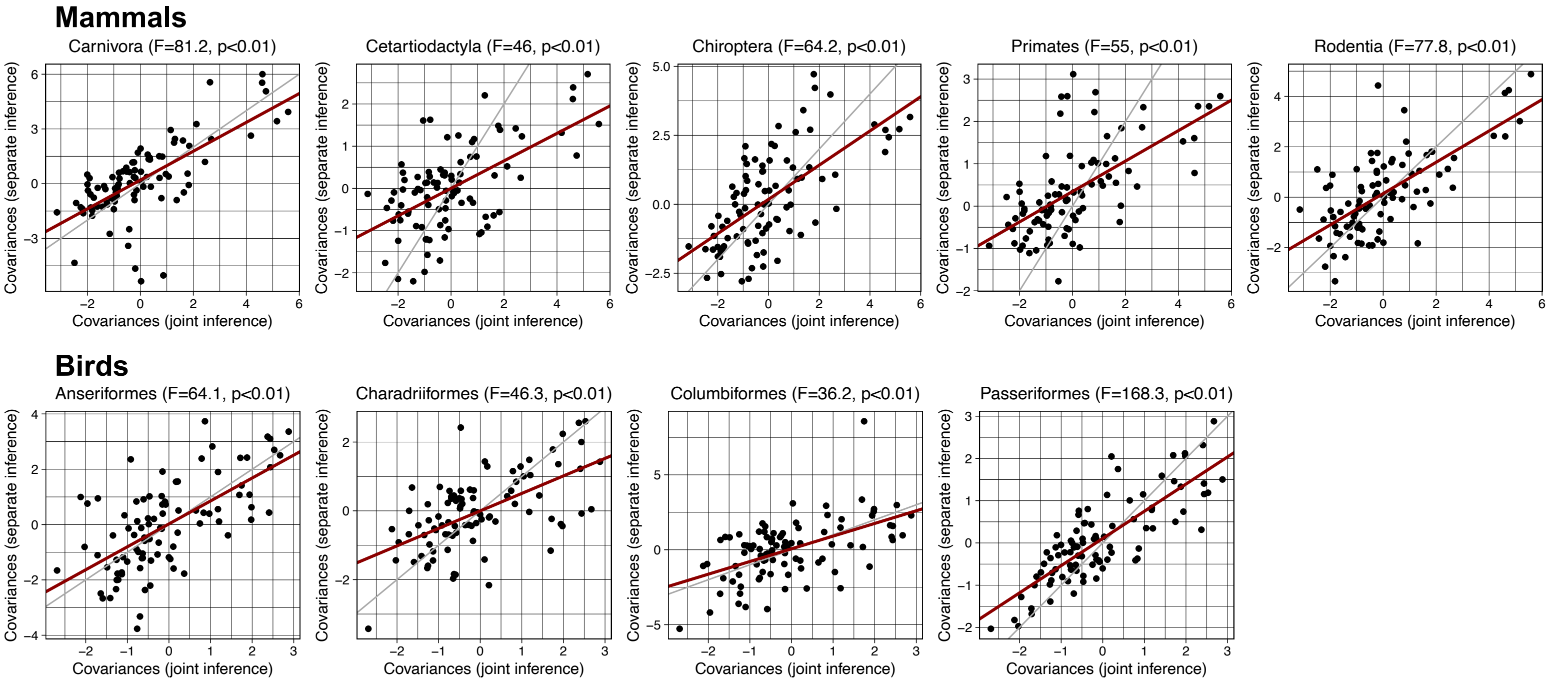

**Supplementary Figure 11: Ancestral gut microbiota composition of mammals and birds in terms of the two main subsets of bacterial orders identified in our study.**

Estimations of the relative abundances of the 2 main subsets of bacterial orders in the gut microbiota of the ancestors of mammals (a) or birds (b). Subsets are based on clustering analyses performed on the variance-covariance matrix (Figure 6).
Extant abundances are indicated with bar charts on the right.
We inferred the ancestral abundances at each node of the phylogenetic trees using generalized least squares (following Martins & Hansen 1997, Cunningham et al. 1998, Clavel et al. 2018).

**(a) Mammals**

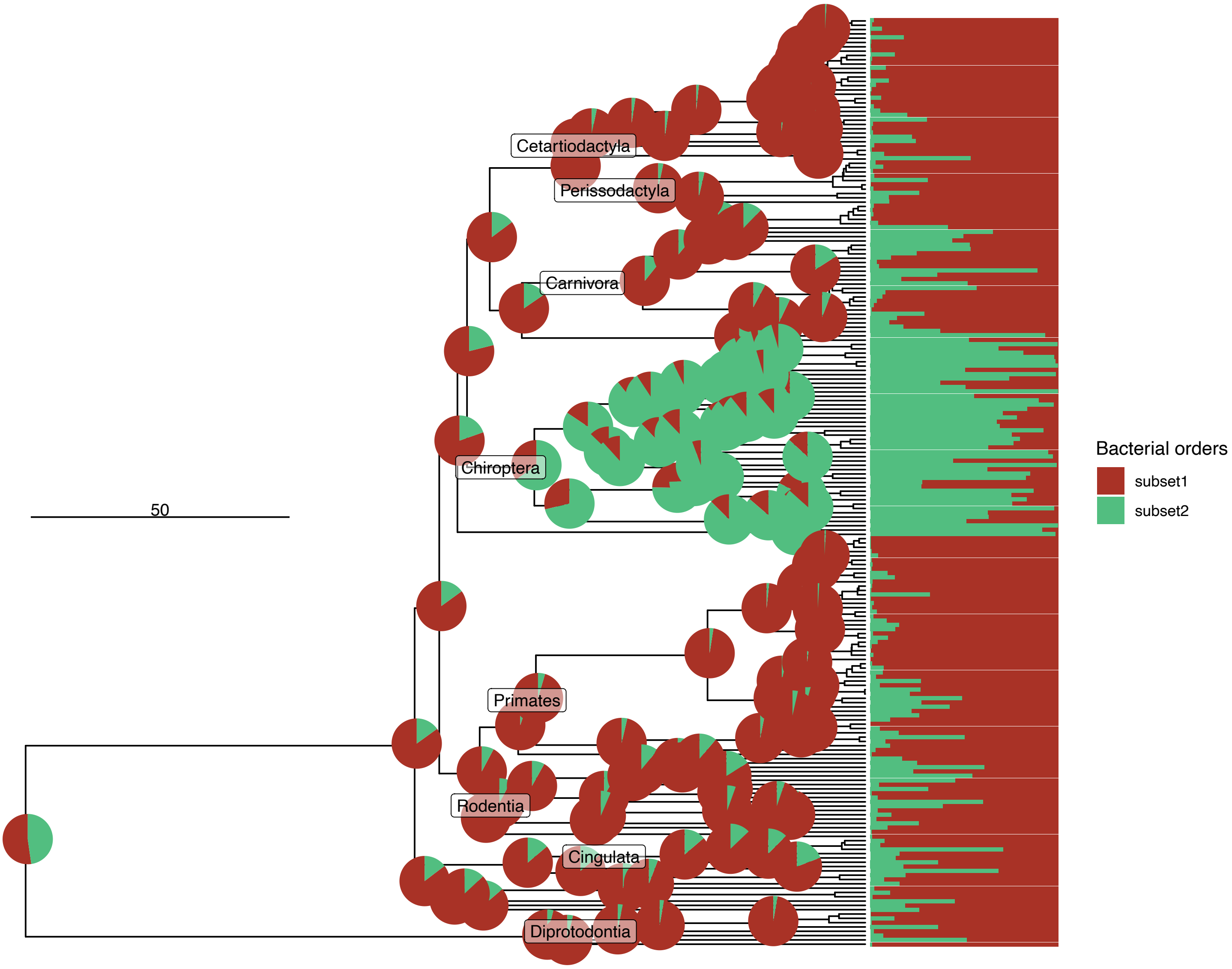

#### (b) Birds

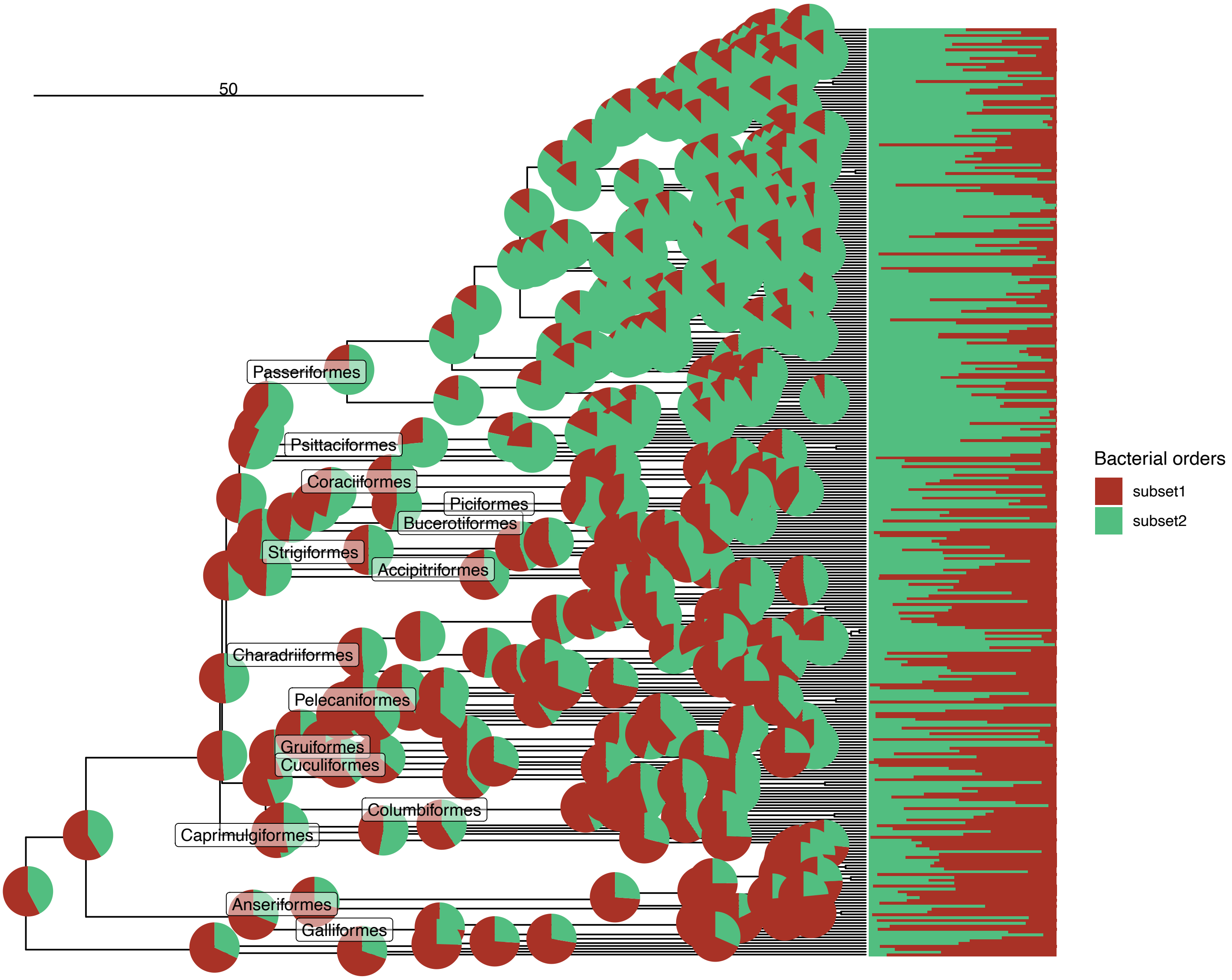
50
Passeriformes
Psittaciformes
Coraciiformes
Piciformes
Bucerotiformes
Strigiformes
Accipitriformes
Charadriiformes
Pelecaniformes
Gruiformes
Cuculiformes
Columbiformes
Caprimulgiformes
Anseriformes
Galliformes
Bacterial orders
subset1
subset2

**Supplementary Figure 12: Posterior predictive checks suggest that our model can generate realistic gut microbiota for extant mammals and birds.**

To assess whether our model for the evolution of the gut microbiota of mammals and birds can generate realistic microbiota, we simulated the process of microbiota evolution on the mammal or bird phylogenies using the parameters estimated for mammals and birds (logZ0, R, and $\lambda$) and we compared the simulated microbiota compositions to the empirical microbiota compositions of the extant mammal or bird species.
Inferences and the associated simulations were performed at the bacterial order level (using the 9 or 14 most abundant orders) or at the bacterial phylum level (using the 5 or 7 most abundant phyla). For each inference, 20 independent simulations were performed.

**(a)** Principal component analyses (PCA) comparing the composition of the simulated microbiota (in blue) to the composition of the empirical microbiota (in red).

**(b)** Distributions of the Bray-Curtis dissimilarities of the empirical microbiota of mammal and bird extant species (in red) or dissimilarities of the simulated microbiota of the extant species (in blue). For illustration purposes, the distributions for only 10 simulations are represented here.

We observed that empirical microbiota often did not occupy the full space of simulated microbiota on PCA plots, meaning that our model predicts a wider range of possible microbiota compositions than is observed in reality. Similarly, average Bray-Curtis dissimilarities between extant species were sometimes higher in simulated than in empirical microbiota, which presented less variability.

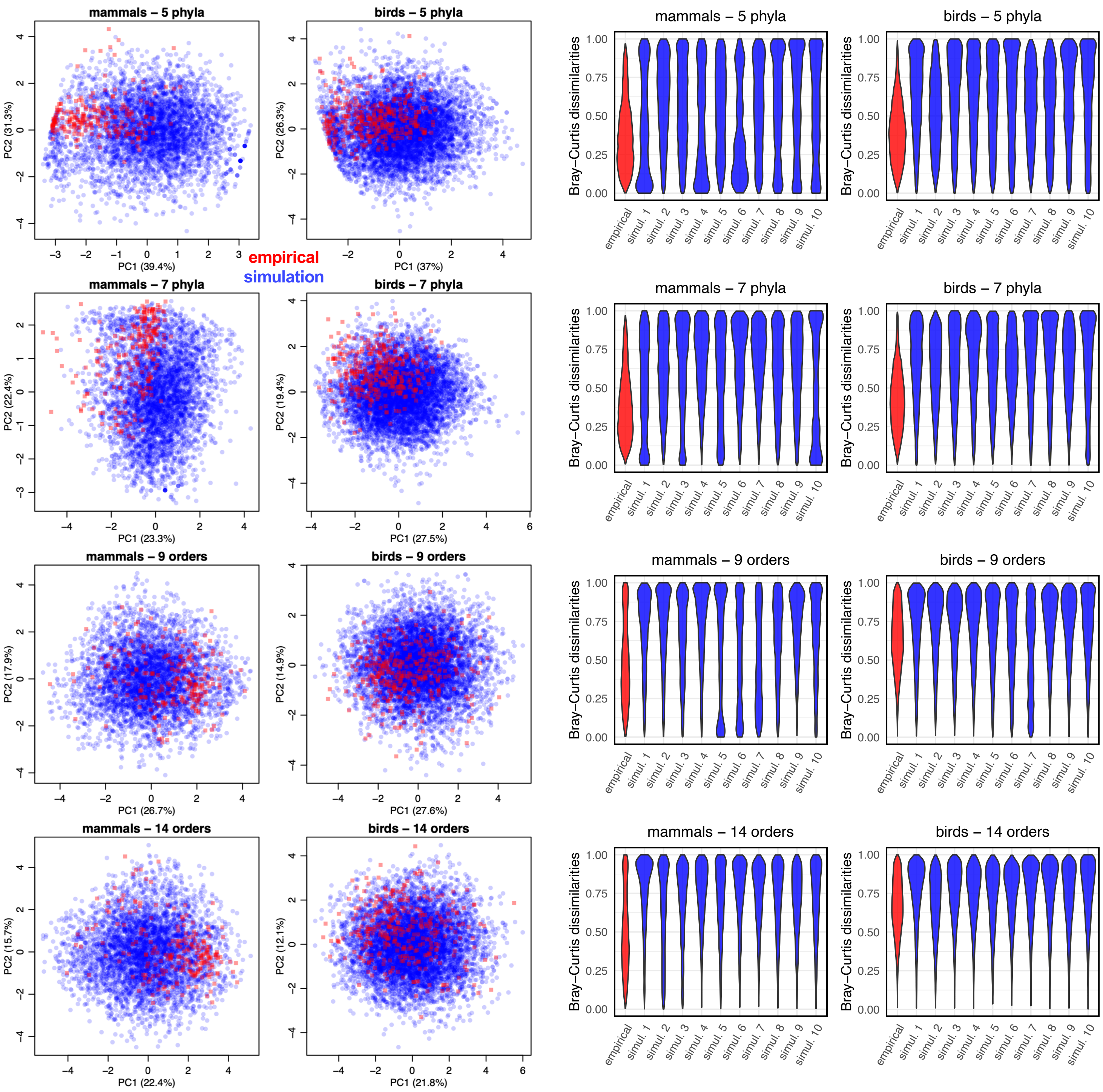
mammals – 5 phyla
birds – 5 phyla
mammals – 7 phyla
birds – 7 phyla
mammals – 9 orders
birds – 9 orders
mammals – 14 orders
birds – 14 orders
empirical
simulation
PC1 (39.4%)
PC2 (31.3%)
PC1 (37%)
PC2 (26.3%)
PC1 (23.3%)
PC2 (22.4%)
PC1 (27.5%)
PC2 (19.4%)
PC1 (26.7%)
PC2 (17.9%)
PC1 (27.6%)
PC2 (14.9%)
PC1 (22.4%)
PC2 (15.7%)
PC1 (21.8%)
PC2 (12.1%)
Bray–Curtis dissimilarities
empirical
simul. 1
simul. 2
simul. 3
simul. 4
simul. 5
simul. 6
simul. 7
simul. 8
simul. 9
simul. 10

**Supplementary Figure 13: Parameters used for the simulations testing the model are similar to the parameter values estimated on the empirical data.**

Distributions of the R, logZ0, and logY parameter values of the empirical inferences (in blue; for all mammals and birds, the main mammal and bird orders, and for different bacterial taxonomical levels) and of the simulations (simulated values in green and inferred values in red).

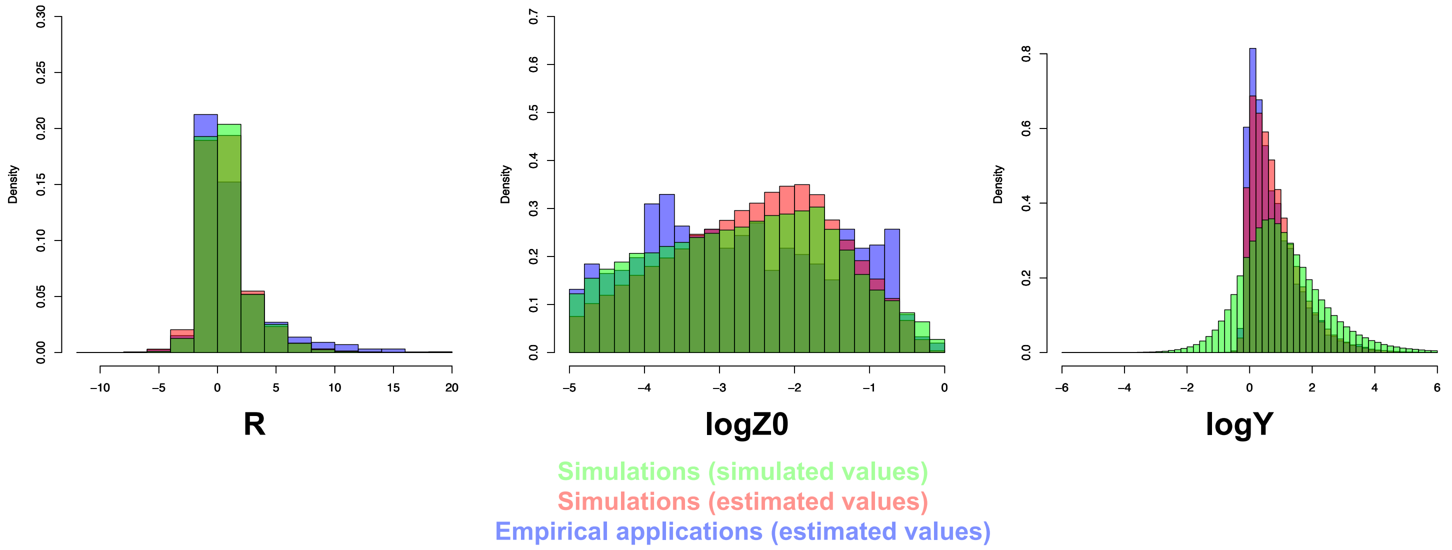

**Supplementary Figure 14: Parameter estimations are similar when randomly picking one sample per host species instead of averaging several samples per host species.**

For each type of parameter (λ, logZ0, the variances of R or the covariances of R), we represented the estimated values obtained when randomly picking one sample per host species (y-axis) as a function of the values obtained when averaging several samples per host species (x-axis). We only represented inferences performed for all mammals and birds at the bacterial order level. The black line represents the y=x axis.

We noticed that λ values tend to be slightly lower when taking only one individual per species because averaging several samples tend to reduce the within-species variability and thus leads to a higher λ value. Similarly, higher variances were often observed when taking only one individual per species, as it resulted in an increased present-day variability in the dataset.

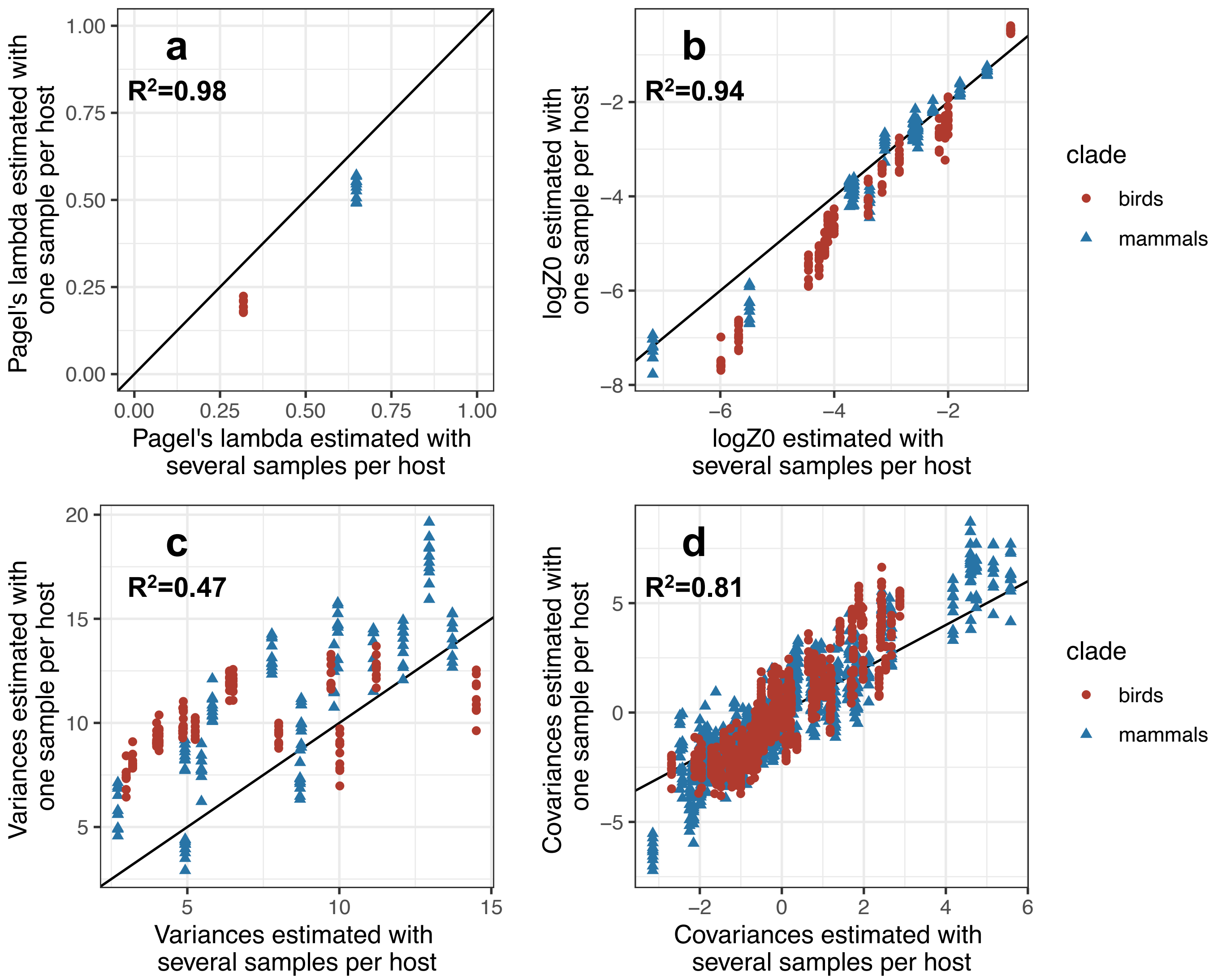

**Supplementary Figure 15: Average gut microbiota composition of mammal and bird species at the bacterial phyla level.**

Relative abundances of the 7 most abundant bacterial phyla in mammals (left) and birds (right).
These bacterial phyla represent 95% and 96% of the gut microbiota of mammals and birds respectively.

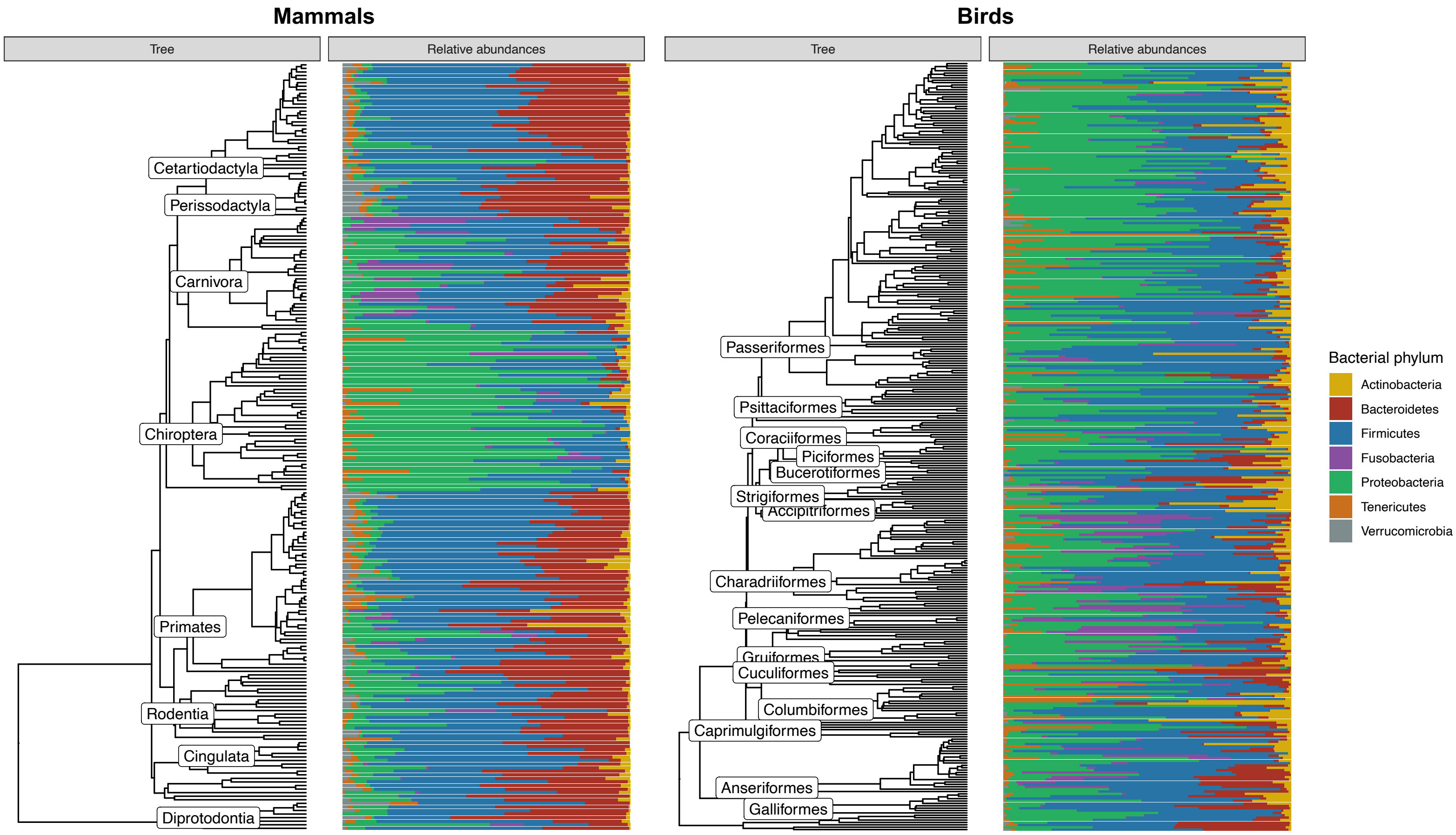

**Supplementary Figure 16: Ancestral reconstructions are robust to the minimal abundance of bacterial taxa (threshold of non-detection).**

We replicated the inferences with a minimal abundance of 0.01% for each bacterial order (instead of 0.001%) and reported its effect on the estimations of bacterial variances (a) and ancestral microbiota compositions (b).

**(a) Estimated variances of rare bacterial taxa are higher when the threshold of minimal abundances is set to lower values:**

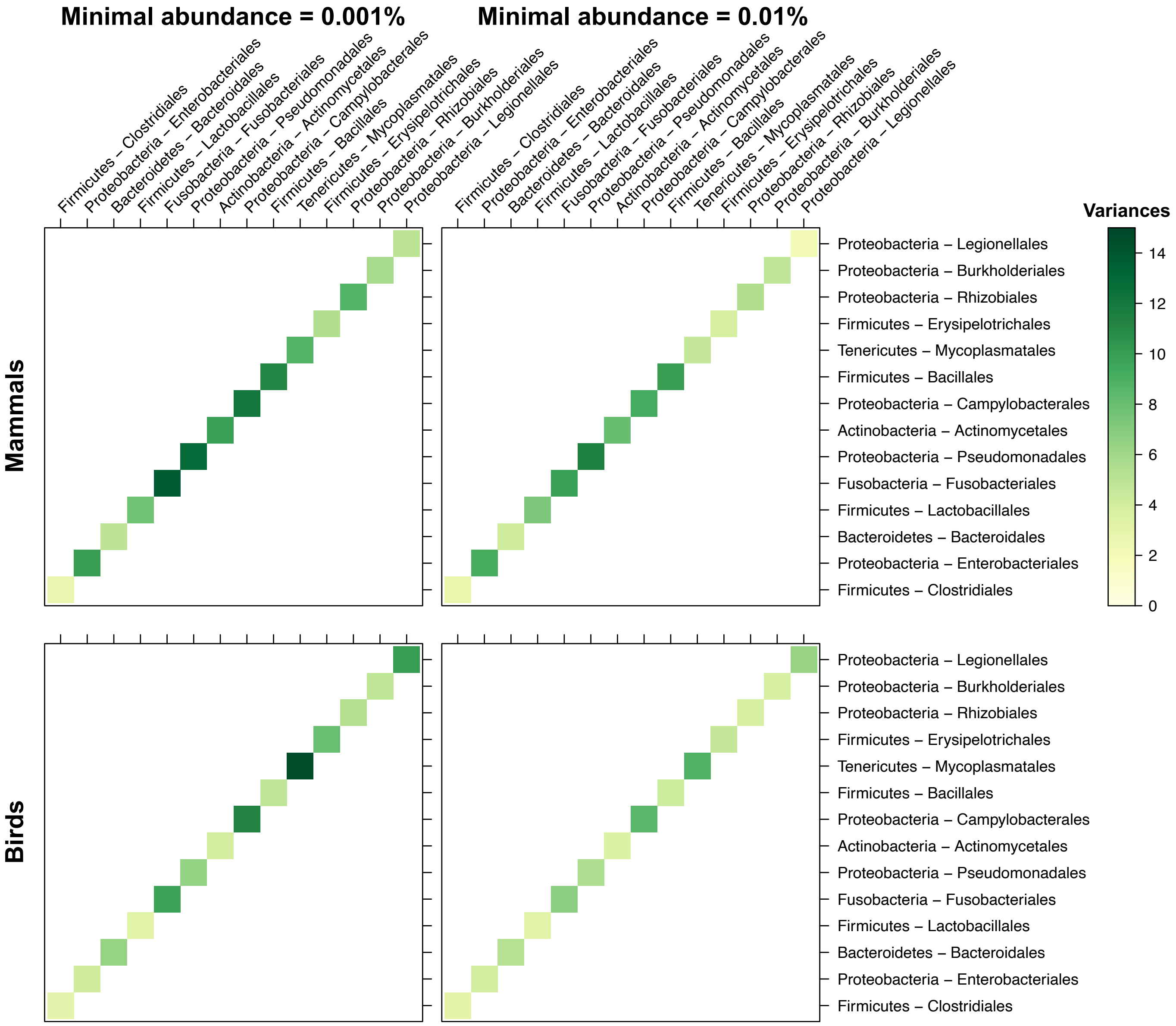

**(b) Ancestral microbiota compositions are not affected by the threshold of minimal abundances:**

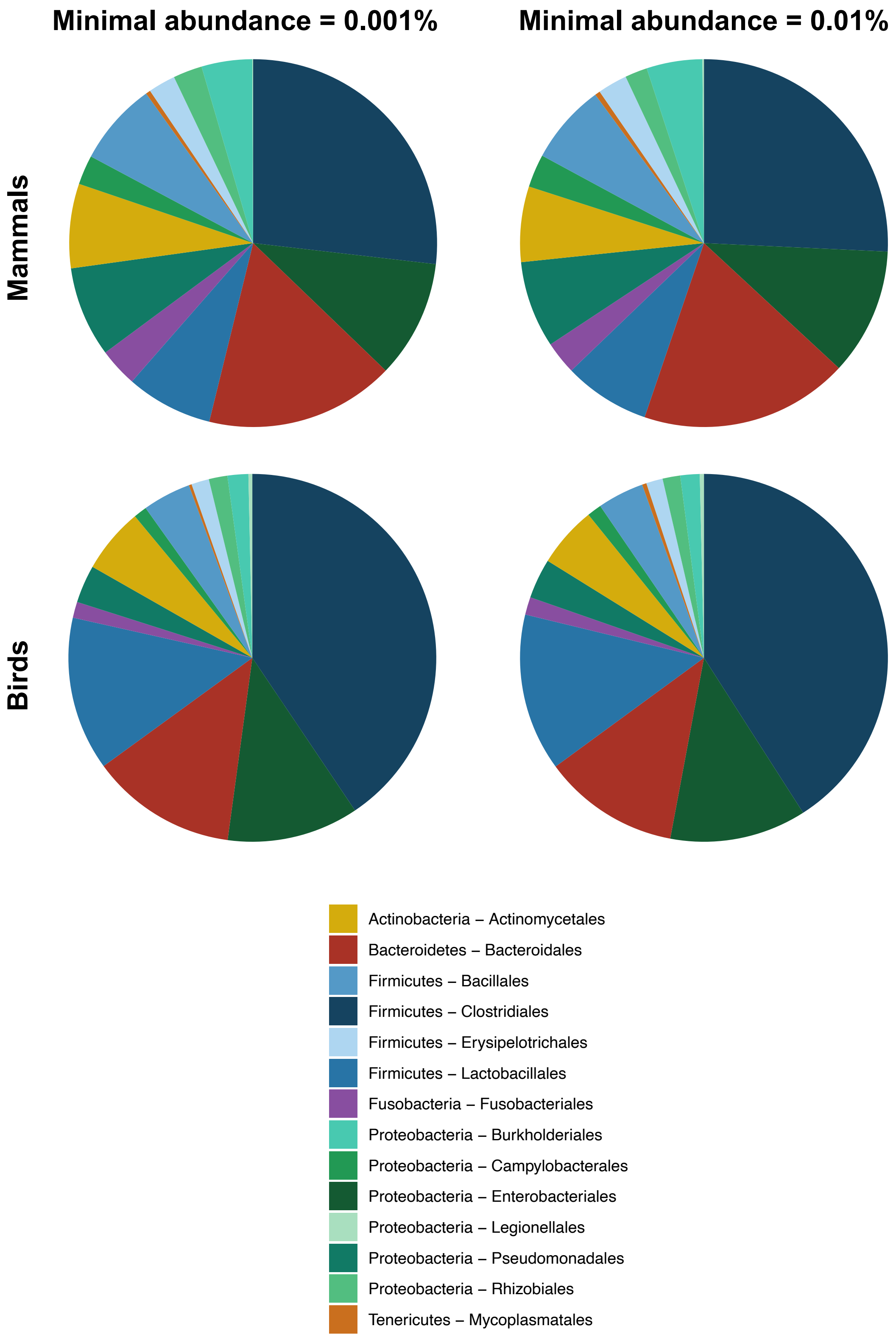

#### Simulations:

**Supplementary Figure 17: Simulations validate the correct estimation of the ancestral microbiota composition (logX0 values):**

For each number of host species (n) and each number of bacterial taxa (p), we represented the estimated ancestral abundances (logX0) as a function of the simulated ancestral abundances. For each combination of n and p, we performed 500 simulations (with $\lambda$=0, 0.25, 0.5, 0.75, or 1).
The black line corresponds to the y=x axis, while the purple lines correspond to the fitted mixed linear model between estimated and simulated logX0 values (with each simulation as a random effect). We fitted a different linear model per simulated $\lambda$ value.

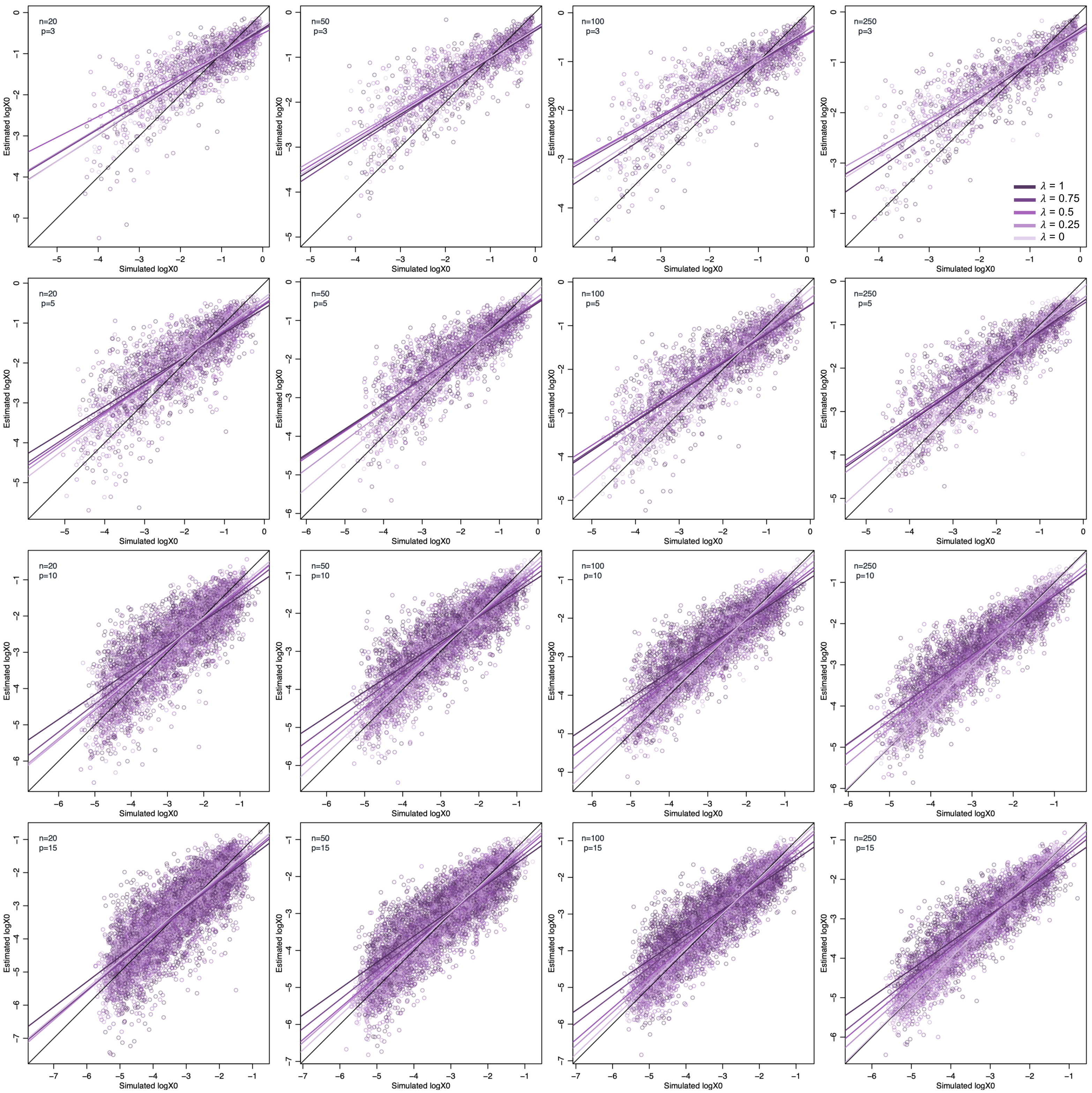
n=20
p=3
n=50
p=3
n=100
p=3
n=250
p=3
n=20
p=5
n=50
p=5
n=100
p=5
n=250
p=5
n=20
p=10
n=50
p=10
n=100
p=10
n=250
p=10
n=20
p=15
n=50
p=15
n=100
p=15
n=250
p=15
Estimated logX0
Simulated logX0
$\lambda$ = 1
$\lambda$ = 0.75
$\lambda$ = 0.5
$\lambda$ = 0.25
$\lambda$ = 0

**Supplementary Figure 18: Simulations validate the correct estimation of the variance and covariance values (R):**

For each number of host species (n) and each number of bacterial taxa (p), we represented the estimated R values as a function of the simulated R values. We evaluated the significance of the R values, by considering a R value as positive (resp. negative) if the whole 95% credible interval is positive (resp. negative).
We represented significant R values using grey rounds, and non-significant R values using brown crosses. For each combination of n and p, we performed 500 simulations (with $\lambda$=0, 0.25, 0.5, 0.75, or 1).
The black line corresponds to the y=x axis, while the orange line corresponds to the fitted mixed linear model between estimated and simulated R values (with each simulation as a random effect). The blue rectangle corresponds to positive R values estimated as positive and the red rectangle to negative R values estimated as negative.

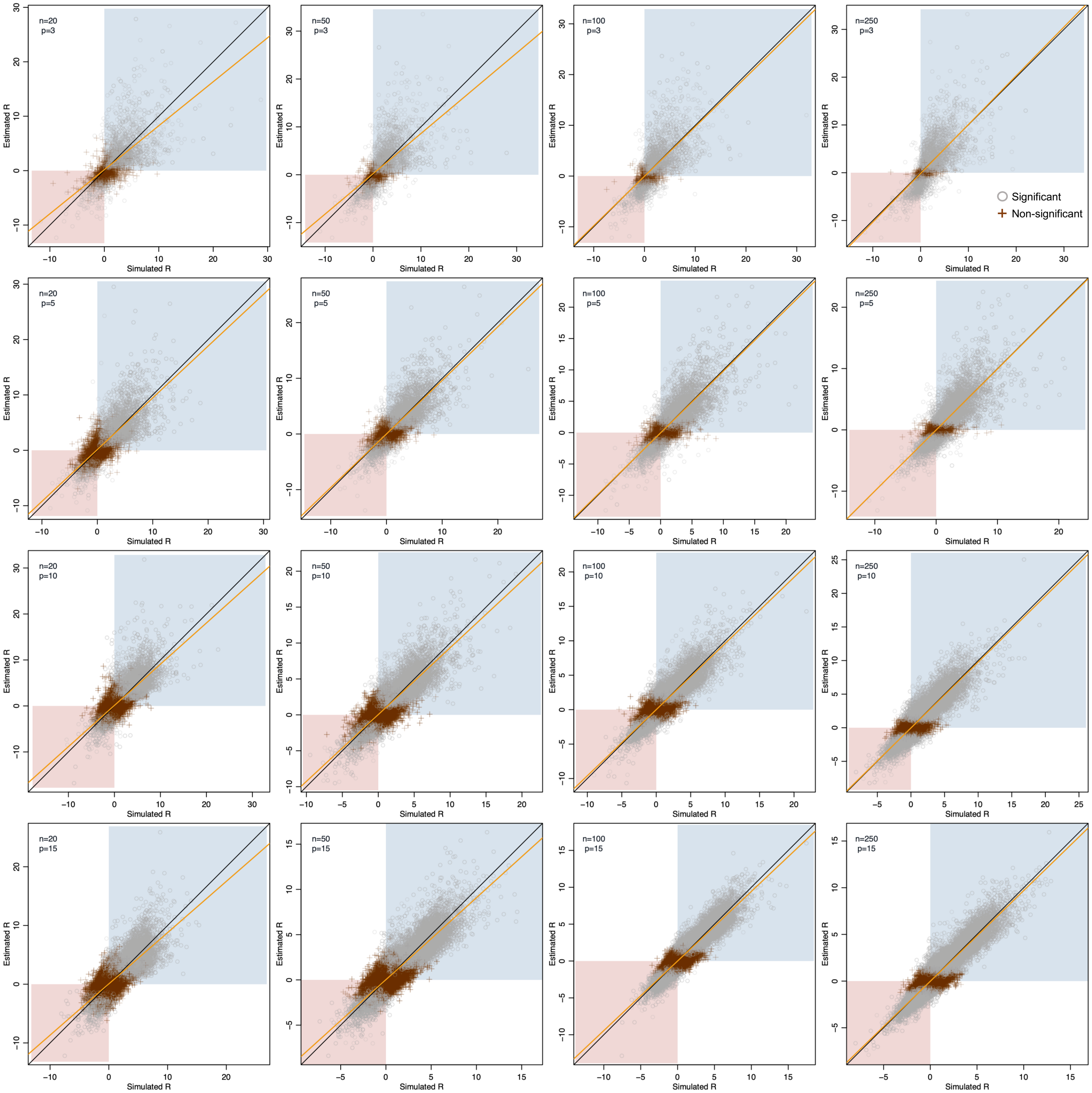
n=20
p=3
n=50
p=3
n=100
p=3
n=250
p=3
Significant
Non-significant
n=20
p=5
n=50
p=5
n=100
p=5
n=250
p=5
n=20
p=10
n=50
p=10
n=100
p=10
n=250
p=10
n=20
p=15
n=50
p=15
n=100
p=15
n=250
p=15
Estimated R
Simulated R

**Supplementary Figure 19: Simulations validate the correct estimation of the level of phylosymbiosis (λ):**

For each number of host species (n) and each number of bacterial taxa (p), we represented the estimated λ values as a function of the simulated λ values. For each simulation, we evaluated the significance of phylosymbiosis using Bayes factors (BF), by considering the phylosymbiosis to be not significant when BF≤1. We represented significant λ using rounds, and non-significant λ using crosses. For each combination of n, p, and λ values, we performed 100 simulations. The black line corresponds to the y=x axis.

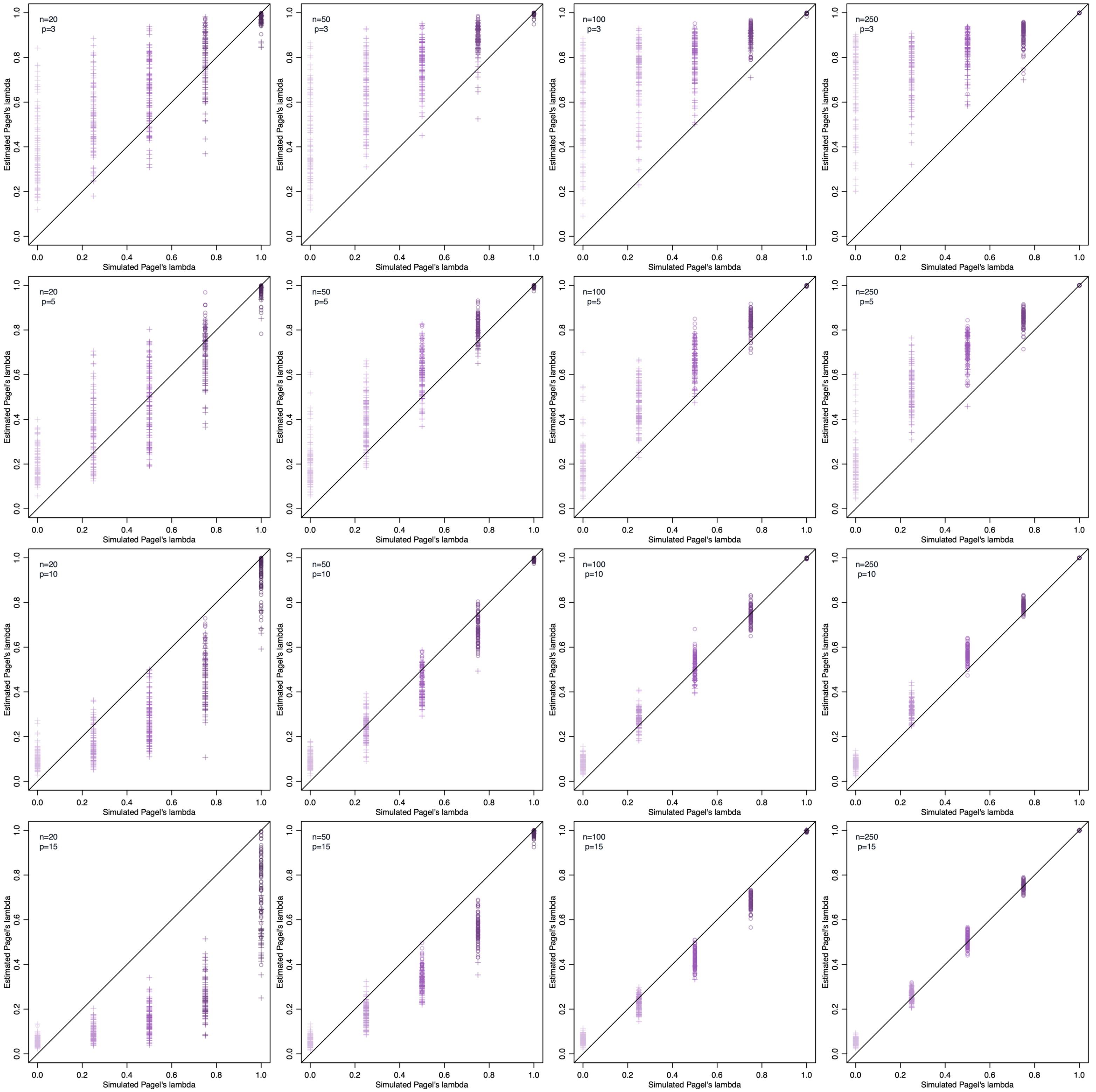

**Supplementary Figure 20: Simulations indicate that increases in the total bacterial abundances (logY>0) are correctly estimated, while decreases in the total bacterial abundances (logY<0) are not.**

For each number of host species (n) and each number of bacterial taxa (p), we represented the estimated total bacterial abundances (logY) as a function of the simulated total bacterial abundances in each extant host species. For each combination of n and p, we performed 500 simulations (with $\lambda$=0, 0.25, 0.5, 0.75, or 1). The black line corresponds to the y=x axis, while the purple lines correspond to the fitted mixed linear model between estimated and simulated logY values (with each simulation as a random effect). We fitted a different linear model per simulated $\lambda$ value.

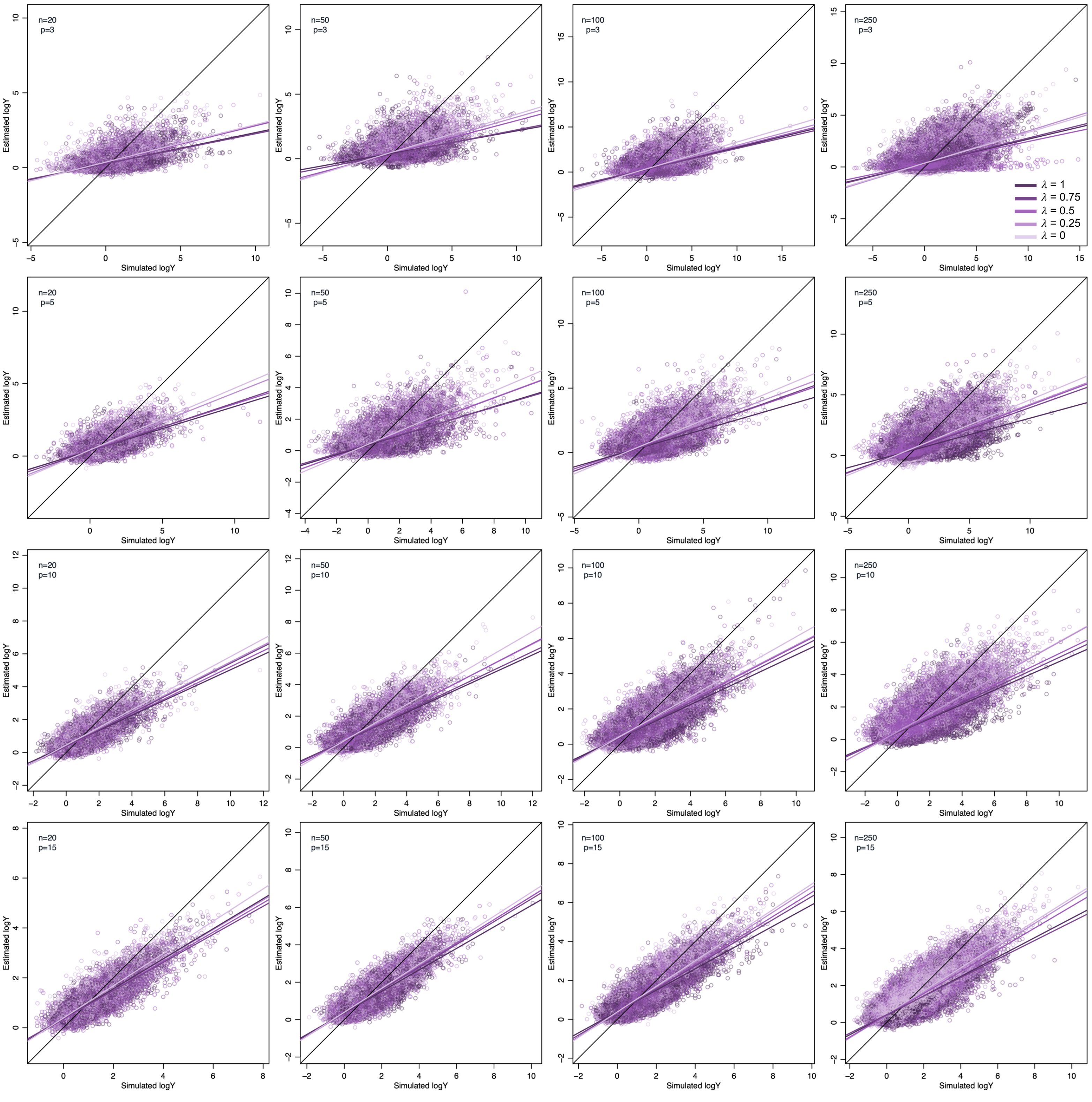
n=20
p=3
n=50
p=3
n=100
p=3
n=250
p=3
n=20
p=5
n=50
p=5
n=100
p=5
n=250
p=5
n=20
p=10
n=50
p=10
n=100
p=10
n=250
p=10
n=20
p=15
n=50
p=15
n=100
p=15
n=250
p=15
Estimated logY
Simulated logY
$\lambda$ = 1
$\lambda$ = 0.75
$\lambda$ = 0.5
$\lambda$ = 0.25
$\lambda$ = 0

#### Simulations on empirical phylogenies:

**Supplementary Figure 21: Simulations on empirical trees validate the correct estimation of the level of phylosymbiosis (λ):**

For each category of simulations (mammals/birds and phyla/orders), we represented the estimated λ values as a function of the simulated λ values. For each simulation, we evaluated the significance of phylosymbiosis using Bayes factors (BF), by considering the phylosymbiosis to be not significant when BF≤1. We represented significant λ using rounds, and non-significant λ using crosses. The black line corresponds to the y=x axis.

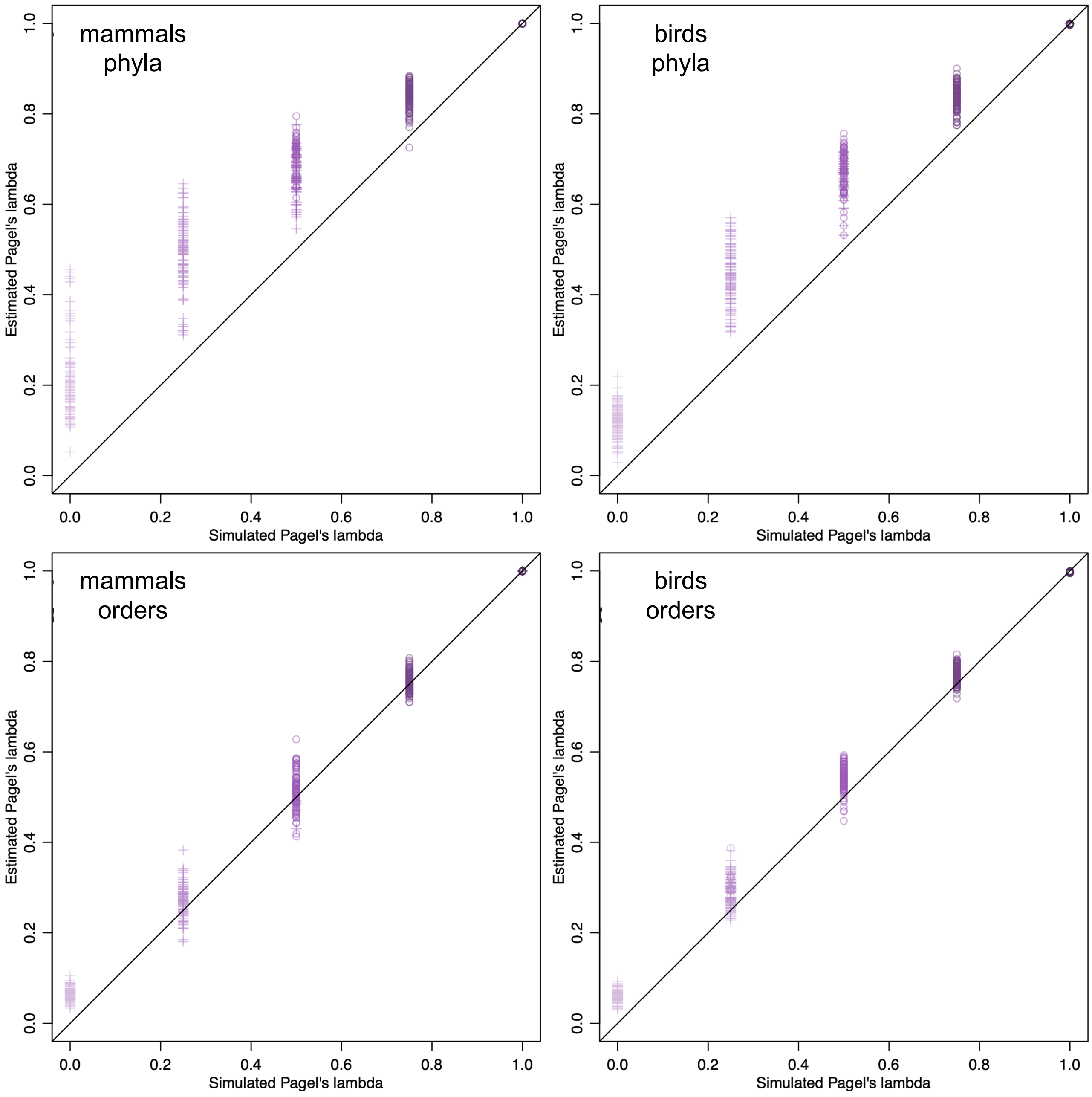

**Supplementary Figure 22: Simulations on empirical trees validate the correct estimation of the ancestral microbiota composition (logX0 values):**

For each category of simulations (mammals/birds and phyla/orders), we represented the estimated ancestral abundances (logX0) as a function of the simulated ancestral abundances.

The black line corresponds to the y=x axis, while the purple lines correspond to the fitted mixed linear model between estimated and simulated logX0 values (with each simulation as a random effect). We fitted a different linear model per simulated λ value.

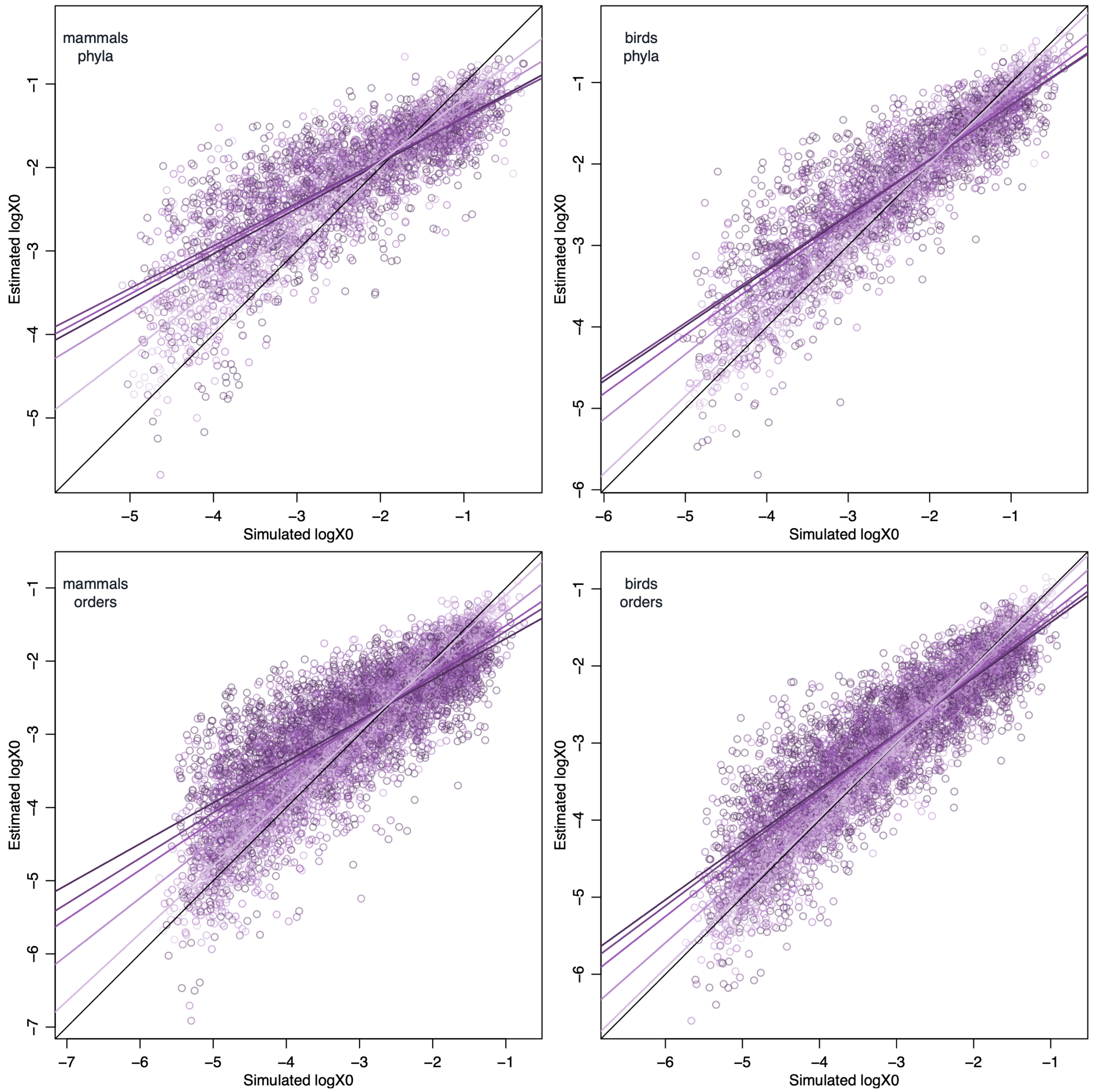

**Supplementary Figure 23: Simulations on empirical trees validate the correct estimation of the variance and covariance values (R):**

For each category of simulations (mammals/birds and phyla/orders), we represented the estimated R values as a function of the simulated R values. We evaluated the significance of the R values, by considering a R value as positive (resp. negative) if the whole 95% credible interval is positive (resp. negative).
We represented significant R values using grey rounds, and non-significant R values using brown crosses.
The black line corresponds to the y=x axis, while the orange line corresponds to the fitted mixed linear model between estimated and simulated R values (with each simulation as a random effect). The blue rectangle corresponds to positive R values estimated as positive and the red rectangle to negative R values estimated as negative.

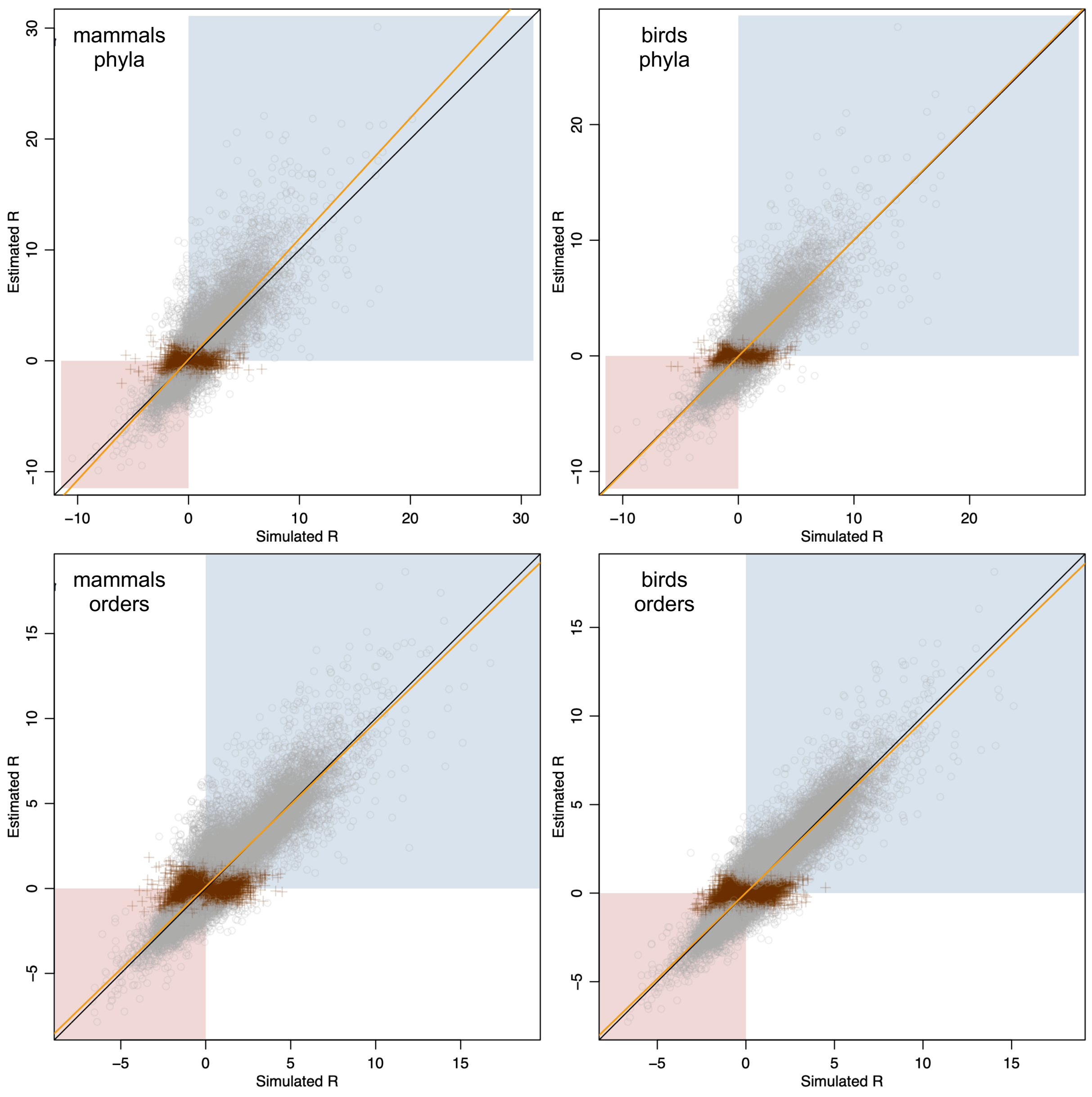

**Supplementary Figure 24: Different experimental conditions for characterizing gut microbiota are not responsible of the phylosymbiosis in the gut microbiota of mammals and birds.**

**(a)** For both mammals and birds, we compared the estimated level of phylosymbiosis (mean λ value in orange) to levels of phylosymbiosis (λ values) estimated when shuffling the species having their microbiota characterized in the same study (i.e. with the same experimental conditions).
Boxplots indicate the median surrounded by the first and third quartiles, and whiskers extend to the extreme values but no further than 1.5 of the inter-quartile range.
Randomizations were performed on gut microbiota at the phylum level using the 7 most abundant phyla (top row) or at the order level using the 14 most abundant phyla (bottom row). Results were qualitatively similar when using only the 5 or 9 most abundant phyla or orders respectively.
**(b-c)** Each separate study tends to characterize closely related species in mammals (b), but not in birds (c). We indicated on the phylogenetic trees the study ID where each species has been described.

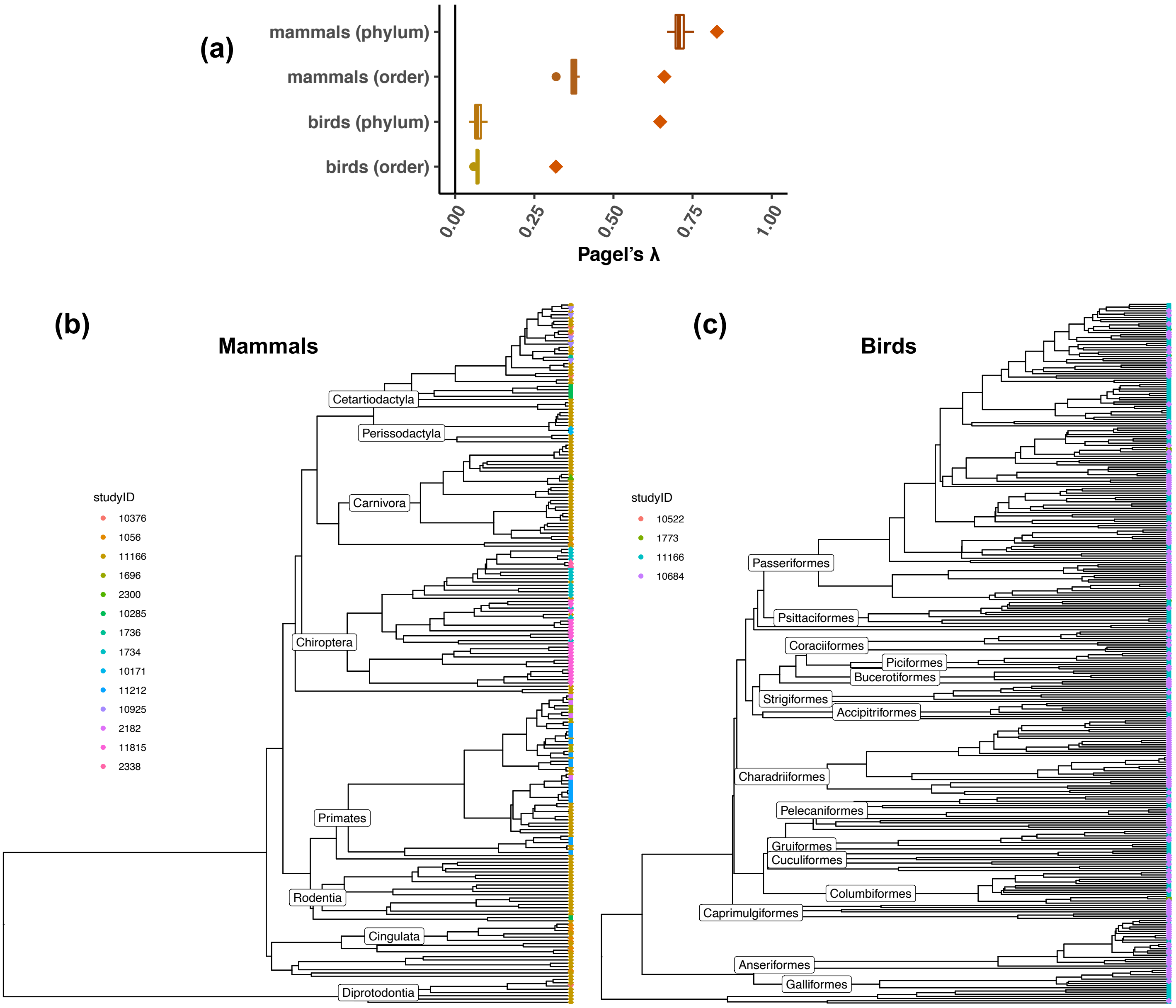
